# A Large-Scale Binding and Functional Map of Human RNA Binding Proteins

**DOI:** 10.1101/179648

**Authors:** Eric L Van Nostrand, Peter Freese, Gabriel A Pratt, Xiaofeng Wang, Xintao Wei, Rui Xiao, Steven M Blue, Jia-Yu Chen, Neal A.L. Cody, Daniel Dominguez, Sara Olson, Balaji Sundararaman, Lijun Zhan, Cassandra Bazile, Louis Philip Benoit Bouvrette, Julie Bergalet, Michael O Duff, Keri E. Garcia, Chelsea Gelboin-Burkhart, Myles Hochman, Nicole J Lambert, Hairi Li, Thai B Nguyen, Tsultrim Palden, Ines Rabano, Shashank Sathe, Rebecca Stanton, Amanda Su, Ruth Wang, Brian A. Yee, Bing Zhou, Ashley L Louie, Stefan Aigner, Xiang-dong Fu, Eric Lécuyer, Christopher B. Burge, Brenton R. Graveley, Gene W. Yeo

**Affiliations:** Department of Cellular and Molecular Medicine, University of California at San Diego, La Jolla, CA; Institute for Genomic Medicine, University of California at San Diego, La Jolla, CA; Program in Computational and Systems Biology, MIT Cambridge MA; Bioinformatics and Systems Biology Graduate Program, University of California San Diego, La Jolla, CA; Institut de Recherches Cliniques de Montréal (IRCM), Montreal, Canada; Department of Genetics and Genome Sciences, Institute for Systems Genomics, UConn Health, Farmington, CT; Department of Biology, MIT, Cambridge MA; Department of Biological Engineering, MIT, Cambridge MA; Department of Biochemistry and Molecular Medicine, Université de Montréal, Montreal, Canada; Division of Experimental Medicine, McGill University, Montreal, Canada

## Abstract

Genomes encompass all the information necessary to specify the development and function of an organism. In addition to genes, genomes also contain a myriad of functional elements that control various steps in gene expression. A major class of these elements function only when transcribed into RNA as they serve as the binding sites for RNA binding proteins (RBPs), which act to control post-transcriptional processes including splicing, cleavage and polyadenylation, RNA editing, RNA localization, stability, and translation. Despite the importance of these functional RNA elements encoded in the genome, they have been much less studied than genes and DNA elements. Here, we describe the mapping and characterization of RNA elements recognized by a large collection of human RBPs in K562 and HepG2 cells. These data expand the catalog of functional elements encoded in the human genome by addition of a large set of elements that function at the RNA level through interaction with RBPs.

**Highlights:**

- 223 eCLIP datasets for 150 RBPs reveal a wide variety of *in vivo* RNA target classes.
- 472 knockdown/RNA-seq profiles of 263 RBPs reveal factor-responsive targets and integration with eCLIP indicates RNA expression and splicing regulatory patterns.
- 78 RNA Bind-N-Seq profiles of *in vitro* binding motifs reveal links between *in vitro* and *in vivo* binding and indicate that eCLIP peaks that contain *in vitro* motifs are more strongly associated with regulation.
- 274 maps of RBP subcellular localization by immunofluorescence indicate widespread organelle-specific RNA processing regulation.
- 63 ChIP-seq profiles of DNA association suggest broad interconnectivity between chromatin association and RNA processing.

## Introduction

RNA binding proteins (RBPs) have emerged as central players in regulating gene expression, controlling when, where, and at what rate RNAs are processed, trafficked, translated, and degraded within the cell. They represent a diverse class of proteins involved in co- and post-transcriptional gene regulation^1,2^. RBPs interact with RNA to form ribonucleoprotein complexes (RNPs), governing the maturation and fate of their target RNA substrates. Indeed, they regulate numerous aspects of gene expression including pre-mRNA splicing, cleavage and polyadenylation, RNA stability, RNA localization, RNA editing, and translation. In fact, many RBPs participate in more than one of these processes. For example, studies on the mammalian RBP Nova using a combination of crosslinking and immunoprecipitation (CLIP)-seq and functional studies revealed that Nova not only regulates alternative splicing, but also modulates poly(A) site usage^3^. Moreover, in contrast to regulation at the transcriptional level, post-transcriptional regulatory steps are often carried out in different sub-cellular compartments of the nucleus (e.g. nucleoli, nuclear speckles, paraspeckles, coiled bodies, etc.) and/or cytoplasm (e.g. P-bodies, endoplasmic reticulum, etc.) by RBPs that are localized within these compartments. These regulatory roles are essential for normal human physiology, as defects in RBP function are associated with diverse genetic and somatic disorders, such as neurodegeneration, auto-immune defects, and cancer4–10.

Traditionally, RBPs were identified by affinity purification of single proteins^11,12^. However, several groups have recently used mass spectrometry-based methods to identify hundreds of proteins bound to RNA in human and mouse cells^13–16^. Recent censuses conducted by us and others indicate that the human genome may contain between 1,072^(ref. 17)^ and 1,542^(ref. 1)^ RBP-encoding genes. This large repertoire of RBPs likely underlies the high complexity of post-transcriptional regulation, motivating concerted efforts to systematically dissect the binding properties, RNA targets and functional roles of these proteins.

The dissection of RBP-RNA regulatory networks therefore requires the integration of multiple data types, each viewing the RBP through a different lens. *In vivo* binding assays such as CLIP-seq provide a set of candidate functional elements directly bound by each RBP. Assessments of *in vitro* binding affinity help understand the mechanism driving these interactions, and (as we show) improve identification of functional associations. Functional assays that identify targets whose expression or alternative splicing is responsive to RBP perturbation can then fortify evidence of function. For example, observation of protein binding by CLIP-seq within introns flanking exons whose splicing is sensitive to RBP levels provides support for the RBP as a splicing factor and for the binding sites as splicing regulatory elements. *In vivo* interactions of RBPs with chromatin can also be assayed to provide insight into roles of some RBPs as transcription regulators and can provide evidence for co-transcriptional deposition of RBPs on target RNA substrates. The regulatory roles of RBPs are also impacted by the subcellular localization properties of RBPs and of their RNA substrates. Furthermore, these data resources comprised of multiple RBPs profiled using the same methodology and cell lines may be integrated to identify factor-specific regulatory modules, and the roles of RBPs in broader cellular regulatory networks, through integrated analyses such as those described below.

## Results

### Overview of data and processing

To work towards developing a comprehensive understanding of the binding and function of the human RBP repertoire, we used five assays to produce 1,223 replicated datasets for 356 RBPs (Fig. 1a,b, Supplementary Data 1,2). The RBPs characterized by these assays have a wide diversity of sequence and structural characteristics and participate in diverse aspects of RNA biology (Fig. 1). Functionally, these RBPs are most commonly known to play roles in the regulation of RNA splicing (98 RBPs, 28%), RNA stability and decay (71, 20%), and translation (70, 20%), with 162 RBPs (46%) having more than one function reported in the literature (Supplementary Data 1). However, 83 (23%) of the characterized RBPs have no known function in RNA biology other than being annotated as binding RNA (Fig. 1b). Although 57% of the RBPs surveyed contain well-characterized RNA binding domains [RNA recognition motif (RRM), hnRNP K homology (KH), zinc finger, RNA helicase, ribonuclease, double-stranded RNA binding (dsRBD), or pumilio/FBF domain (PUM-HD)], the remainder possess either less well studied domains or lack known RNA-binding domains altogether (Fig. 1b, Supplementary Data 1). Many RBPs had high expression in ENCODE cell lines and across a broad range of human tissues, including ribosomal proteins (RPL23A, RPS11, RPS24), translation factors (EIF4H, EEF2), and ubiquitously expressed splicing factors (HNRNPC, HNRNPA2B1) among the 10 least tissue-specific RBPs (Extended Data Fig. 1a, Supplementary Data 3). However, several other RBPs had highly tissue-specific expression exhibiting either a pattern of high expression in one or a small number of human tissues (e.g., LIN28B, IGF2BP1/3) or being differentially expressed by orders of magnitude across several human tissues (e.g., IGF2BP2 and APOBEC3C), indicating that the RNA targets and regulatory activity of these RBPs are likely modulated through cell type-specific gene expression programs.

**Figure 1.**
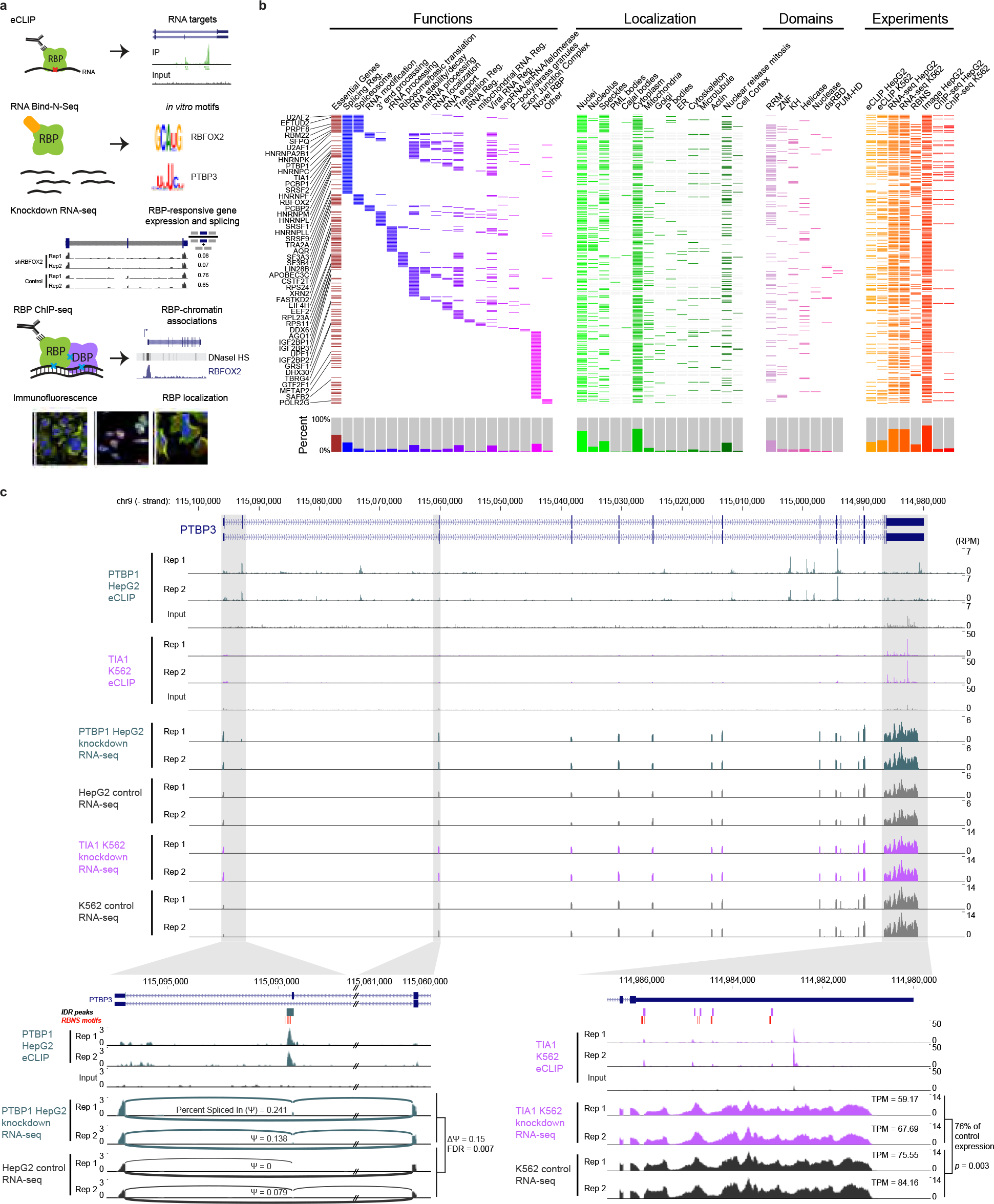
Overview of experiments and data types. (a)Models of the five assays performed to characterize RNA binding proteins (RBPs): enhanced crosslinking and immunoprecipitation (eCLIP) to identify RNA targets in HepG2 and K562 cells, RNA Bind-N-Seq (RBNS) to identify *in vitro* binding affinities, knockdown followed by RNA-seq to identify RBP-responsive genes and splicing events, ChIP-seq to identify DNA association (either direct or indirect through DNA binding proteins (DBPs)), and immunofluorescence to identify protein sub-cellular localization patterns. (b)The 352 RNA binding proteins (RBPs) profiled by at least one ENCODE experiment (orange/ red) are shown, with localization by immunofluorescence (green), essential genes from CRISPR screening (maroon), manually annotated RBP functions (blue/purple), and annotated protein domains (pink). Histograms for each category are shown on bottom, and select RBPs highlighted in this study are indicated on left. (c)Combinatorial expression and splicing regulation of *PTBP3*. Tracks indicate eCLIP and RNA-seq read density (as reads per million, RPM). (bottom left) The alternatively spliced exon 1-3 region is shown with PTBP1 eCLIP and RNA-seq, with lines indicating junction-spanning reads and percent spliced in (Ψ) is indicated. Boxes above indicate reproducible (by IDR) PTBP1 peaks in HepG2, with red boxes indicating RBNS motifs for PTB family member PTBP3 located within (or up to 50 bases upstream of) peaks. (bottom right) The 3’UTR is shown with TIA1 eCLIP and RNA-seq in K562 cells, with overall gene transcripts per million (TPM) as indicated. Boxes above indicate IDR peaks, with red boxes indicating TIA1 RBNS motifs located within (or up to 50 bases upstream of) peaks.

### Each of the five assays used focused on a distinct aspect of RBP activity

#### Transcriptome-wide RNA binding sites of RBPs

We identified and validated hundreds of immunoprecipitation-grade antibodies that recognize human RBPs^17^ and developed enhanced CLIP (eCLIP)^18^. We identified high-quality eCLIP profiles for 120 RBPs in K562 cells and for 103 RBPs in HepG2 cells, for a total of 150 RBPs (of which 73 were characterized in both cell types) (Supplementary Data 4). In sum, this effort identified 844,854 significantly enriched peaks (relative to size-matched input controls for each RBP) that cover 18.5% of the annotated mRNA transcriptome and 2.6% of the pre-mRNA transcriptome.

#### RBP-responsive genes and alternative splicing events

To obtain insight into the functions of eCLIP peaks, we used shRNA- or CRISPR-mediated depletion followed by RNA-seq of 237 RBPs in K562 and 235 RBPs in HepG2 cells, for a total of 263 RBPs (of which 209 were characterized in both cell types) (Supplementary Data 5). Comparison against paired non-target control datasets identified 375,873 instances of RBP-mediated differential gene expression involving 20,542 genes affected upon knockdown of at least one RBP, as well as 221,612 cases of RBP-mediated alternative splicing events involving 38,555 alternatively spliced events impacted upon knockdown of at least one RBP. In addition to within-batch controls for each experiment, we performed batch correction to enable integrated analyses across the entire dataset (Extended Data Fig. 2).

#### In vitro RBP binding motifs

To identify the RNA sequence and structural binding preferences of RBPs *in vitro*, we developed a high-throughput version of RNA Bind-N-Seq (RBNS)^19^ that assays binding of recombinant purified RBPs to pools of random RNA oligonucleotides. In all, we identified the binding specificities of 78 RBPs^20^ (Supplementary Data 6). Short oligonucleotides of length *k*=5 (*k*mers) highly enriched in RBNS reads clustered into a single motif for about half of the RBPs assayed (37/78). The remaining RBPs had more complex patterns of binding, best described by two motifs (32/78), or even three or more motifs (9 RBPs). These data also indicate that many RBPs are sensitive to the sequence and RNA structural context in which motifs are embedded.

#### RBP sub-cellular localization

Post-transcriptional gene regulation occurs in different intracellular compartments. For instance, rRNA maturation and pre-mRNA splicing primarily occur in sub-regions of the nucleus, whereas mRNA translation and default mRNA decay pathways operate in the cytoplasm. To illuminate functional properties of RBPs in intracellular space, we took advantage of our validated antibody resource^17^ to conduct systematic immunofluorescence (IF) imaging of 274 RBPs in HepG2 cells and 268 RBPs in HeLa cells, in conjunction with a dozen markers for specific organelles and sub-cellular structures (Supplementary Data 1). These data, encompassing ~230,000 images and controlled vocabulary localization descriptors, have been organized within the RBP Image Database (http://rnabiology.ircm.qc.ca/RBPImage/).

#### RBP association with chromatin

Recent work has suggested that RBP association with chromatin may play roles in transcription and co-transcriptional splicing^21,22^. To generate a large-scale resource of chromatin association properties for RBPs, we performed ChIP-seq to identify the DNA elements associated with 30 RBPs in HepG2 cells and 33 RBPs in K562 cells for a total of 37 RBPs (of which 26 were characterized in both cell types) (Supplementary Data 7). These experiments identified 792,007 ChIP-seq peaks covering 3.8% of the genome.

To facilitate integrated analyses, all data for each data type were processed by the same data processing pipeline, and consistent, stringent quality control metrics and data standards were uniformly applied to all experiments. Although only 8 RBPs were investigated using all five assays, 249 of the 352 RBPs (71%) were studied using at least two different assays and 129 (37%) were subjected to at least three different assays, providing opportunities for integrated analysis using multiple datasets. As an example of how these complementary datasets provide distinct insights into RNA processing regulation, we considered *PTBP3* (also known as Regulator of Differentiation 1 / ROD1) (Fig. 1D). Inclusion of *PTBP3* exon 2 has been shown to alter start codon usage and increase cytoplasmic localization, and we observed by RNA-seq that *PTBP3* exon 2 inclusion was low in control cells but increased upon PTBP1 knockdown, consistent with previous studies^23^. This splicing event is likely directly regulated by PTBP1, as we observed eCLIP peaks at the 3’ splice site of *PTBP3* exon 2 which contained U-rich motifs shown to bind PTB family proteins by RBNS. Intriguingly, we also observe significant binding to *PTBP3* exon 10, which does not show alternative splicing itself but is orthologous to *PTBP1* exon 10 and *PTBP2* exon 11, which are each alternatively spliced in a PTBP1/2-regulated manner to generate transcripts targeted for nonsense-mediated mRNA decay^24^. Thus, it appears that the absence of PTBP1 regulation of *PTBP3* exon 10 splicing is not due to the loss of PTBP1 binding in this paralog. Considering mRNA levels, we observed that knockdown of TIA1 in K562 cells showed a 1.3-fold decrease in *PTBP3* mRNA, and that the *PTBP3* 3’UTR contained multiple eCLIP peaks for TIA1 in K562 cells, many of which overlapped with the TIA1 RBNS motif (UUUUU). This expression change paralleled the average change observed for many genes with TIA1 3’UTR eCLIP enrichment (see later discussion in Fig. 4). Similar integrated analysis can provide insight into mechanisms of cryptic exon repression and many other types of regulation. As an example, we observed HNRNPL eCLIP enrichment at a region downstream of a *GTPBP2* cryptic exon that contains repeats of the top HNRNPL RBNS motif, likely repressing splicing of the exon and contributing to production of *GTPBP2* mRNA with a full-length open reading frame (Extended Data Fig. 1c).

### Scalable quality assessment and analysis of eCLIP datasets

To generate the 223 high quality eCLIP datasets, we performed a total of 488 eCLIP experiments, each including biological duplicate immunoprecipitations along with a paired size-matched input (Fig. 2a, Extended Data Fig. 3-6, Supplementary Data 4, 8, 9 and 10). Quality assessment was performed manually using heuristics based on immunoprecipitation validation, library yield, presence of reproducible peak or repeat family signal, motif enrichment (for RBPs with known binding motifs), and consistency with well-characterized biological functions, yielding 223 eCLIP datasets released at the ENCODE Data Coordination Center (https://www.encodeproject.org). These manual quality assessments were then used to derive automated metrics that could accurately classify quality for 83% of eCLIP datasets (Extended Data Fig. 4). Datasets passing manual but not automated quality assessment were released with specific exceptions noted (Supplementary Data 8). An additional 50 datasets, which did not meet the stringent ENCODE standards but contained reproducible signal and could thus serve as useful entry points for future validation, have been deposited at the Gene Expression Omnibus (GSE107768) but were not included in the analyses described below (Extended Data Fig. 3c; Supplementary Data 9). We note that the eCLIP protocol does not include the direct visualization of protein-associated RNA that has been used in previous methods to assess whether RNA bound to co-purified RBPs of different size is present, and non-antigen IP of similar sized proteins is not easily detectable^18^. Although we have observed that UV crosslinking and stringent IP wash conditions generally limit the identification of indirect interactions, independent validation of peaks and binding properties identified by eCLIP through comparison with orthogonal *in vitro* motifs, knockdown-responsive changes, or other data types as described below therefore provides an essential validation to identify true binding signal.

**Figure 2.**
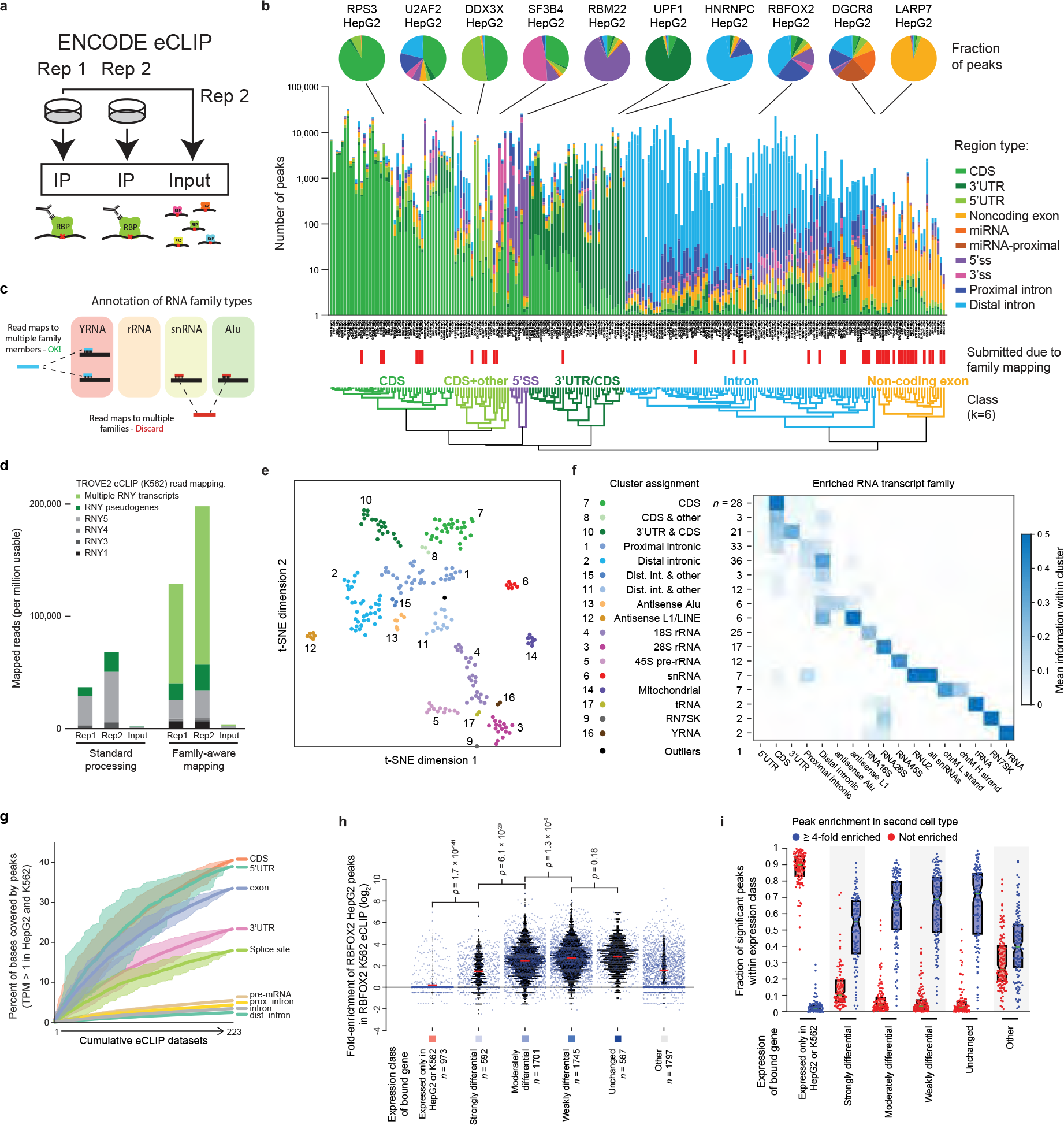
Integrated analysis of RBP:target association networks. (a) Schematic of ENCODE eCLIP experiments. (b) Stacked bars indicate the number of significantly enriched eCLIP peaks (with fold-enrichment ≥ 8, p-value ≤ 0.001, and meeting biological reproducibility criteria in RBP immunoprecipitation versus size-matched input). Number of peaks is shown on a logarithmic scale; bar heights are pseudo-colored based on the linear fraction of peaks that overlap the indicated regions of pre-RNA, mRNA, and non-coding RNAs. Datasets were hierarchically clustered to identify 6 clusters based on similar region profiles (Extended Data Figure 7a). (c) Model of eCLIP analysis pipeline for quantitation of eCLIP signal at RNA families with multiple transcript or pseudogene copies. (d) Stacked bars indicate the number of reads from TROVE2 eCLIP in K562 that map either uniquely to one of four primary Y RNA transcripts, map uniquely to Y RNA pseudogenes (identified by RepeatMasker), or (for family-aware mapping) map to multiple Y RNA transcripts but not uniquely to the genome or to other repetitive element families. (e) tSNE plot showing clusters of RBPs based on unique genomic as well as multicopy element signal. 16 clusters plus one outlier were identified using the MATLAB DBSCAN package. (f) For each cluster identified in (e), heatmap indicates the average relative information for RBPs in that cluster for each of the listed RNA regions or elements. (g) Lines indicate the cumulative fraction of bases covered by peaks for 100 random orderings of the 223 eCLIP datasets, separated by transcript regions as indicated. Shaded region indicates tenth through ninetieth percentiles. (h) Each point indicates the fold-enrichment in K562 eCLIP of RBFOX2 for a reproducible RBFOX2 eCLIP peak in HepG2, with underlaid black histogram. Peaks are separated based on the relative expression difference of the bound gene between K562 and HepG2: unchanged (fold-difference ≤ 1.2), weakly (1.2 < fold-difference ≤ 2), moderately (2 < fold-difference ≤ 5) or strongly (fold-difference > 5) differential (each of which required expression TPM ≥ 1 in both K562 and HepG2), or cell-type specific genes (TPM < 0.1 in one cell type and TPM ≥ 1 in the other). Mean is indicated by red lines, with significance determined by Kolmogorov-Smirnov test. (i) For each RBP profiled in both K562 and HepG2, points indicate the fraction of peaks in the first cell type associated with a given gene class that are (blue) at least four-fold enriched, or (red) not enriched (fold-enrichment ≤ 1) in the second cell type. Boxes indicate quartiles, with mean indicated by green lines.

Standard CLIP-seq analyses often identify thousands to hundreds of thousands of clusters of enriched read density (Extended Data Fig. 5, Supplementary Data 4). However, we previously showed that requiring enrichment in IP versus paired input experiments significantly improves specificity in identifying biologically relevant peaks by removing non-specific signal at abundant transcripts^18^. Thus, although data for all clusters identified from IP-only analysis has been made available, in this study we required stringent enrichment relative to input (fold-enrichment ≥ 8 and p-value ≤ 0.001). We further required that significant peaks be reproducibly identified across both biological replicates using an approach based off the Irreproducible Discovery Rate (IDR) method (Extended Data Fig. 5). Finally, we removed peaks overlapping 57 ‘blacklist’ regions (many of which contain either adapter sequences or tRNA fragments) that show consistent artefactual signal (Supplementary Data 11). Down-sampling analysis indicated that peaks were robustly detected at standard sequencing depth even in genes with low expression (TPM near or even below 1) (Extended Data Fig. 6).

Overlaying peaks onto GENCODE transcript annotations, we observed that peaks for most RBPs overlapped specific regions within transcripts, consistent with previous functional roles of many RBPs (Fig. 2b). Based on the dominant transcript region type bound, we clustered these RBPs into 6 “RNA type classes”, which provided reference comparisons for later peak-based analyses (Fig. 2b, Extended Data Fig. 7a, Supplementary Data 4). However, we observed that uniquely mapped reads represented a minority of the total for many eCLIP datasets, with the remainder coming from multi-copy elements including gene families with multiple pseudogenes (such as ribosomal RNA or Y RNA), retrotransposons, and other repetitive elements (Extended Data Fig. 7b). To quantify this signal accurately, we developed a family-aware mapping strategy which enabled quantitation of relative enrichment at mRNA versus other RNA types (Fig. 2c-d). Incorporating this approach, we observed clusters of RBPs dominated by rRNA or snRNA signal consistent with known functions, as well as unexpected clusters dominated by antisense Alu and L1/LINE signal that suggests an underappreciated role for retrotransposable elements encoded within protein-coding transcripts (particularly in the antisense orientation) in the global RBP binding landscape (Fig. 2e-g, Extended Data Fig. 7c-e).

### Saturation of the discovery of RNA processing events and regulatory sites

The scale of our data enabled us to query the degree to which we have saturated the discovery of eCLIP peaks and RBP-associated RNA processing events. In total, 20,542 genes were differentially expressed in at least one knockdown experiment, including 92.1% of genes expressed in both cell types and 91.8% of those expressed in at least one of the two (Extended Data Fig. 8a-c). Similarly, 17,839 genes had a peak in at least one eCLIP dataset, representing 84.2% of genes expressed in both cell types and 92.0% of those expressed in at least one (Extended Data Fig. 8a-c). Only 4,889 genes had eCLIP peaks from and were responsive to knockdown of the same RBP, suggesting that a large fraction of knockdown-responsive expression changes result from indirect effects, consistent with previous observations that only a relatively minor subset of RBPs affect RNA stability (see later discussion and Fig. 4). Similar analysis of alternative splicing changes revealed that differentially spliced events were saturated to a lesser degree than differentially expressed genes, likely because the transient nature of pre-mRNA reduces the window for detection by eCLIP, particularly for the many constitutively spliced exons that show incomplete inclusion upon knockdown of spliceosomal components. The significant variability observed in splicing event downsampling was driven by over 13,000 splicing changes in one knockdown dataset (the RNA helicase and spliceosomal protein AQR^25^ in K562 cells), which had nearly 3 times as many changes as the next largest dataset (Extended Data Fig. 8d).

Considering eCLIP alone, we observed a total of 25.8 Mb (2.6%) of annotated pre-mRNA transcripts covered by at least one reproducible eCLIP peak, representing 10.2 Mb (18.5%) of exonic and 15.6 Mb (1.7%) of intronic sequence (Extended Data Fig 8e-f). Restricting our analysis to genes expressed (TPM>1) in both cell types, 3.4% of annotated intronic sequence (2.4% of distal intronic, 4.3% of proximal intronic, and 17.9% of splice site), and 33.5% of annotated exonic sequences (39.0% of 5’ UTR, 40.6% of CDS, and 23.3% of 3’ UTR, respectively) were covered by at least one peak (Fig. 2g). We found that, although profiling a new RBP often resulted in greater increases in covered bases of the transcriptome than did re-profiling the same RBP in HepG2 or K562, re-profiling the same RBP in a more distinct cell type (H1 or H9 stem cells) yielded even greater increases, suggesting that many additional RBP binding sites remain to be detected in cell types distinct from K562 and HepG2 (Extended Data Fig. 8g-i). While these results are consistent with previous work suggesting that RNAs are often densely coated by RBPs^26^, it remains to be seen what fraction of these peaks mark regulatory interactions rather than constitutive RNA processing. Indeed, many peaks may reflect association of proteins that coat or transiently interact with RNAs as part of their basic function, such as interaction of RNA Polymerase II component POLR2G with pre-mRNAs, or recognition of splice sites by spliceosomal components.

Next, we evaluated whether RBP regulation is consistent across cell types. We observed that RBFOX2 eCLIP peaks with at least 8-fold enrichment in HepG2 cells were also typically enriched in K562 cells (average enrichment of 6.2-fold) if the target RNA was expressed within a factor of five of the level in HepG2 cells (Fig. 2h). Extending this to all 73 RBPs with eCLIP data in both cell types, 65.7%, 64.8%, and 62.7% of peaks in unchanging, weakly, or moderately differentially expressed genes, respectively, were enriched by at least 4-fold in the second cell type, and often overlapped a reproducible and significant peak call in the other cell type (Fig. 2i, Extended Data Fig. 8j-k). In contrast, an average of 46.3% of RBP peaks that showed no enrichment in the second cell type occurred in genes with cell type-specific expression (a 3.0-fold enrichment), whereas only 21.6% occurred in unchanging, weakly, or moderately differentially expressed genes, respectively (a 3.0-fold depletion) (Extended Data Fig. 8l). Thus, these results suggest that most RBP eCLIP signal is preserved across cell types for similarly expressed genes, whereas peak discrepancies often reflect cell type-specific RNA expression instead of differential binding.

### *In vivo* binding is determined to a substantial extent by *in vitro* binding specificity

Binding of an RBP to RNA *in vivo* is determined by the combination of the protein’s intrinsic RNA binding specificity and other influences such as RNA structure and protein cofactors. To compare the binding specificities of RBPs *in vitro* and *in vivo*, we calculated the raw enrichment (*R* value) of each 5mer in RBNS-bound sequences relative to input sequences and compared these to the corresponding enrichments of 5mers in eCLIP peaks relative to randomized locations in the same genes (*R*_eCLIP_). We focused on 5mers because most proteins analyzed by RBNS contained RRM or KH domains, which are known from structural studies to individually bind ~3-5 bases of RNA^27,28^. Significantly enriched 5mers *in vitro* and *in vivo* were mostly in agreement, with 15 of the 23 RBPs having significant overlap in the 5mers that comprise their motif logos (Fig. 3a, left). The top RBNS 5mer for an RBP was almost always enriched in eCLIP peaks of that RBP (Fig. 3a, center, Extended Data Fig. 9a). For 18 of 21 RBPs in well represented RNA type classes, the RBNS motifs explained more of the corresponding eCLIP peaks than of eCLIP peaks of other RBPs in the same RNA type class (Extended Data Fig. 9b-d). In most cases, similar degrees of enrichment and similar motif logos were observed in eCLIP peaks located in coding, intronic or UTR regions, suggesting that RBPs have similar binding determinants in each of these transcript regions (Fig. 3a, center; Extended Data Fig. 9e, 10a). Strikingly, the most enriched RBNS 5mer occurred in 30% or more peaks for several RBPs including SRSF9, TRA2A, RBFOX2, PTBP3, TIA1, and HNRNPC, and for most RBPs at least half of eCLIP peaks contained at least one of the top five RBNS 5mers. Therefore, instances of these 5mers provide candidate nucleotide-resolution binding locations for the RBP (Fig. 3a, right), which have applications including identification of genetic variants likely to alter function at the RNA level (see Extended Data Fig. 4 from Moore et al. ENCYCLOPEDIA Companion manuscript, in revision). When two or more distinct motifs were enriched in both RBNS and eCLIP, the most enriched motif *in vitro* was usually also the most enriched *in vivo* (5 out of 7 cases). These observations are consistent with the idea that intrinsic binding specificity observed *in vitro* explains a substantial portion of *in vivo* binding preferences for most RBPs, with the caveat that most RBNS data are from RBPs that contain single-stranded RNA-binding domains.

**Figure 3.**
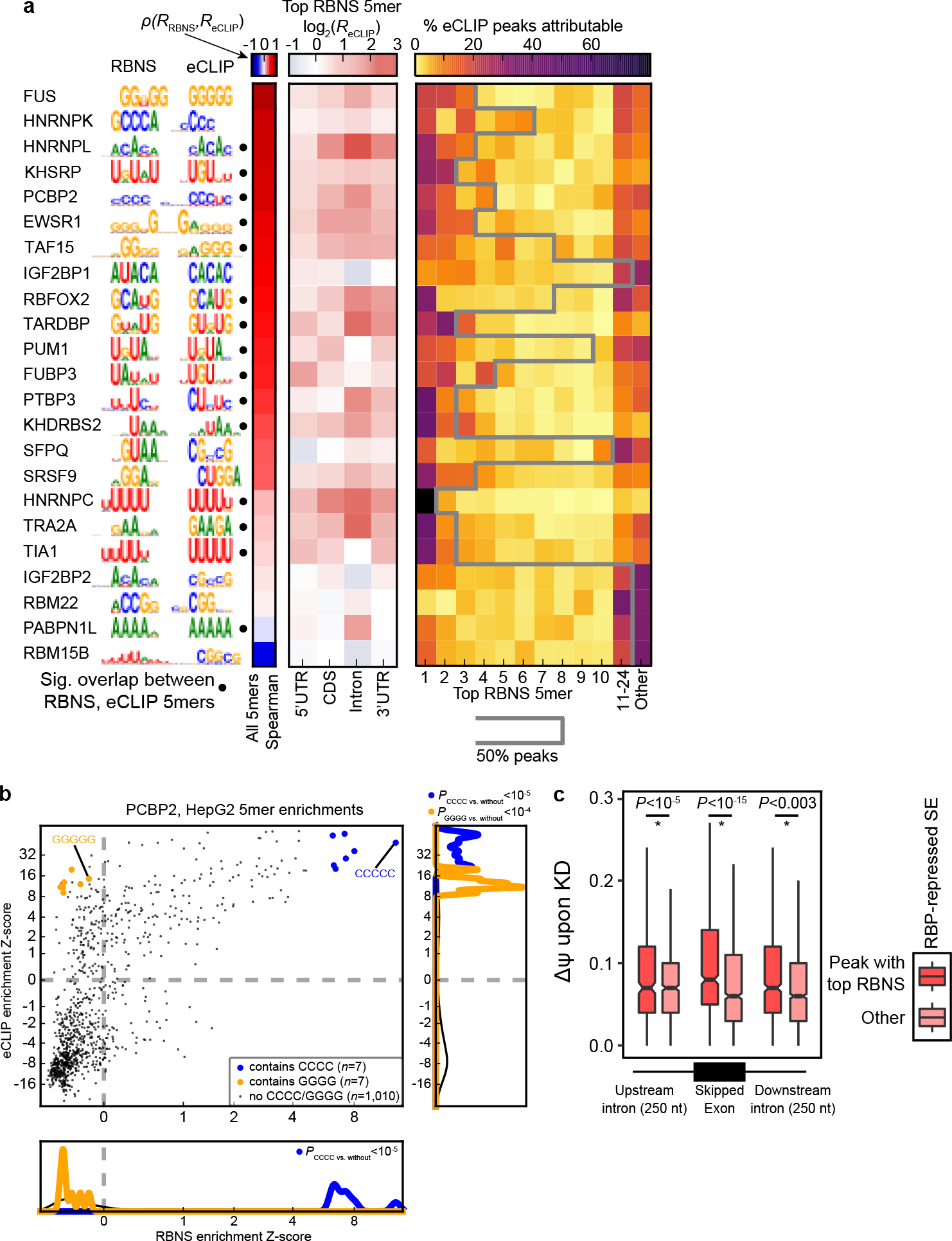
Sequence-specific binding in vivo is determined predominantly by intrinsic RNA affinity of RBPs. (a) Left: Top sequence motif of RBNS versus eCLIP-derived enriched 5mers clustered by similarity of RBNS motifs. Filled circles to the right of the eCLIP logo indicate if the groups of 5mers comprising the RBNS and eCLIP motifs overlap significantly (hypergeometric *P*<0.05). Center-left: Heatmap indicates correlation between RBNS and eCLIP enrichments for all 5-mers. Center: Enrichment of the top RBNS 5mer in eCLIP peaks (*R*_eCLIP_) within different genomic regions. Right: The proportion of eCLIP peaks attributed to each of the 10 highest affinity RBNS 5mers, as well as the #11-24 RBNS 5mers combined. The black line indicates the number of top RBNS 5mers required to explain >50% of eCLIP peaks for each RBP (maximum, 24 5mers). (b) Comparison of PCBP2 *in vivo* versus *in vitro* 6mer enrichments, with 5mers containing CCCC and GGGG highlighted. Significance was determined by Wilcoxon rank-sum test and indicated if P < 0.05. x- and y-axes are plotted on an arcsinh scale. Similar results were obtained when analyzing 6mers rather than 5mers. (c) Comparison of the magnitude of splicing change upon RBP knockdown for SEs containing eCLIP peaks with versus without the top RBNS 5mer, for RBP-repressed SEs grouped by the location of the eCLIP peak relative to the SE. The numbers of peaks for each region were as follows: exon peaks with RBNS motif: 368, without RBNS: 1758; upstream intron peaks with RBNS: 223, without RBNS: 2195; downstream intron peaks with RBNS 250, without RBNS 953. Significance was determined by Wilcoxon rank-sum test and indicated if P < 0.05.

**Figure 4.**
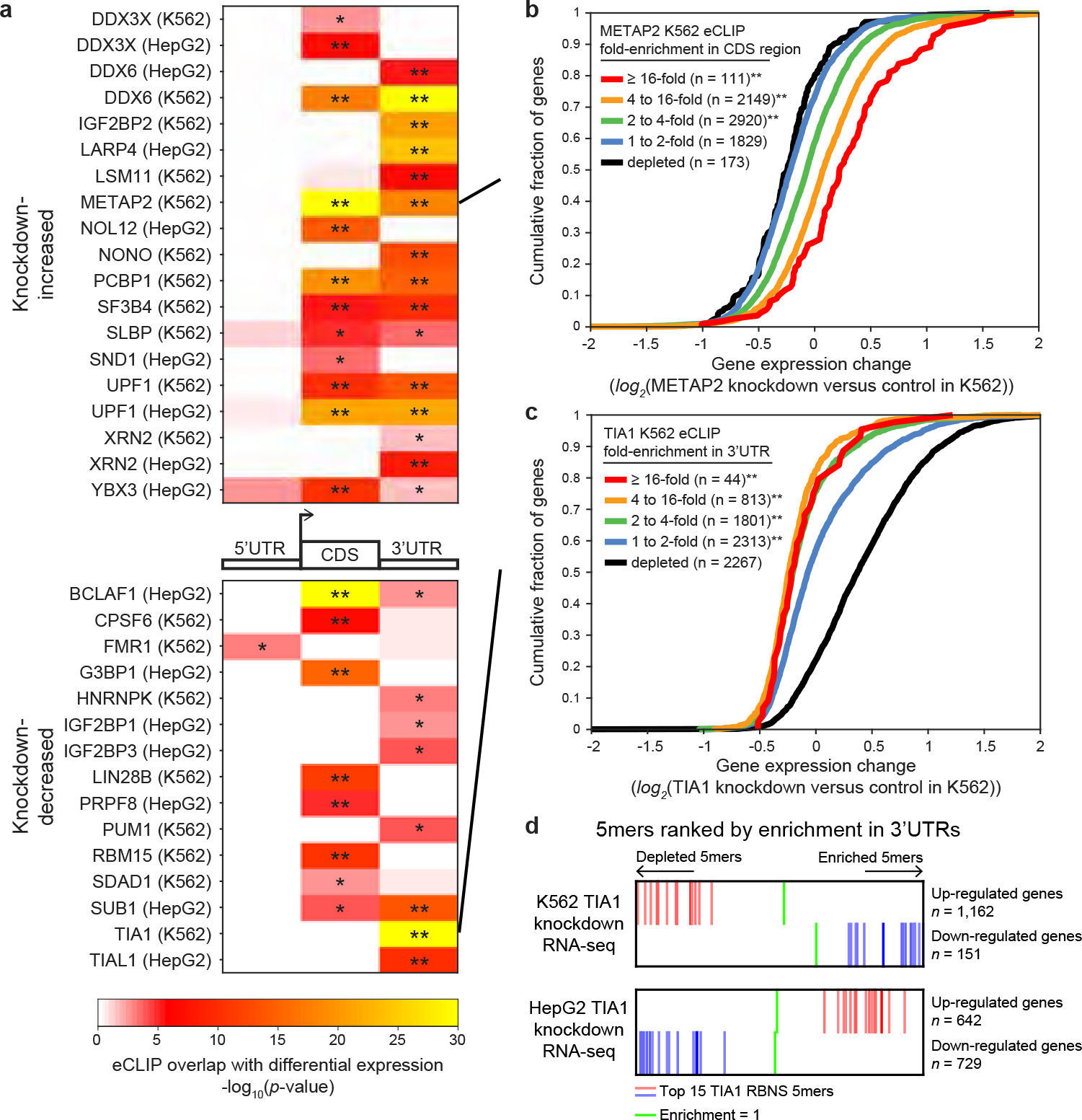
Association between RBP binding and RNA expression upon knockdown. (a) Heatmap indicates significance of overlap between genes with regions significantly enriched (*p* ≤ 10^−5^ and ≥4-fold enriched in eCLIP versus input) and genes significantly (top) increased or (bottom) decreased (*p* < 0.05 and FDR < 0.05) in RBP knockdown RNA-seq experiments. Significance was determined by Fisher’s Exact test or Yates’ Chi-Square approximation where appropriate; * indicates *p* < 0.05 and :: indicates *p* < 10^−5^ after Bonferroni correction. Shown are all overlaps meeting a *p* < 0.05 threshold; see Extended Data Fig. 14 for all comparisons. (b-c) Lines indicate cumulative distribution plots of gene expression fold-change (knockdown versus control) for indicated categories of eCLIP enrichment of (b) METAP2 in K562 and (c) TIA1 in K562. ** indicates *p* < 10^−5^ by Kolmogorov-Smirnov test. (d) Enrichment or depletion of the top 15 TIA1 RBNS 5mers in 3’UTRs of genes that are up− and down-regulated upon TIA1 knockdown in K562 and HepG2, relative to their frequency in control gene 3’UTRs (green lines indicate an enrichment of 1 (equal frequency in regulated gene 3’UTRs and control gene 3’UTRs)). All 1,024 5mers are ordered from lowest to highest enrichment from left to right in each row.

For slightly under half of the interrogated RBPs (10/23), the top five RBNS 5mers explained fewer than half of the eCLIP peaks. Some of these RBPs appear to have affinities to RNA structural features or to more extended RNA sequence elements not well represented by 5mers^20^, while the sequence-specific binding of others may be driven to a large extent by interacting proteins. In some cases, RBNS revealed affinity to a subset of the motifs that were enriched in eCLIP peaks. For example, C-rich 6mers were most enriched in PCBP2 RBNS data and also in PCBP2 eCLIP peaks (Fig. 3b). In this example, and in several others, a subset of similar eCLIP-enriched *k*mers were not enriched at all by RBNS (e.g., the G-rich 6mers in Fig. 3b). Such “eCLIP-only” motifs, which were often G-, GC-, or GU-rich (Extended Data Fig. 10b), may represent RNA binding of other proteins that interact with the targeted RBP – e.g., G-rich motifs enriched near RBFOX2 peaks may represent sites bound by HNRNPF, HNRNPH and HNRNPM in complex with RBFOX2^29,30^ – or could represent copurification or crosslinking artifacts or biases in the composition of genomic sequences located near crosslinked positions^31,32^. In the case of PCBP2, we observed that C-rich motifs but not G-rich motifs were enriched adjacent to PCBP2-regulated exons (Extended Data Fig. 11a-b). These observations support C-rich motifs but not G-rich motifs as sites of PCBP2-specific regulation.

The extent to which strength and mode of binding are reflected in eCLIP read density and regulatory activity is not well understood. We focused on regulation of splicing because a large proportion of the available cell type/RBP combinations that have knockdown/RNA-seq, eCLIP, and RBNS data involved RBPs with known roles in splicing, and splicing changes could be readily detected in the knockdown data. For most datasets involving knockdown of known splicing RBPs (18/28), eCLIP enrichment to one or more specific regions near alternative exons was associated with increased splicing changes upon knockdown of the RBP. In contrast, this association was observed for only one of the seven datasets involving RBPs that lacked known splicing functions (hypergeometric P<0.05, Extended Data Fig. 11c). To explore the relationship between sequence-specific binding and regulation, we classified eCLIP peaks as RBNS+ or RBNS− depending on whether they contained the highest-affinity RBNS motif (Supp. Methods). We then asked whether these classes of peaks differed in their association with splicing regulation. Examining exon-proximal regions commonly associated with splicing regulation, we found that RBNS+ eCLIP peaks were associated with stronger repression of exon skipping, with an average ~25% increase in change of exon inclusion (commonly referred to as change of Percent Spliced In, or ∆Ψ) than RBNS− peaks (Fig. 3c). Thus, eCLIP peaks that reflect sequence-specific binding appear to confer stronger regulation than other eCLIP peaks. Alternatively, such peaks may simply have a lower false positive rate, though the fairly stringent peak calling criteria used here make this explanation seem less likely. Either way, RBNS motifs can be used to distinguish a subset of eCLIP peaks that have greater regulatory activity. The *in vitro* data were needed to make this distinction, because a similar analysis of eCLIP peaks classified by presence/absence of the top eCLIP-only 5mer yielded minimal differences in splicing regulatory activity (Extended Data Fig. 11d). Unlike RBP-repressed exons, RBP-activated exons showed only a marginally significant (P<0.02) difference between RBNS+ and RBNS− peaks (in the opposite direction), not significant in either intronic region (Extended Data Figure 11e). Why a stronger effect should be observed for RBP-repressed than RBP-activated exons is not clear, though perhaps RNA binding directed by intrinsic RNA affinity may generally involve longer-duration interactions that more consistently impact (e.g., repress) recruitment of splicing machinery.

### Functional Characterization of RBP Maps

Analysis of the knockdown/RNA-seq data enables inference of the function of some RNA elements identified by eCLIP. First, we considered significant changes in transcript abundance identified upon RBP knockdown via RNA-seq (Extended Data Fig. 12-13). Regulation of RNA stability, which alters steady-state mRNA levels, can be observed by an increase or decrease in mRNA expression upon knockdown of an RBP. To identify potential regulators of RNA stability, we compared differentially expressed genes upon RBP knockdown with eCLIP enrichment in three regions of mRNAs: 5’UTR, CDS, and 3’UTRs. We observed that eCLIP enrichment for 15 RBPs (including 4 in both cell types) correlated with increased expression upon knockdown whereas eCLIP enrichment for another 15 RBPs correlated with decreased expression (Fig. 4a, Extended Data Fig. 14a). Comparing against RBPs of the same binding class (Fig. 2b), the targeted RBP showed the greatest enrichment in 14 out of 34 cases and was among the top RBPs for most comparisons (Extended Data Fig. 14b-c).

Correlation between eCLIP and genes with increased expression upon RBP knockdown included RBPs with previously identified roles in induction of RNA decay (such as UPF1, XRN2, and DDX6) (Fig. 4a, Extended Data Fig. 15a), as well as previously uncharacterized RBPs including METAP2, a methionyl aminopeptidase that has been co-purified with polyA-selected RNA but has no known RNA processing roles^14^. METAP2 eCLIP showed an average 3.4-fold enrichment in CDS regions, above the 2.4-fold average enrichment of 3’UTR and 1.2-fold depletion of intronic regions (Extended Data Fig. 15b-d). We further observed a trend in which increasing METAP2 eCLIP fold-enrichment correlated with progressively stronger increases in RNA expression upon knockdown, supporting an RNA regulatory role (Fig. 4b).

In contrast, the 15 RBPs for which eCLIP enrichment correlated with decreased RNA levels following knockdown (Fig. 4a) included stress granule components TIA1, TIAL1, and G3BP1 among other RBPs. Surprisingly, although our transcriptome-wide analysis indicated that transcripts with 3' UTR TIA1 eCLIP enrichment decreased upon knockdown in K562 cells (suggesting a globally stabilizing role for TIA1) (Fig. 4c), little to no stabilization activity was observed for mRNAs with 3' UTR enrichment for TIA1 in HepG2 cells (Extended Data Fig. 15e). Using TIA1 RBNS motif content in 3' UTRs rather than eCLIP enrichment, we additionally observed cell-type specific enrichment of TIA1 motifs in destabilized transcripts upon KD in K562, with no significant effect (though a slight motif enrichment in stabilized genes upon KD) in HepG2 (Fig. 4d, Extended Data Fig. 15f-g). This distinction is reminiscent of previous studies, which indicate that TIA1 can either induce RNA decay when tethered to a 3' UTR^33^, or stabilize target mRNA levels through competition with other RBPs including HuR^34^. Indeed, we observe that although TIA1-knockdown destabilized transcripts in K562 do not show correlated expression changes upon knockdown in HepG2, TIA1 eCLIP enrichment is similar between K562 and HepG2 for these transcripts (Extended Data Fig. 15h-i). Thus, our results provide further evidence that TIA1 can regulate mRNA stability through varying regulatory mechanisms that likely involve cell-type-specific co-factors.

### RBP association with splicing regulation

RBP binding to an exon (or its flanking introns) can regulate exon inclusion or exclusion, or alternative 5’ or 3’ splice site usage, through a variety of interactions with the splicing machinery^35^. To consider how RBP enrichment was associated with splicing regulation, we identified all significant alternative splicing events from comparison of RBP knockdown versus paired non-target control RNA-seq (Extended Data Fig. 16-17). Next, we generated an ‘RNA splicing map’ for each RBP^36^, in which the eCLIP enrichment in IP versus input is identified for all exons that increase (or decrease) exon inclusion upon RBP knockdown and then averaged to create a meta-exon plot (Extended Data Fig. 18). Comparison of these meta-exon plots can then reveal position-dependent regulation. For example, RBFOX2 eCLIP enrichment at the downstream proximal intron correlates with exon exclusion upon knockdown of RBFOX2 (Extended Data Fig. 18), consistent with previous studies of RBFOX2 motif enrichment and CLIP binding^37^. We performed this analysis for all 203 pairings of eCLIP and knockdown/RNA-seq performed in the same cell line (139 RBPs total) and we observed a wide variety of RNA maps for skipped exons (SEs, also referred to as cassette exons) (Fig. 5a-b, Extended Data Fig. 19a). Binding of SR proteins was typically associated with decreased SE inclusion upon knockdown while binding of hnRNP proteins was associated with increased SE inclusion upon knockdown, consistent with classical models of antagonistic effects of SR and hnRNP proteins on splicing^38^ (Extended Data Fig. 19b). We observed that the same RBP across cell types had higher splicing map correlation (particularly for knockdown-included exons) than random pairings of RBPs, with SR and hnRNP proteins contributing the majority of highly cross-correlated signals across RBPs (Extended Data Fig. 19c-e).

**Figure 5.**
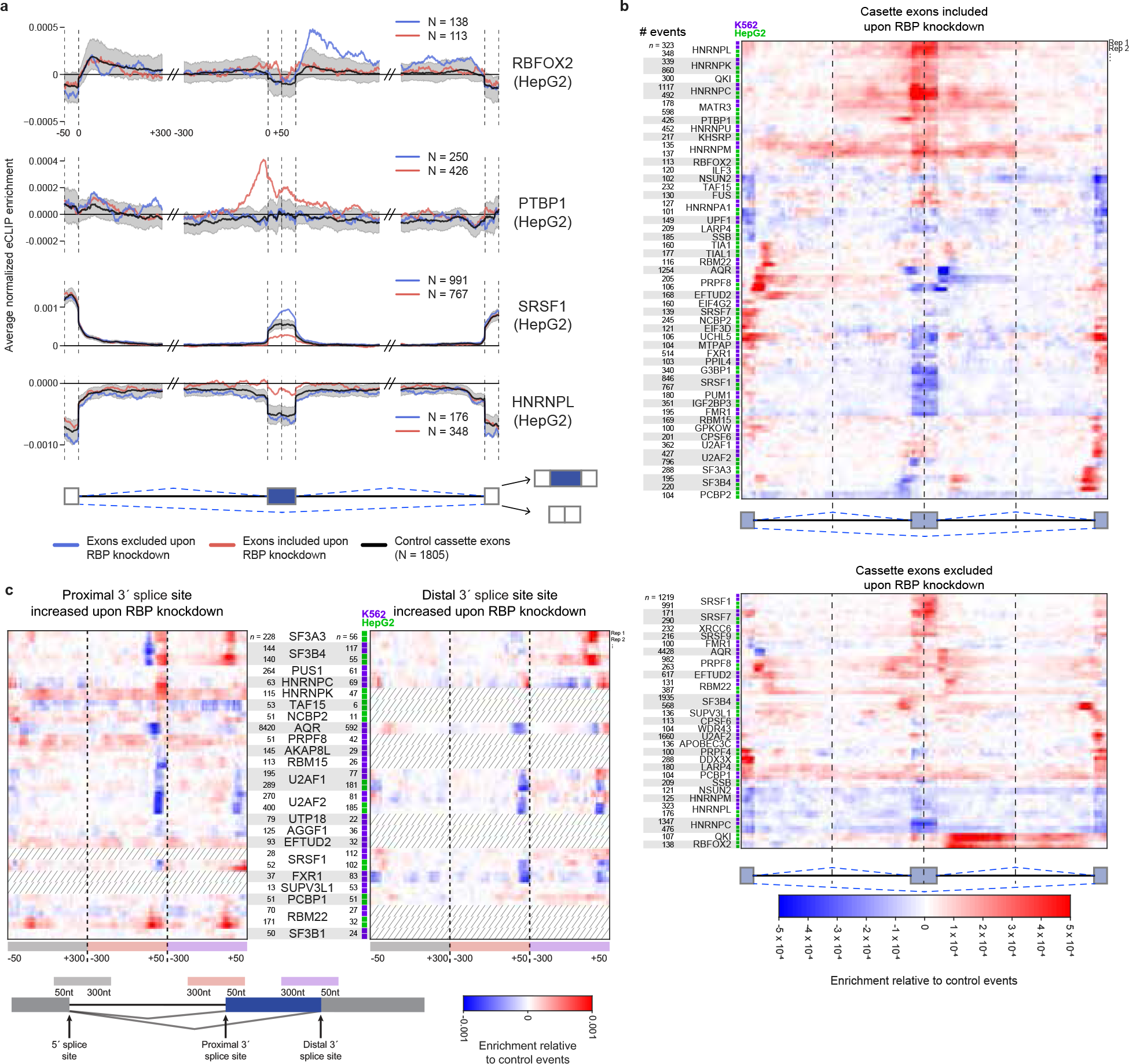
Integration of eCLIP and RNAseq identifies splicing regulatory patterns. (a) Normalized splicing maps of RBFOX2, PTBP1, SRSF1, HNRNPL for cassette/skipped exons (blue) excluded and (red) included upon knockdown, relative to a set of ‘native’ cassette exons (nSE) with 0.05 < inclusion rate < 0.95 in controls. Lines indicate average eCLIP read density in IP versus input for indicated exon categories. Shaded area indicates 0.5^th^ and 99.5^th^ percentiles observed from 1000 random samplings of native events. The displayed region shown extends 50 nt into exons and 300 nt into introns. (b) Heatmap indicates the difference between normalized eCLIP read density at cassette exons (top) included or (bottom) excluded upon RBP knockdown, versus native cassette exons. Shown are all RBPs with any position meeting *p* < 0.005 significance and 0.0002 normalized enrichment cutoffs (see Extended Data Fig. 19a for all RBPs). (c) As in (b), shown for RBP-responsive alternative 3’ splice site events relative to ‘native’ A3SS events with 0.05 < proximal 3’ splice site usage < 0.95 in controls. Dashed lines indicate datasets with less than 50 significantly altered events. The displayed regions include the upstream common 5′ splice site (grey box), the extended alternative 3′ splice site (orange box) and the distal alternative 3′ splice site (purple box)

RBPs that are components of the spliceosome displayed higher association at the upstream 5’ and downstream 3’ splice site at cassette exons and alternative exons sensitive to RBP-depletion (Fig. 5b, Extended Data Fig. 19f), consistent with previous observations of weaker splice sites flanking cassette exons^39^. When considering non-spliceosomal RBPs, we observed that RBP association was higher at cassette-bordering proximal intron regions relative to constitutive exons (CEs) that are always included, consistent with previous studies indicating increased RBP-mediated regulation of alternative events. Intriguingly, the upstream 5’ splice site showed an even greater enrichment than the intronic regions directly flanking the alternative exon (Extended Data Fig. 19f), suggesting that the 5’ splice site of the intron upstream of alternative exons represents an underappreciated regulatory region for RBPs.

As an additional control, we compared each knockdown dataset against all eCLIP datasets within the same RNA type class (as defined in Fig. 2b). Normalizing against this all-RBP background yielded overall similar splicing maps (Extended Data Fig. 20a). Whereas some individual RBPs, such as HNRNPC, showed only same-RBP enrichment (Extended Data Fig 20b), we observed that others indicated potential co-regulation. For example, when considering RBFOX2 knockdown-excluded exons we observed an enrichment for QKI slightly downstream of the RBFOX2-enriched region (Extended Data Fig. 20c). This appears to reflect complex coordination, as RBFOX2 and QKI rarely have enriched eCLIP signal for the same intron (Extended Data Fig. 20d) but we observe significant correlation in splicing changes upon RBFOX2 and QKI knockdown (*R*^2^ = 0.19, *p* = 1.2 × 10^−5^) (Extended Data Fig. 20e) which matches a previous observation in SKOV3ip1 ovarian cancer cells^40^. In contrast, we observe that TIA1 and TIAL1 show overlapping enrichment patterns at TIA1 knockdown-included exons (Extended Data Fig. 20f) despite little co-immunoprecipitation of the other factor (Extended Data Fig. 20g), confirming a previous observation showing similar iCLIP binding patterns of TIA1 and TIAL1^41^. However, TIA1 and TIAL1 knockdown-responsive exons show little correlation in splicing change (*R*^2^ = 0.03, *p* = 0.06) (Extended Data Fig. 20h), suggesting that although they share binding sites they may not share regulation at these sites. Thus, our results suggest that this approach may not only identify individual splicing regulatory patterns, but also provide insight into the regulatory relationships between RBPs.

Splicing maps constructed for alternative 5’ (A5SS) and alternative 3’ splice site (A3SS) events (Fig. 5c, Extended Data Fig. 21a-b) revealed differential association of spliceosomal components (Fig. 5c). We noted that branch point factors SF3B4 and SF3A3 interact at the branch point region ~50 nt upstream of the 3’ splices site. As a control set of native A3SS events, we utilized events which have both distal (upstream) and proximal 3’ splice sites in control shRNA datasets. When comparing the native set to A3SS events where the distal 3’ splice site has increased usage upon depletion of either SF3B4 or SF3A3, we find that average eCLIP enrichment for both proteins was decreased at the typical branch point location but increased towards the 3’ splice site (Extended Data Fig. 21c-d). Consistent with previous mini-gene studies showing that 3’ splice site scanning and recognition originates from the branch point and can be blocked if the branch point is moved too close to the 3’ splice site AG^42^, these results provide further evidence that use of branch point complex association to restrict recognition by the 3’ splice site machinery may be a common regulatory mechanism^43^ (Extended Data Fig. 21e).

In summary, the RBPs we have surveyed that participate in alternative splicing display a wide diversity of regulatory modes. Moreover, although the splicing events differ, the splicing map of a given RBP is often highly consistent between cell types. Thus, performing eCLIP and knockdown/RNA-seq in a single cell type may be sufficient to elucidate the general splicing rules for an RBP, but multiple cell types must be surveyed to identify the full repertoire of direct regulatory events.

### RBP Association with Chromatin

It is now broadly accepted that epigenetic marks can affect RNA processing through co-transcriptional deposition of splicing regulators, and conversely that regulatory RNAs interact with and coordinate regulation of chromatin and transcriptional states^21,44,45^. To explore further evidence of DNA association of specific RBPs, we selected RBPs for analysis based on their complete or partial localization in the nucleus and on the availability of antibodies and performed ChIP-seq to survey 58 RBPs in HepG2 and 45 RBPs in K562 cells for their association with DNA. 30 of 58 RBPs (52%) profiled by ChIP-seq in HepG2 and 33 of 45 RBPs (64%) in K562 showed significant reproducible ChIP-seq signal, with at least 200 (up to more than 50,000) peaks (Supplementary Data 7). These RBPs belong to a wide range of functional categories, including SR and hnRNP proteins, spliceosomal components and RBPs that have been generally considered to function as transcription factors, such as POLR2G and GTF2F1.

First, we characterized the RBP ChIP-seq peaks with respect to established chromatin features, including DNase I hypersensitive sites and various histone marks. This analysis revealed a general preference of RBPs for euchromatin relative to heterochromatin, especially gene promoters, although there was some variability among individual RBPs (Fig. 6a, Extended Data Fig. 22a). However, when we directly compared ChIP-seq peaks across RBPs we saw little overlap, with high concordance observed only for a small number of specific RBP pairs (Fig. 6b, Extended Data Fig. 22b). Collectively, even this moderately sized set of RBPs occupied ~30% of all DNase hypersensitive or open chromatin regions and ~70% of annotated gene promoters in both cell types. This is suggestive of broad interconnection between RBPs and actively transcribed regions in the human genome. Although some RBPs have been shown to also bind DNA, we note that this RBP-dependent specificity in ChIP-seq signal may instead reflect differential association of these RBPs with a variety of complexes containing transcription factors, epigenetic regulators, or other transcriptional machinery that binds DNA directly.

**Figure 6.**
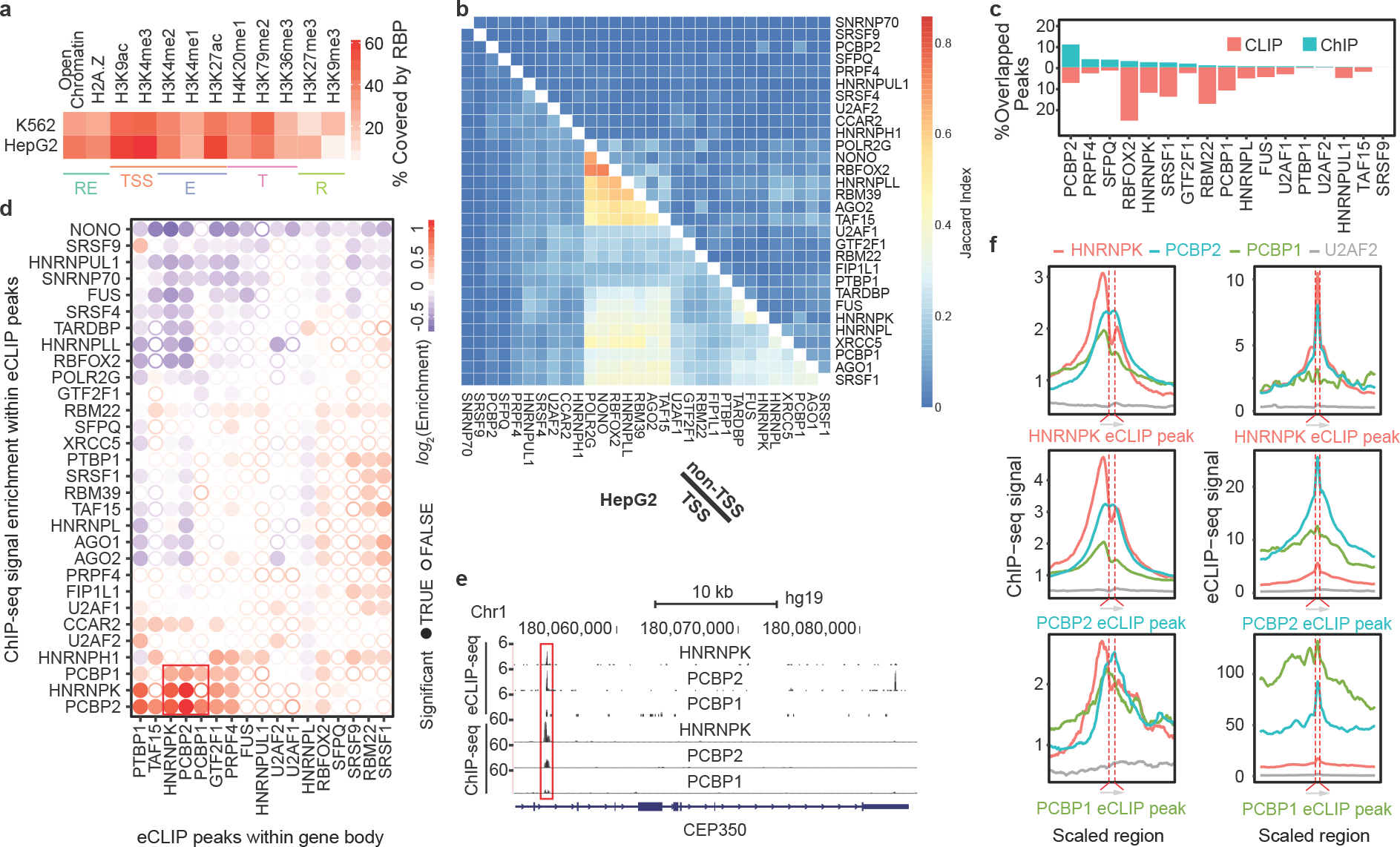
Chromatin-association of RBPs and overlap with RNA binding. (a) Overlap between RBP ChIP-seq and DNase I hypersensitive sites and various histone marks in HepG2 and K562 cells. Labels indicate marks associated with regulatory regions (RE), promoters (TSS), enhancers (E), transcribed regions (T) and repressive regions (R). (b) Heatmap indicates the Jaccard indexes between ChIP-seq peaks of different RBPs at (b)promoter regions (bottom left) or non-promoter regions (top right) for all HepG2 ChIP-seq datasets. See Extended Data Fig. 22b for K562 datasets. (c) Percentage of RBP eCLIP peaks overlapped by ChIP-seq peaks (red) or percentage of RBP ChIP-seq peaks overlapped by eCLIP peaks (green) for the same RBP. RBPs are sorted by decreasing level of overlapped ChIP-seq peaks. (d) Clustering of overlapped chromatin and RNA binding activities of different RBPs at non-promoter regions in HepG2. Color indicates the degree of ChIP enrichment at eCLIP peaks relative to surrounding regions. Significant enrichments (*p* ≤ 0.001) are indicated by filled circles. (e) A representative genomic region showing eCLIP and ChIP-seq signal for HNRNPK, PCBP2 and PCBP1 proteins in HepG2. (f) Cross-RBP comparison of chromatin and RNA binding activities in HepG2. Left: ChIP-seq density of indicated RBPs around HNRNPK, PCBP2 or PCBP1 eCLIP peaks. Right: eCLIP average read density of indicated RBPs around HNRNPK, PCBP2 or PCBP1 eCLIP peaks.

Next, we queried the degree to which DNA targets identified from ChIP-seq and RNA targets identified by eCLIP overlapped for the same RBP. Considering RBPs with both data types, we observed an average overlap of only 6% of eCLIP peaks and 2.4% of ChIP-seq peaks (Fig. 6c) (Supplementary Data 12). However, higher overlap was observed for a limited set of RBPs including the previously characterized DNA Polymerase II-interacting splicing regulator RBFOX2^46^. Focusing on non-promoter regions, we find that few RBPs displayed overlap between their ChIP and eCLIP signal, suggesting that ChIP signal reflects interactions with DNA or DNA-binding proteins independent of direct RNA binding for most RBPs (Fig. 6d). However, we observed an interesting association between poly(rC) binding proteins HNRNPK and PCBP1/2 (red box in Fig. 6d) which share a common evolutionary history and domain composition yet perform diverse functions^47^ and showed no clear overlap in ChIP-seq peaks at the global level but have overlap in ChIP-seq and eCLIP peaks at gene bodies (Fig. 6c, Extended Data Fig. 22b). To further explore the relationship between their RNA and chromatin interactions, we plotted the ChIP-seq and eCLIP read density of these three RBPs (as well as U2AF2 as an outgroup control) relative to PCBP1, PCBP2, and HNRNPK eCLIP peaks in non-promoter regions (Fig. 6e). We found that ChIP-seq signals were typically centered around eCLIP peaks, although HNRNPK (and to a lesser degree PCBP1) had a slight shift upstream of the eCLIP peak, which could reflect a specific topological arrangement of these potential RBP complexes on chromatin in a manner dependent on the direction of transcription (Fig. 6f, left panels). We observed that eCLIP signal also generally showed high overlap between these three RBPs but not unrelated spliceosomal component U2AF2 (Fig. 6f, right panels). Thus, these data suggest that although ChIP-seq signals for many RBPs may simply reflect pre- or co-transcriptional association at promoter regions, a subset show overlaps between both DNA and RNA targets within gene bodies that likely reflect distinct mechanisms of recruitment. Further work will be required to distinguish which of these potential interactions reflect single complexes, more complex recruitment modes, or simply reflect co-immunoprecipitation with other RBPs.

Finally, we investigated the potential for correspondence between DNA association and downstream effects on gene expression or splicing. First, we observed that the probability of ChIP-seq association correlated with increasing RNA expression levels for many RBPs, including DNA Polymerase II subunit POLR2G, suggesting that this may be a general property of RBPs which associate with the transcriptional machinery (Extended Data Fig. 22c, left). Next, we compared the frequency with which genes were differentially expressed upon RBP knockdown as a function of whether or not the RBP was chromatin-associated at that gene, using a background of randomly selected genes of similar expression level to control for the bias observed above. This analysis yielded a small number of RBPs (including HNRNPL and HNRNPLL) which showed significant enrichment for differential expression among ChIP-seq targets (Extended Data Fig. 22c, center). Performing the parallel analysis for differential alternative splicing events, we similarly observed significant overlap for three spliceosomal RBPs (RBM22, U2AF1, and SNRNP70) (Extended Data Fig. 22c, right). These data support the hypothesis that association of RBPs to chromatin is linked to downstream RNA processing, although the generally low odds ratios suggest the presence of additional properties that distinguish regulatory from non-regulatory interactions.

### RBP regulatory features in subcellular space

As RNA processing steps occur at an array of distinct locations within the cell, knowledge of the subcellular localization of each RBP is important to interpret the biological function of interactions or regulation observed in other assays. Our systematic immunofluorescence imaging screen revealed that RBPs display a broad diversity of localization patterns (Fig. 7a), with most factors exhibiting targeting to multiple structures in the nucleus and cytoplasm (Fig. 7b). Next, we integrated RBP localization features with other datasets generated in this study. To confirm the robustness of these orthogonal datasets, we first considered organelles with known roles in processing specific types of RNA. As expected, we observed significant overlap between localization of RBPs to nucleoli and eCLIP enrichment at the 45S precursor rRNAs and snoRNAs, mitochondria with enrichment at mitochondrial RNAs, and nuclear speckles with enrichment at proximal intronic regions (Fig. 7c). Nucleolar RBPs included 18 factors known to play roles in rRNA processing, including BOP1, UTP18, and WDR3. Intriguingly, we observed nucleolar localization for 15 additional RBPs with no annotated RNA processing function in humans (Supplementary Table 1), 3 of which showed enriched eCLIP signal at the 45S rRNA: AATF and PHF6, which both showed rRNA processing defects in a large-scale screening effort^48^, and UTP3, a human ortholog of yeast rRNA processing factor SAS10 (Extended Data Fig. 23a). Similarly, 14 out of 18 RBPs (78%) with at least 5-fold enrichment for one or more snRNAs exhibited nuclear speckle localization, whereas only 51% of all RBPs with both eCLIP and immunofluorescence data in HepG2 cells colocalized with speckles (*p* = 0.016 by Fisher Exact test). Focusing specifically on the nuclear to cytoplasmic ratios for each RBP, we observed a significant shift towards eCLIP signal at unspliced transcripts for nuclear RBPs, whereas cytoplasmic RBPs were enriched for spliced transcripts (Extended Data Fig. 23b-c). We also observed similar correspondence between RBP localization and altered RNA processing upon RBP knockdown. For example, analysis of splicing changes associated with RBP depletion revealed that speckle-localized RBPs impact larger numbers of splicing events compared to non-speckle associated proteins (Extended Data Fig. 23d), consistent with key roles of nuclear speckles in organization and regulation of the splicing machinery^49^.

**Figure 7.**
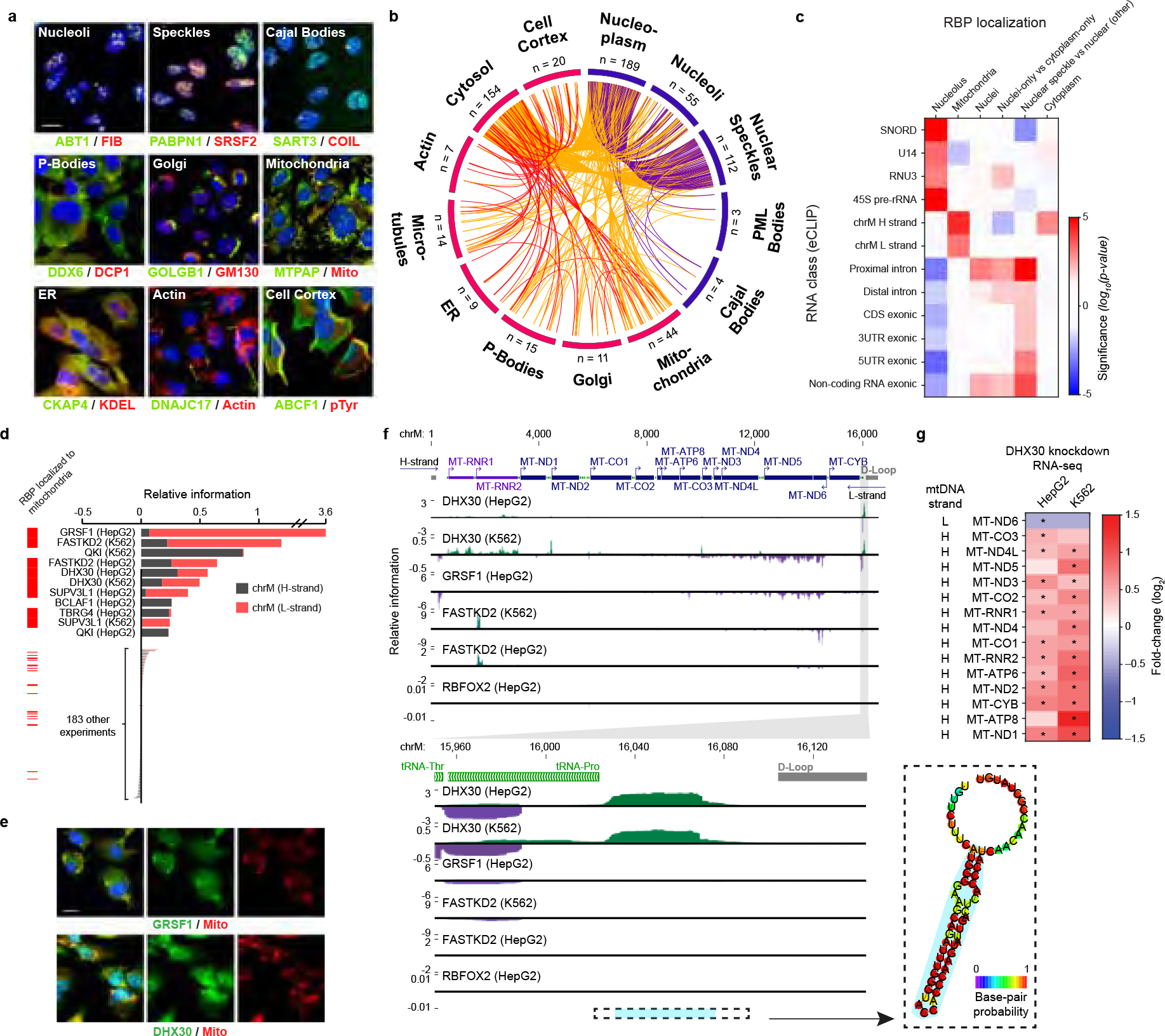
RBP sub-cellular localization features and their links to transcriptome binding and regulation. (a) Example RBPs (green) co-localized with nine interrogated markers (red). (b) Circos plot with lines indicating co-observed localization patterns (red: within cytoplasm; purple: within nucleus; orange: between cytoplasm/nucleus). (c) For localization patterns with known localized RNA classes, heatmap indicates significance (from Wilcoxon rank-sum test) comparing eCLIP relative information for the indicated RNA class (y-axis) for RBPs with versus without the indicated localization (x-axis). (d) Bars indicate eCLIP relative information content (IP versus input) for mitochondria H-strand (grey) or L-strand (red). (left) RBPs with mitochondrial localization in HepG2 are indicated in red. (e) Immunoflourescence images of mitochondrial localization of GRSF and DHX30. (f) Genome browser tracks indicate eCLIP relative information content along (top) the mitochondrial genome or (bottom) a ~300nt region for indicated RBPs. (right) Inset shows RNA secondary structure prediction (RNAfold) for the indicated region in blue. (g) Heatmap indicates gene expression change upon DHX30 knockdown for all mitochondrial protein-coding and rRNA transcripts. * indicates significant expression changes (p < 0.05 and FDR < 0.05 from DEseq analysis).

Focusing on localization to specific cytoplasmic organelles, we noted that 42 RBPs exhibited localization to mitochondria, an organelle with unique transcriptional and RNA processing regulation^50^. These mitochondrial-localized RBPs shared high overlap with RBPs with significant eCLIP enrichment on mitochondrial RNAs on either the Heavy (H) strand (QKI, TBRG4), Light (L) strand (GRSF1, SUPV3L1), or both strands (FASTKD2, DHX30), and mitochondrial localization by immunofluorescence was generally associated with significantly increased eCLIP enrichment on mitochondrial RNAs (Fig. 7d-e, Extended Data Fig. 23e). Next, we focused on DHX30, which is essential for proper mitochondrial ribosome assembly and oxidative phosphorylation^51^. Intriguingly, in addition to widespread association with many mitochondrial transcripts consistent with previous RIP-seq findings^51^ (Extended Data Fig. 23f), we observed dramatic enrichment at an unannotated region which has strong potential to form a stem-loop structure and is located on the mitochondrial H-strand downstream of all annotated genes and just upstream of the replication D loop (Fig. 7f). We further observed that DHX30 knockdown resulted in increased expression of nearly all H-strand transcripts, but decreased expression of L-strand transcript ND6 (Fig. 7g). As the termination signal for mitochondrial H-strand transcription has remained elusive, it is tempting to speculate that this site of DHX30 association could mark such a signal. These examples illustrate how intracellular localization of RBPs can be used as a powerful feature, in combination with binding and loss-of-function data, to infer aspects of post-transcriptional regulation that occur in different cellular compartments and organelles.

## Discussion

Our study represents the largest effort to date to systematically study the functions of human RBPs using integrative approaches. The resulting catalog of functional RNA elements substantially expands the repertoire of regulatory components encoded in the human genome. While the impact of DNA binding proteins mostly culminates in effects on gene expression levels, RBP function encompasses a broader range of activities. RBP functions extend outside the nucleus and into the cytoplasm and organelles, contributing to multiple paths by which RNA substrates are altered (splicing, RNA editing/modification, RNA stability, localization, translation), expanding transcriptome and proteome complexity. We demonstrate the effectiveness of combining *in vivo* maps of RNA binding sites identified with eCLIP with orthogonal approaches, such as *in vitro* evaluation of RNA affinity for the same RBPs, chromatin association by ChIP-seq, and functional assessment of transcriptome changes by RBP depletion and RNA-seq. At the molecular level, we confirm that *in vivo* and *in vitro* preferences are highly correlated for RBPs, and show that eCLIP peaks containing motifs reflective of intrinsic RNA affinity are more predictive of regulation. We confirm, using unbiased genome-wide analyses, that SR and hnRNP proteins have broadly antagonistic effects on alternative splicing. Moreover, we implicate the upstream 5’ splice sites of cassette exons in splicing regulation and extend previous findings that alternative 3’ splice site choice results from an “AG” scanning process that initiates with branch point recognition. We also implicate an RNA structure bound by an RBP in processing of mitochondrial transcripts, and elucidate new RNA splicing maps for many RBPs. Furthermore, our data provide the first systematic investigation of chromatin-associated gene regulation and RNA processing at the level of RBP-nucleic acid interactions. At the cellular level, immunofluorescence analysis with our extensive repository of RBP-specific antibodies place these molecular interactions within particular subcellular contexts. We confirm localization of many RBPs to nuclear speckles, mitochondria and other compartments, and identify many new proteins resident at these sites, emphasizing the necessity of localization data for interpreting RBP-RNA regulatory networks.

Here, we have surveyed the *in vivo* binding patterns of 150 RBPs, comprising the products of roughly 10% of the human genes predicted to encode proteins that interact directly with RNA. Within K562 and HepG2 cells, our observation that additional mapping of new RBPs continues to identify new RBP-associated regions argues that expansion of these approaches to additional RBPs will be particularly informative. Additionally, while we observe that *in vivo* binding patterns are highly consistent across genes expressed similarly in our two cell lines assayed (K562 and HepG2), our data indicates that mapping of previously characterized RBPs in drastically different cell types with highly distinct transcriptomes (particularly embryonic stem cells, post-mitotic cells such as neurons and muscle cells, or human tissues) will undoubtedly yield new discoveries. Additionally, RNA processing is dynamically regulated during acute or chronic environmental influences such as stress, as new binding sites may arise from both environmental changes in RBP or RNA concentrations, as well as from changes in post-translational modifications, binding partners, or subcellular distribution of RBPs. Thus, studying RBP subcellular localization and RBP-RNA substrate regulation under these conditions has potential to reveal new biology.

We expect that the data reported here will provide a useful framework upon which to build analyses of other aspects of RNA regulation, such as microRNA processing^52^, RNA editing and modifications such as pseudouridylation and m6A methylation, translation efficiency, and mRNA half-life measurements. We have yet to integrate *in vivo* RNA structure probing data to evaluate how RBP-mediated RNA processing are influenced by local^53^ and long-range RNA structures^54^. As we continue to embark on comprehensively characterizing all functional RNA elements, genome-scale CRISPR/Cas9 genome-editing^55^ and RNA modulation^56^ technologies will ultimately provide opportunities to study the impact on cellular and organismal phenotypes resulting from disruption of these RNA elements.

## Methods

### General information

Raw and processed datasets are accessible using accession identifiers provided in Supplementary Data 2 or can be found using the following publication file set accession identifiers at the ENCODE Data Coordination Center (https://www.encodeproject.org): eCLIP (ENCSR456FVU), knockdown RNA-seq (HepG2: ENCSR369TWP; K562: ENCSR795JHH; secondary analysis files including DEseq, rMATS, MISO, and CUFFDIFF output: ENCSR413YAF; batch corrected gene expression and splicing analysis: ENCSR870OLK), RBNS (ENCSR876DCD), and ChIP-seq (ENCSR999WIC). In addition to the methods described below, expanded experimental and computational protocols are linked to each experiment on the ENCODE DCC (https://www.encodeproject.org). All analyses in this manuscript used the *hg19* genome annotation and GENCODE v19 transcript annotations (unless otherwise noted), with *hg38* processed data available at the ENCODE DCC.

### RNA binding protein annotations and domains

RBPs were chosen from a previously described list of 1072 known RBPs, proteins containing RNA binding domains, and proteins characterized as associated with polyadenylated RNA, based on the availability of high quality antibodies^17^. Annotation of RBP function was performed by integration of published literature, with manual inspection of references for less well-established annotations. Annotation of RNA binding domain presence was determined by UniProt Domain Descriptions, and a database of cell-essential genes was obtained from published high-throughput CRISPR screening efforts^57^.

### eCLIP - experimental methods

Antibodies for eCLIP were pre-screened using a set of defined metrics^17^. A ‘biosample’ of HepG2 or K562 cells was defined as a batch of cells starting from a single unfrozen stock, passaged for less than 30 days under standard ENCODE reference conditions, and validated for high viability and non-confluent at the time of crosslinking. All cells within a biosample were pooled and UV crosslinked on ice at 400 mJoules/cm^2^ with 254 nm radiation. The biosample was then split into 20 million cell aliquots for eCLIP experiments.

eCLIP experiments were performed as previously described in a detailed Standard Operating Procedure^18^, which is provided as associated documentation with each eCLIP experiment on the ENCODE portal (https://www.encodeproject.org/documents/fa2a3246-6039-46ba-b960-17fe06e7876a/@@download/attachment/CLIP_SOP_v1.0.pdf). Briefly, 20 million crosslinked cells were lysed and sonicated, followed by treatment with RNase I (Thermo Fisher) to fragment RNA. Antibodies were pre-coupled to species-specific (anti-Rabbit IgG or anti-Mouse IgG) Dynabeads (Thermo Fisher), added to lysate, and incubated overnight at 4°C. Prior to immunoprecipitation (IP) washes, 2% of sample was removed to serve as the paired input sample. For IP samples, high- and low-salt washes were performed, after which RNA was dephosphorylated with FastAP (Thermo Fisher) and T4 PNK (NEB) at low pH, and a 3’ RNA adapter was ligated with T4 RNA Ligase (NEB). 10% of IP and input samples were run on an analytical PAGE Bis-Tris protein gel, transferred to PVDF membrane, blocked in 5% dry milk in TBST, incubated with the same primary antibody used for IP (typically at 1:4000 dilution), washed, incubated with secondary HRP-conjugated species-specific TrueBlot antibody (Rockland), and visualized with standard enhanced chemiluminescence imaging to validate successful IP. 90% of IP and input samples were run on an analytical PAGE Bis-Tris protein gel and transferred to nitrocellulose membranes, after which the region from the protein size to 75 kDa above protein size was excised from the membrane, treated with Proteinase K (NEB) to release RNA, and concentrated by column purification (Zymo). Input samples were then dephosphorylated with FastAP (Thermo Fisher) and T4 PNK (NEB) at low pH, and a 3’ RNA adapter was ligated with T4 RNA Ligase (NEB) to synchronize with IP samples. Reverse transcription was then performed with AffinityScript (Agilent), followed by ExoSAP-IT (Affymetrix) treatment to remove unincorporated primer. RNA was then degraded by alkaline hydrolysis, and a 3’ DNA adapter was ligated with T4 RNA Ligase (NEB). qPCR was then used to determine required amplification, followed by PCR with Q5 (NEB) and gel electrophoresis to size-select the final library. Libraries were sequenced on either the HiSeq 2000, 2500, or 4000 platform (Illumina). Each ENCODE eCLIP experiment consisted of IP from two independent biosamples, along with one paired size-matched input (sampled from one of the two IP lysates prior to IP washes).

### Experimental quality control of eCLIP experiments

eCLIP experiments for the ENCODE project were performed using two biological replicates, paired with a size matched input control subsampled from one of the two replicate samples (Extended Data Fig. 3a). Prior to sequencing, we utilized two metrics for assessing the quality of eCLIP experiments: successful immunoprecipitation of the desired RBP, and successful library generation and sequencing.

First, we required successful immunoprecipitation of the targeted RBP (assayed by IP-western blot analysis). This prerequisite first requires the identification of a RBP-specific immunoprecipitation-grade antibody, which we previously addressed by screening over 700 antibodies to identify 438 “IP-grade” antibodies against 365 RBPs in K562 cells^17^. Using these and other RBP antibodies validated by the RNA community, we performed 488 eCLIP experiments in K562 and HepG2 cell lines and observed successful immunoprecipitation during the eCLIP procedure for 400 (82%). 51 out of 270 (19%) and 37 out of 218 (17%) experiments gave failed IP-western blot results in K562 or HepG2 respectively, indicating either potential sensitivity to enzymatic steps and additional buffer exchanges performed during the eCLIP procedure, or a lack of expression in HepG2 cells (Extended Data Fig. 3b-c). IP-western images are provided for each ENCODE eCLIP experiment as part of the antibody metadata available at https://www.encodeproject.org.

Next, we assessed the quality of the amplified eCLIP sequencing library, as failure to obtain high-quality amplified libraries from both replicates can indicate a failed experiment, lack of RNA binding, or lack of RBP-RNA crosslinking. First, we abandoned 15 (4%) experiments that generated adapter-only sequencing libraries in either replicate. Next, we considered library complexity, defined as the fraction of unique RNA fragments relative to PCR duplicated fragments or other artifacts contained. Although library complexity is easily empirically calculated after sequencing and data processing, a quantitative metric for library complexity that can be applied prior to sequencing enables rapid culling of poor quality experiments and could help guide a desired sequencing depth by estimating an upper bound on the number of recovered RNA fragments. We previously introduced the extrapolated CT (eCT) metric that estimates the number of PCR cycles needed to obtain sufficient material for sequencing. This metric had appealing characteristics, as it was RBP-specific, showed high correlation with PCR duplication rate, and could be directly compared against eCLIP experiments performed with IgG isotype controls or antibodies in null cell lines^18,58^.

However, although the initial eCT calculation assumed an idealized 2-fold amplification rate per PCR cycle, we observed that this rate is frequently lower in practice. To properly estimate PCR efficiency during eCLIP, we noted that at our standard sequencing depths some experiments had saturated the discovery of unique fragments, which enabled us to accurately estimate the total number of pre-PCR unique fragments for these datasets. Using 6 datasets with a PCR duplication rate of greater than 90%, we observed that the best fit between the number of observed unique fragments and the estimated number of unique fragments occurred at a PCR efficiency of 1.84 (Extended Data Fig. 3d-e). We therefore defined an accurate-eCT (a-eCT) as the eCT calculated with 1.84-fold amplification per cycle instead of 2-fold.

To validate the a-eCT metric, we considered datasets that were beginning to saturate (PCR duplication rate greater than 60%). We observed that a-eCT showed strong predictive power for the number of unique RNA fragments observed (*R*^2^ = 0.46, *p* < 7.1 × 10^−38^) (Extended Data Fig. 3f), an improvement on the prior eCT metric (MSE 0.19 versus 0.86), confirming that a-eCT provides a robust estimate of library complexity (Extended Data Fig. 3g). Thus, a-eCT enables prediction of unique fragments prior to sequencing and indicates that eCLIP of distinct RBPs can yield a range from hundreds of thousands to billions of unique fragments (Extended Data Fig. 3h).

Next, we compared a-eCT against a manual annotation of experiment quality. We observed that experiments that pass manual quality assessment have a significantly lower a-eCT than experiments that failed manual quality assessment with mean a-eCTs of 13.3 versus 14.4 respectively (Extended Data Fig. 3i, students t-test; *p* < 10^−7^). Low a-eCT (corresponding to a highly complex library) did not always indicate high-quality eCLIP datasets, with failures due to poor reproducibility, lack of significant binding signal, and other failure modes. However, a high a-eCT value was a strong predictor of failure, typically due to a lack of the required number of unique fragments to produce reproducible peaks. To establish a maximum a-eCT threshold beyond which data are unreliable, we observed that the mean a-eCT for IgG control eCLIP experiments (which only immunoprecipitate background RNA) was 19.6. With that threshold applied, 21 out of 24 datasets with an a-eCT > 19.6 also independently failed manual QC. In all datasets examined no successful experiment had an a-eCT > 20.7, while there were still 9 experiments that did not pass manual quality control that had a higher a-eCT (Extended Data Fig. 3i).

In total, 331 out of 400 (83%) experiments had higher yield than this IgG-only value in both replicates, indicating successful immunoprecipitation of significant protein-bound RNA in the majority of experiments (Extended Data Fig. 3j). As we did observe a small number of high quality datasets with a-eCT values above this cutoff (typically RBPs with high specificity for a single or small number of RNA transcripts), we queried experiments with high a-eCT values with low-depth sequencing prior to full analysis and abandoned 36 such experiments which showed no significant binding specificity, leaving 349 datasets for analysis (Extended Data Fig. 3c).

### eCLIP - data processing and peak identification

Processing of raw eCLIP sequencing data is complex, as adapter sequences, double-adapter ligation products, retrotransposable elements and other multi-copy sequences, PCR duplicates, and underlying differences in RNA abundances all contribute to false negatives and false positives at both the read mapping and peak identification stages. To address these issues, we developed a rigorous standard eCLIP processing and analysis pipeline that was previously published18 and is provided (including description of steps as well as commands run) as a ‘Pipeline Protocol’ attached to each eCLIP dataset available on the ENCODE website at https://www.encodeproject.org/documents/3b1b2762-269a-4978-902e-0e1f91615782/@@download/attachment/eCLIP_analysisSOP_v2.0.pdf (Extended Data Fig. 5a).

Briefly, sequencing reads are first demultiplexed using dual indices with standard tools provided by Illumina. Next, reads were further demultiplexed based on in-line barcodes (present in read 1) (Supplementary Data 13). At this step, a unique molecular identifier (either N_5_ or N_10_) was removed from the beginning of read 2 and saved for use at the later PCR duplicate removal step. Next, potential adapter sequences were removed using cutadapt (v1.8.1), performed in two steps to properly remove non-full length adapter sequences we observed to drive artifact peak identification. At this step, reads with less than 18 bases were removed from further analysis. Next, we mapped reads using STAR (2.4.0i)^59^ against a database of repetitive elements (derived from RepBase (18.05)^60^ with the addition of elements including the 45S ribosomal RNA precursor), and removed reads with identified mapping (an independent method was derived to quantify mapping to repetitive elements, as described below). Reads were then mapped against the human genome using STAR (v 2.4.0i), requiring unique mapping (all analyses described in this manuscript used mapping to GRCh37 and GENCODE v19 annotations, but mapping to GRCh38 and GENCODE v24 annotations were also deposited at the ENCODE portal). PCR duplicate reads were then identified as those with the same mapped start position and unique molecular identifier and were removed using custom scripts to obtain unique fragments. Read clusters were identified using CLIPper^54^, which applies spine-fitting to identify clusters of enriched read density above local, transcript (both pre-mRNA and mRNA), and whole-genome background. Finally, clusters identified in IP samples were compared against paired size-matched input to obtain significantly enriched peaks. An average of 6.9% of clusters were significantly enriched, although this was highly variable across the 223 datasets (Extended Data Fig. 5b). The number of significantly enriched peaks was highly correlated between replicates, indicating the capture of RBP-specific biological signal (Extended Data Fig. 5c) (Supplementary Data 4).

To identify reproducible and significantly enriched peaks across biological replicates, we used a modified Irreproducible Discovery Rate (IDR) method (Extended Data Fig. 5d). IDR requires that peaks are ranked by an appropriate metric, but we found undesirable results ranking peaks by either significance (due to the dependence on underlying expression) or fold enrichment (due to the large variance of fold-enrichment when few reads are observed in input). Thus, we adapted relative entropy to better estimate the strength of binding in IP relative to input by defining the relative information content of a peak as 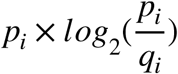, where *p*_*i*_ and *q*_*i*_ are the fraction of total reads in IP and input respectively that map to peak *i*. To confirm that this metric captures true binding signal, we considered the RBFOX2 eCLIP dataset in HepG2. We observed 14,595 reproducible clusters when we ranked by fold enrichment, whereas 32,431 clusters were reproducible when we ranked by information content (Extended Data Fig. 5e). Given the increased number of reproducible clusters detected, we used information content to perform standard IDR analysis to identify reproducibly bound regions^61^. We then identified the set of non-overlapping peaks from both replicates that maximized information content to define a final set of reproducibly enriched peaks that corresponded to CLIPper-identified regions (Extended Data Fig. 5d). Unless otherwise noted, the final set of reproducible and significant peaks was identified by requiring that the replicate-merged peak meet an irreproducible discovery rate cutoff of 0.01 as well as p-value ≤ 0.001 and fold-enrichment ≥ 8 (using the geometric mean of log_2_(fold-enrichment) and −log_10_(p-value) between the two biological replicates). Finally, 57 ‘blacklist’ regions were identified which we observed to be common artefacts across multiple datasets and lacked normal peak shapes (manual inspection indicated these often contain either adapter sequences or tRNA fragments) (Supplementary Data 11). IDR peaks overlapping these blacklist regions were removed to yield the final set of reproducible peaks used in all analyses in this manuscript (unless otherwise indicated) (Supplementary Data 4). This method revealed that an average of 53.1% of peaks identified as significantly enriched in individual replicates were significant and reproducible, indicating high reproducibility for most experiments (Extended Data Fig. 5f). Furthermore, the number of reproducible peaks identified upon profiling the same RBP in K562 and HepG2 cells was highly correlated, providing further validation that this approach reproducibly captures RBP-specific signal (Extended Data Fig. 5g).

Annotation of peaks was based on overlap with GENCODE v19 transcripts. If a peak overlapped multiple annotation types within a single annotated gene (across one or several isoform annotations), the peak annotation was chosen in the following priority order: tRNA, miRNA, miRNA-proximal (within 500 nt), CDS, 3’UTR, 5’UTR, 5’ splice site (within 100nt of exon), 3’ splice site (within 100nt of exon), proximal intron (within 400nt of splice site region), distal intron (within 400nt of splice site region), followed by noncoding exonic. If the peak overlaps multiple gene annotations, the final annotation was chosen as follows: tRNA, miRNA, CDS, 3’UTR, 5’UTR, miRNA-proximal, noncoding exonic, 5’ splice site, 3’ splice site, proximal intron, distal intron. To determine RBP clusters, the fraction of peaks annotated to each class out of the total number of peaks was calculated, and hierarchical clustering was performed in MATLAB (2018a) using correlation distance and average linkage. Clusters were obtained by cutting the tree at 6 clusters (chosen by comparing the sum of squared error between each dataset and the mean of all datasets within the cluster containing that dataset, which showed a leveling off after 6 clusters (Extended Data Fig. 7a)).

### Quantitation of eCLIP signal at multi-copy and other repetitive elements

A separate pipeline was developed to quantify enrichment for retrotransposable and other multi-copy elements. A database of multicopy elements was generated, including 5606 transcripts obtained from GENCODE v19 covering 34 abundant non-coding RNAs including rRNA, snRNA, and vault RNAs as their pseudogenes, 606 tRNA transcripts obtained from GtRNAdb (including versions with both genome flanking sequences and including the canonical CCA tail)^62^, 705 human repetitive elements obtained from the RepBase database (v. 18.05)^60^, 501 60-mer sequences containing simple repeats of all 1 to 6-nt k-mers, and the rRNA precursor transcript NR_046235.1 obtained from GenBank. Each transcript was assigned to one of 185 families of multi-copy elements (e.g. RNA18S, Alu, antisense Alu, Simple Repeat, etc.). Within each family, transcripts were given a priority value, with primary transcripts prioritized over pseudogenes.

Post-adapter trimming paired end sequencing reads were mapped to this repetitive element database using bowtie2 (v. 2.2.6) with options “-q --sensitive -a -p 3 --no-mixed – reorder” to output all mappings. Read mappings were then processed according to the following rules. First, for each read pair only mappings with the lowest mismatch score (least number of mismatches and insertions or deletions) were kept. Next, for equally scoring mappings within a repeat family described above, the mapping to the transcript with the highest priority was identified as the ‘primary’ match. Only read pairs mapping to a single repeat family were considered, whereas read pairs mapping with equal score to multiple repeat families were discarded from quantitation at this stage. Mapping to the reverse strand of a transcript was considered distinct to forward strand mapping, such that each family paired with a separate antisense family composed of the same transcripts with the same priority order (except for simple repeats, which were all combined into one family).

Next, repeat mappings were integrated with unique genomic mappings identified from the standard eCLIP processing pipeline (described above) as follows. If a read pair mapped both uniquely to the genome as well as to a repetitive element, the mapping scores were compared; if the unique genome mapping was more than 2 mismatches per read (24 alignment score for the read pair) better than to the repeat element, the unique genomic mapping was used; otherwise, it was discarded and only the repeat mapping was kept. Next, PCR duplicate removal was performed (similar to the standard eCLIP processing pipeline) by comparing all read pairs based on their mapping start and stop position (either within the genome or within the mapped primary repeat) and unique molecular identifier sequence, removing all but one read pair for read pairs sharing these three values. Finally, the number of post PCR-duplicate removal read pairs mapping to each multi-copy family was counted in both IP and paired input sample and normalized for sequencing depth (counting post-PCR duplicate read pairs from both unique genomic mapping as well as repeat mapping). Additionally, to better quantify signal to RepBase elements, RepeatMasker-identified repetitive elements in the hg19 genome were obtained from the UCSC Genome Browser. Element counts for RepBase elements were determined as the sum of repeat family-mapped read pairs plus uniquely genome mapped read pairs that overlapped RepeatMasked RepBase elements. After removing repeat-mapping elements, remaining reads were grouped and quantified based on transcript region annotations (CDS, 3’UTR, 5’UTR, proximal or distal intronic, non-coding exonic, intergenic, or antisense to GENCODE transcripts). Significance was determined by Fisher’s Exact test, or Pearson’s Chi-Square test where appropriate.

To summarize overall eCLIP signal, we applied the relative information content metric. Relative information content of each element in each replicate was calculated as 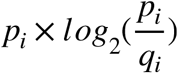, where *p*_*i*_ and *q*_*i*_ are the fraction of total reads in IP and input respectively that map to element *i*. A merged relative information for both replicates was calculated by defining *p*_*i*_ as the average fraction of total reads between the two biological replicates. To cluster datasets, dimensionality reduction was performed on element relative information from the combination of both replicates using the tSNE algorithm in MATLAB (2018a) with cosine distance, ‘exact’ algorithm, and perplexity = 10. To identify clusters, clustering was performed in using the DBSCAN (v1.0) MATLAB package, with options epsilon = 3 and MinPts = 2.

### Effect of sequencing depth on eCLIP peak identification

How deeply to sequence a CLIP-seq dataset is a major consideration (particularly at large scale), as samples must be sequenced sufficiently to robustly detect true binding signals while minimizing experimental cost. To query whether the ENCODE eCLIP datasets were sequenced to sufficient depth, we considered two questions: first, how does sequencing depth affect identification of true binding sites, and second, how many reads are required to detect binding sites in any gene when accounting for variability in gene expression.

First, we asked whether peaks discovered at deeper sequencing depths were still likely to be biologically relevant. To do this we looked at RBFOX2, which is known to bind to the GCAUG motif. Overall, we observed significant enrichment for RBFOX2 binding to its motif, with 36% of RBFOX2 peaks overlapping the motif versus a mean of 6% of peaks overlapping the motif in all other datasets (Extended Data Fig. 6a). We then down-sampled the unique genomic fragments, re-called peaks, and asked how many peaks discovered at each down-sampling step overlapped the RBFOX2 motif. We observed that peaks discovered using only 10% of unique genomic fragments showed the highest motif overlap (38% on average), whereas peaks that were only discovered when going from 90% to 100% of unique genomic fragments were less likely to contain GCAUG (27% on average) (Extended Data Fig. 6b). Although this suggests that signal to noise is highest among the most abundantly covered peaks, we note that later discovered peaks were still 3.0- to 7.4-fold enriched above non-RBFOX2 datasets, indicating they still contain significant true binding signal (Extended Data Fig. 6b). Supporting this, we observed that conservation of later-discovered peaks was similar to those discovered earlier with a mean phastcons conservation score of 0.136 versus 0.132 (Extended Data Fig. 6c). Considering an independent dataset, PRPF8, we observed similar results when testing its known association with the 5’ splice site: although peaks discovered at low sequencing depth were less enriched for true signal, we continued to see significant true positive signal throughout the range of down-sampling, indicating that it is true that deeper sequencing allows for the continued discovery of high quality peaks (Extended Data Fig. 6d-e).

Second, we considered the identification of peaks as a function of transcript abundance. To explore if there was a correlation between sequencing depth and the discovery of peaks in lowly expressed genes, we calculated the correlation between gene expression and the number of reads in each peak for RBFOX2. We observed that lowly expressed genes had fewer reads per peak (as expected), whereas highly expressed genes displayed a large variation in the number of reads per peak, with only a weak correlation overall for both RBFOX2 (*R*^2^ = 0.03) (Extended Data Fig. 6f). All other RBPs showed a similar week correlation (mean *R*^2^ = 0.13) (Extended Data Fig. 6g). Next, we asked whether peaks at lowly expressed genes could be detected at standard sequencing depths. Surprisingly, we found that lowly expressed genes (defined as those with TPM < 1) need on average only 670,000 unique genomic fragments to allow for detection of a peak in the gene, and this estimate was similar when varying the fraction of peaks required to be discovered or TPM thresholds (Extended Data Fig. 6h-j). As ENCODE eCLIP datasets have a mean sequencing depth of over 4.3 million unique genomic fragments, these results suggest that an inability to detect peaks on lowly expressed genes is not a major concern in eCLIP data sequenced to standard depths.

Our analysis above indicates that continued sequencing until fragment saturation can recover true peaks even at extremely high read depths. However, sequencing until fragment saturation is not typically economically reasonable. Thus, we set out to quantify diminishing returns upon deeper sequencing to identify whether we were observing saturation of detected peaks at typical eCLIP sequencing depths (Extended Data Fig. 6k-m). First, we developed a metric to quantify the diminishing returns of deeper sequencing in eCLIP datasets. Considering the discovery of significant peaks, we queried how many peaks were newly discovered when comparing peaks observed when 90% or 100% of fragments in a dataset were used to identify peaks. We observed that 67% of experiments passing manual QC saturated the discovery of significant peaks (defined as the discovery of fewer than 5% new peaks in the above metric), suggesting that simple peak detection was saturating for most but not all high-quality datasets (Extended Data Fig. 6k).

Next, we considered whether binding information by total information content was saturating even when peak discovery was not. Summing the total information content across all peaks, we observed that information recovered saturated for 97% of manually accepted datasets (using the same 5% or less discovery metric between 90% and 100% of fragments used to call peaks in a dataset) (Extended Data Fig. 6k). Exploring downsampling experiments further, we found that 90% of all eCLIP datasets that passed manual quality assessment had saturated information discovery by 8.5M unique fragments (corresponding to 4.3M unique genomic fragments) (Extended Data Fig. 6l-m). Thus, these results suggest that although additional peaks can be identified, the majority of peak information content is already captured at current sequencing depths for the majority of eCLIP experiments described here.

### Automated QC Metrics to verify data quality

We next developed a set of metrics to assess the quality of ENCODE eCLIP experiments. We ultimately arrived at two metrics for individual replicates (a minimal unique fragment cutoff, and a “total information in peaks” cutoff) as well as a third metric to assess reproducibility across the two biological replicates (Extended Data Fig. 4a). To evaluate these metrics, we used manual quality assessment of datasets to define a reference set of high- and low-quality eCLIP datasets.

The number of unique fragments per dataset varies widely, depending on library complexity and sequencing depth (as described above). We observed that a required number of 1.5M unique fragments maximized the predictive power for datasets passing manual quality assessment (f-score = 0.79) (Extended Data Fig. 4b). Only 7 of 446 manually passed datasets do not meet this threshold: two (TBRG4, PABPC4) are not yet saturated and thus could be rescued by re-sequencing, whereas the 5 other datasets (one replicate of SLBP, two replicates of SF3B1, and SUPV3L1) are already highly saturated, but were considered high quality due to presence of signal at a small number of specific RNAs matching previous studies of these RBPs (histones, the U2 snRNP, and mitochondrial RNA respectively)^63–65^. Although the classification power of this model is low (AUC = 0.57), datasets not meeting this threshold were more than 7 fold more likely to fail manual quality assessment (Extended Data Fig. 4c). Conversely, 30 of 222 manually failed datasets do not meet the criteria (Extended Data Fig. 4d).

Next, we considered a metric based on whether the dataset contained significant binding signal. As described above, we observed that the relative information of a peak better captures the binding information of peaks across genes with widely varying expression levels. Thus, to validate that a dataset contains significant binding information, we calculated the sum of relative information across all peaks in the dataset. We observed that this total information content score maximized the f-score of manually annotated high- and low-quality datasets at a total information content of 0.042 bits (f-score = 0.81) (Extended Data Fig. 4e). The information content model was more accurate (AUC = 0.71) (Extended Data Fig. 4f), accurately classifying 63% of ENCODE datasets with 0.36 specificity and 0.93 sensitivity (Extended Data Fig. 4g).

Next, we developed criteria to assay biological reproducibility, using two metrics based upon the Irreproducible Discovery Rate (IDR) approach that has previously been used to assay reproducibility of ChIP-seq peaks: reproducibility between real and pseudo-replicates (Rescue Ratio) and confirmation that the number of reproducible peaks between both replicates is similar (Self-Consistency Ratio)^66^. We found that cutoffs previously used for ChIP-seq data could be similarly applied to eCLIP^66^, and observed that 81.9% of experiments have a passing rescue ratio of <2 (Extended Data Fig. 4h) and 71.1% of experiments have a passing self-constancy ratio of <2 (Extended Data Fig. 4i). 223 experiments pass both thresholds, while 88 are borderline (passing one of the two thresholds), and 38 fail both thresholds (Extended Data Fig. 4j). Notably, these IDR metrics have high specificity, as 151 out of 196 (77%) of experiments that meet unique fragment and total information content cutoffs and were manually judged to be high quality passed this IDR criteria. In contrast, IDR detects potential false positives by correctly failing 9 out of 56 (16%) datasets that met read depth and information content metrics, but failed manual inspection (Extended Data Fig. 4j).

Finally, we combined these metrics into one overall automated quality call requiring that each experiment passes minimum read and entropy cutoffs as well as either being classified as passing or borderline based on IDR metrics (Extended Data Fig. 4k). Overall our model accurately classified 77% of eCLIP datasets with a sensitivity of 0.84 and a specificity of 0.62 (Extended Data Fig. 4l), better than any individual classification scheme.

### Quantitation of eCLIP signal at region level

For analyses using binding considered at the level of regions (e.g. 3’UTR, CDS, or proximal intronic), read density was counted for the indicated region for both IP and paired input, and significance was determined by Fisher Exact test (or Yates’ Chi-Square test if all observed and expected values were above 5). Only regions with at least 10 reads in one of IP or input, and where at least 10 reads would be expected in the comparison dataset given the total number of usable reads, were considered, and significant regions were defined as those with fold-enrichment ≥ 4 and p-value ≤ 0.00001.

### Knockdown followed by RNA-seq (KD/RNAseq)— experimental methods

Individual RBPs were depleted from HepG2 or K562 cells by either RNA interference (RNAi) or CRISPR-mediated gene disruption. RNAi was performed by transducing cells with lentiviruses expressing shRNAs (TRC collection) targeting an RBP followed by puromycin selection for 5 days. CRISPR-mediated gene disruption was performed by transfecting cells with a plasmid expressing Cas9 and a gRNA targeting an RBP, followed by puromycin selection for 5 days. In each case, knockdowns were performed in biological duplicate along with a pair of control knockdowns using a scrambled shRNA or gRNA. Protein was extracted from half of each sample and used to confirm knockdown of the target RBP by Western blotting. RNA was extracted from half of each sample and used to perform qRT-PCR to confirm knockdown of the targeted RBP transcript. We strived to obtain a knockdown efficiency of the target protein and/or RNA of at least 50% compared to the scrambled control, and for the knockdown efficiency to be within 10% between replicates. We used the extracted RNA to prepare RNA-seq libraries with the Illumina Tru-seq stranded mRNA library preparation kit. Paired-end 100 bp reads were generated from the RNA-seq libraries to an average depth of 63 million reads per replicate, and a minimum of 20 million reads per replicate, on an Illumina HiSeq 2500.

### KD/RNA-seq - data processing

#### Primary Data Processing

Reads were aligned to both GRCh37 using the GENCODE v19 annotations and GRCh38 using the GENCODE v24 annotations using both TopHat version 2.0.8^67^ with Bowtie2 version 2.1.0^68^, and STAR version 2.4.0^59^. All analyses described in this manuscript used the GRCh37/ GENCODE v19 alignments, but the GRCh38/GENCODE v24 alignments are also available at the ENCODE portal. In all cases, alignments were performed against the male reference genome sequence for HepG2 cells or the female reference genome for K562 cells and simultaneously to the ERCC spike-in sequences. The command line parameters for the TopHat alignments were: -a 8 -m 0 --min-intron-length 20 --max-intron-length 1000000 --read-edit-dist 4 --read-mismatches 4 -g 20 --no-novel-juncs --no-discordant --no-mixed. In some rare cases, TopHat 2.0.8 misassigned some reads to both strands or did not assign reads to either strand. To correct these errors, we used a custom script, tophat_bam_xsA_tag_fix.pl, to properly assign the SAM flag values. Gene expression levels were quantitated using RSEM (v1.2.23)^69^ and Cufflinks (v2.0.2)^70^. Only samples with a Pearson correlation coefficient on FPKM values of 0.9 or greater between replicates were used for further analysis. Samples with a correlation below 0.9 were repeated. We used the custom script (makewigglefromBAM-NH.py) to convert the single bam alignment files into plus or minus strand and unique and multi-mapped bam files, and then convert the intermediate bam files into bigwig files. A single, final bam file was generated per each RNA-Seq sample by merging the bam files containing the aligned read with that containing the unmapped reads. The merged bam and bigwig files were submitted to the ENCODE Data Coordination Center (https://www.encodeproject.org/). In total, 228 HepG2 knockdown experiments (221 shRNA and 7 CRISPR) and 224 K562 knockdown experiments (218 shRNA and 6 CRISPR) were used for further analysis.

#### Splicing Quantitation

Differential alternative splicing (AS) events were analyzed by rMATS (v 3.2.1.beta)^71^. The knockdown replicate bam files and their control replicate bam files with the Gencode v19 annotation file were analyzed using rMATS, to report five types of the differential AS events: SE (Skipped Exon), MXE (Mutually Exclusive Exons), A3SS (Alternative 3' Splice Site), A5SS (Alternative 5' Splice Site) and RI (Retained Intron). Events with abs(IncLevelDifference) > 0.05, PValue < 0.05 and FDR < 0.05 were identified as significantly differentially expressed AS events.

MISO (Mixture of Isoforms) (v misopy-0.5.2)^72^ was used to detect differentially processed Tandem 3’ UTR events (alternatively poly(A) site usage). Four pairwise comparisons between the two knockdown samples and two controls were run using compare-miso: KD-rep1 versus CN-rep1, KD-rep1 versus CN-rep2, KD-rep2 versus CN-rep1 and KD-rep2 versus CN-rep2. Significant Tandem 3’ UTR events were identified if abs(basin-factor) ≥ 5 and p-value < 0.05 on both the more_reads(KD-rep1, KD-rep2) versus fewer_reads(CN-rep1, CN-rep2) comparison and the fewer_reads(KD-rep1, KD-rep2) versus more_reads(CN-rep1, CN-rep2) comparison.

For the purposes of simplifying the analysis, we considered significant differential alternative splicing levels to be strong if |∆Ψ| ≥ 30%, moderate when 15% ≥ |∆Ψ| < 30%, and weak if 5% < |∆Ψ| < 15%.

#### Gene Expression Quantitation

Both DESeq (v1.28.0)^73^ and Cuffdiff (v2.2.0)^74^ were used to perform differentially expressed (DE) gene analysis with the KD RNA-Seq data. The expected_count values from the RSEM^69^ gene quantitation results of the STAR alignments were analyzed with DESeq and significant differentially expressed genes were identified as those with a p-value < 0.05 and padj < 0.05. In parallel, CuffDiff was used to analyze the TopHat aligned files using the default pooled Cross replicate dispersion estimation method and the multi-read-correct option. Genes were defined as significant if p-value < 0.05 and q-value < 0.1. Finally, a gene was reported as significant only if it was defined as a significant DE gene by both DESeq and Cuffdiff.

For the purposes of simplifying the analysis, we considered significant differential expression to be strong if the |log_2_(Fold-Change)| ≥ 2, moderate when 1< |log_2_(FoldChange)| < 2, and weak when |log_2_(FoldChange)| ≤ 1.

#### Batch Normalization of RBP knockdown RNA-Seq data

Batch effects are common in large datasets and must be corrected and accounted for^75^. We therefore designed our knockdown RNA-seq experiments at the onset of the project in way that would allow us to correct for batch effects. To do this, for each batch of experiments performed on a given day, we utilized the same scrambled shRNA or gRNA as non-specific controls alongside a batch of experimental shRNAs or gRNAs targeting a set of RBPs. This provided us with consistent non-specific control experiments in every batch that could be used to normalize data downstream. In addition to biological controls, if a given batch of biological samples was too large to make all the RNA-seq libraries in parallel, we made libraries from the non-specific control RNA samples in each sub-set of libraries made from a given biological batch. Analyses comparing eCLIP peaks with gene expression or alternative splicing changes in RNA-seq upon RBP knockdown used changes identified relative to these within-batch paired controls. However, to enable further integrated analyses, we performed additional batch correction as described below.

#### Batch Correction for Gene Expression Analysis

Gene expression batch effects were reduced with ComBat75. The HepG2 and K562 samples were normalized separately. Genes whose expected counts were 0 on more than 80% samples of the set were filtered out prior to normalization. After ComBat normalization and Quantile Normalization, the normalized values were rounded to integers and flatted to zeros if the values were less than zero. Instead of the original control samples, two “virtual” control replicates were created by averaging the normalized expression values of all rep-1 control samples or all rep-2 control samples for each gene. Then, DESeq was used to quantitate differential expression between the normalized knockdown samples and the virtual control samples. The batch normalized gene expression results are available at www.encodeproject.org (See Supplementary Data 2 for accession identifiers).

#### Batch Correction for Splicing Analysis

ComBat^75^ was also used to reduce batch effects for alternative splicing analysis. The HepG2 and K562 samples were normalized separately. The inclusion junction counts and skipping junction counts of all samples were collected from rMATS temporary files and used to form a table with each sample in columns. “Noise” junction counts were filtered out if their values were 0 on more than 80% samples. After ComBat batch normalization and Quantile Normalization on the filtered datasets, the normalized values were rounded to integers and flatted to zeros if the values were less than zero. Next, normalized rMATS temporary files were formed using the normalized junction counts and the “noise” junction counts. Instead of the original control samples, two “virtual” control replicates were created by averaging the normalized or “noise” junction counts of all rep-1 control samples or all rep-2 control samples for each event. rMATS was then resumed on the normalized knockdown samples and the virtual control samples to detect differential alternative splicing events. The batch normalized splicing results are available at www.encodeproject.org (See Supplementary Data 2 for accession identifiers).

### RNA Bind-N-Seq (RBNS) - experimental methods

RBNS experiments were performed as indicated in the protocol included on each experiment at the ENCODE portal. Briefly, randomized RNA oligonucleotides (20 or 40 nt) flanked by constant adapter sequences were synthesized and incubated with an SBP-tagged recombinant RBP (consisting minimally of all annotated RNA binding domains) at several concentrations (typically five, ranging from 5-1300 nM). RNA-protein complexes were isolated with streptavidin-conjugated affinity resin and eluted RNA was prepared for deep sequencing, resulting in 10-20 million reads per RBP pulldown concentration with a similar number of input reads sequenced per *in vitro* transcription reaction.

### RBNS - data processing

RBNS *k*mer enrichments (*R* values) were calculated as the frequency of each *k*mer in the pulldown library reads divided by its frequency in the input library; enrichments from the pulldown library with the highest individual *k*mer R value were used for each RBP. Mean and SD of *R* values were calculated across all *k*mers for a given *k* to calculate the RBNS Z-score for each *k*mer. RBNS pipeline source code is available at: https://bitbucket.org/pfreese/rbnspipeline.

RBNS motif logos were made using the following iterative procedure for *k*=5: the most enriched 5mer was given a weight equal to its excess enrichment over the input library (=*R*−1), and all occurrences of that 5mer were masked in both the pulldown and input libraries to eliminate subsequent counting of lower-affinity ‘shadow’ 5mers (e.g., GGGGA, shifted by 1 from GGGGG). All enrichments were then recalculated on the masked read sets to obtain the most enriched 5mer and its corresponding weight, with this process continuing until the enrichment Z-score (calculated from the original *R* values) was less than 3. All 5mers determined from this procedure were aligned to minimize mismatches to the most enriched 5mer, with a new motif initiated if the number of mismatches + offsets exceeded 2. The frequencies of each nucleotide in the position weight matrix, as well as the overall percentage of each motif, were determined from the weights of the individual aligned 5mers that went into that motif; empty unaligned positions before or after each aligned 5mer were assigned pseudocounts of 25% of each nucleotide, and outermost positions of the motif logo were trimmed if they had >75% unaligned positions. To improve the robustness of the motif logos, the pulldown and input read sets were each divided in half and the above procedure was performed independently on each half; only 5mers identified in corresponding motif logos from both halves were included in the alignments to make the final motif logo. In Fig. 3a, only the top RBNS motif logo is shown if there were multiple logos (all motifs displayed on the ENCODE portal within the “Documents” box of each experiment).

### Immuno-Fluorescence, Microscopy Imaging and Data Processing

HepG2 cells were seeded in Poly-L-Lysine coated 96-well clear bottom plates (Corning Inc; plate number 3882 half-area microplates), at a concentration of 2,000 cells per well in DMEM + 10% FBS. After 72h in standard growth conditions (i.e. 37°C and 5% CO_2_), cells were fixed with 3.7% formaldehyde, permeabilized in PBS + 0.5% Triton X-100 and blocked in PBS + 0.2% Tween-20 + 2% BSA (PBTB), all conducted for 20 min at room temperature. Primary antibodies directed against specific RBPs (all rabbit antibodies) and marker proteins were subsequently applied to the cells at a final concentration of 2 μg/mL in PBTB and incubated overnight at 4°C. The cells were next washed 3 times for 10 min each in PBST and incubated with secondary antibodies (Alexa647 donkey anti-rabbit and Alexa488 donkey anti-mouse, both diluted 1:500 in PBTB) for 90 min at room temperature. After 3 PBTB washes, the cells were counter stained with DAPI for 5 min, washed 3 times in PBS and stored in PBS at 4°C. Subcellular marker antibodies and dilutions used are as follows: rat anti-Alpha Tubulin, MCA78G, 1:200 (Serotec, Bio-Rad); mouse anti-CD63, ab8219, 1:200 (Abcam); mouse anti-Coilin, GTX11822, 1:100 (GeneTex Inc); mouse anti-DCP1a, sc100706, 1:200 (Santa Cruz Biotechnology); mouse anti-Fibrillarin, ab4566, 1:200 dilution (Abcam); mouse anti-GM130, #610822, 1:200 (Becton Dickinson); mouse anti-KDEL, ENZSPA827D, 1:200 (Enzo Life Sciences); mouse anti-Phospho Tyrosine, #9411S, 1:200 (NEB); mouse anti-PML, sc-966, 1:50 (Santa Cruz Biotechnology); mouse anti-SC35, GTX11826, 1:200 (GeneTex Inc). For staining with Mitotracker (Molecular Probes, M22426), cells were incubated with 100nM of dye in tissue culture media for 45 min at 37°C prior to fixation. For staining with Phalloidin (Sigma, P5282), cells were incubated with 50ug/ml of Phalloidin for 20 min prior DAPI staining.

Imaging was conducted on an ImageXpress Micro high content screening system (Molecular Devices Inc). For each RBP/marker combination, 10-20 high resolution images were acquired in the DAPI, FITC and Cy5 channels, using a 40x objective. Automated laser based auto-focusing and auto-exposure functions were employed for sample imaging, with exposure times ranging from 250-3000ms, 100-500ms and 50-100ms, for RBP, Marker and DAPI channels respectively. Raw unprocessed grayscale images from individual channels were acquired as high resolution TIF files of 726kb each. An in house Matlab script was developed to batch normalize image intensity values and add blue, green or red colors to the respective channels, which were subsequently merged as colour JPEG files. The final images were uploaded on a server accessible through the RBP Image Database website. A MySQL relational database (version 5.1.73) was implemented, along with a MyISAM storage engine, to store the images, data annotations and characteristics. A controlled vocabulary of descriptors was devised to document RBP subcellular localization features.

Image analysis to quantify nuclear/cytoplasmic staining ratios, or to assess the degree of RBP targeting to punctate subcellular structures (e.g. Cajal bodies, nuclear speckles, nuceloli, Golgi, P-bodies), was conducted using ‘Granularity’, ‘Colocalization’ and ‘Multi Wavelength Cell Scoring’ analysis modules from the MetaXpress v3.1 software (Molecular Devices Inc), according to manufacturer recommendations. For localization categories including microtubules, actin, cell cortex, ER, focal adhesions, mitochondria and mitotic apparatus, manual localization grading was conducted by ranking candidate RBPS as strongly or weakly co-localized with respective protein markers. The Circos plot of localization co-occurrance (Fig. 7b) was generated by drawing one line between every pair of categories for each RBP that shared both localization annotations. Nuclear annotations are indicated in purple, cytoplasmic in red, and lines between nuclear and cytoplasmic annotations are indicated in yellow.

### ChIP-seq - experimental methods

Chromatin immunoprecipitation was implemented according to ChIP Protocol optimized for RNA-binding proteins (http://www.encodeproject.org/documents/e8a2fef1-580b-45ad-b29c-fffc3d527202/@@download/attachment/ChIP-seq_Protocol_for_RNA-Binding_Proteins_ENCODE_Fu_lab_RuiXiao.pdf). In brief, prior to coupling with RBP antibodies, magnetic beads were equilibrated by washing with ChIP dilution buffer and blocked with glycogen, BSA and tRNA in ChIP dilution buffer. 10-20 million HepG2 and K562 cells were crosslinked in 1% formaldehyde diluted in 1xPBS for 20 minutes and then quenched by adding glycine. Cell nuclei were extracted by resuspending the cell pellet with cell lysis buffer with occasional inversion. Nucleus pellets resuspended in nuclear lysis buffer were sonicated with a Branson Sonifier cell disruptor. 95% of nuclear lysate was diluted to a final concentration of 1% triton X-100, 0.1% sodium deoxycholate and 1X proteinase inhibitor cocktail and was subjected to immunoprecipitation with antibody coupled beads; the other 5% of nuclear lysate was used as input chromatin. Stringent washes were performed before elution. Input and immunoprecipitated chromatin DNAs were recovered by decrosslinking, RNase A digestion, proteinase K treatment, phenol/chloroform extraction and precipitation with ethanol. Library construction was performed using the ChIP-seq Sample Prep Kit (Illumina). DNA Libraries between 200-400 bp were gel purified, quantified with Qubit and sequenced on the Illumina HiSeq 2000/2500. All RBP ChIP-seq experiments were performed in duplicate. Antibodies used in RBP ChIP-seq experiments were validated by immunoprecipitation and shRNA/CRISPR knockdown according to ENCODE RBP antibody characterization guidelines.

### ChIP-seq - data processing

RBP ChIP-seq datasets used in this study were processed by ENCODE Data Coordinating Center with the same uniform processing pipelines previously described for transcription factor ChIP-seq (https://www.encodeproject.org/chip-seq/transcription_factor/). After removing low quality and PCR duplicate reads, peaks were identified with SPP and reproducible peaks across biological replicates were identified with the IDR pipeline to yield 2 sets (optimal and conservative) of peaks at IDR threshold=0.05 ^66^. Data reproducibility was assayed using the IDR pipeline, requiring ‘pass’ or ‘borderline’ as previously described for ChIP-seq analysis^66^. Generally, > 10 million usable reads from each replicate were required; however, a limited set of datasets were exempted if manual inspection indicated significant reproducible signal at lower read depths. 5 datasets that showed reproducible signal but less than 200 reproducible peaks in the ‘optimal’ set were released at the ENCODE DCC but not included in further analysis (Supplementary Data 7).

### Integrated Analysis

#### Saturation Analysis

Saturation analysis of eCLIP and KD/RNA-seq data was performed by randomly shuffling the order of datasets 100 times, subsampling 1 through all datasets, and calculating the desired metrics. Gene level saturation analysis of RBP binding was calculated first by taking all unique genes that were bound by an IDR filtered peak in an eCLIP experiment. Then, each eCLIP experiment was iteratively added to the previous experiment, counting only unique genes in any experiment. Saturation analysis of differentially expression genes from KD/RNA-seq was similarly performed, based on differentially expressed genes identified with DESeq2. Genes were identified as differentially expressed if they had an adjusted p-value < 0.05 between knockdown and control. Alternative versions of this analysis used (Extended Data Fig. 8a) all genes, (Extended Data Fig. 8b) only genes with TPM > 1 in HepG2 and K562, or (Extended Data Fig. 8c) only genes with TPM > 1 in either HepG2 or K562, using average gene-level expression from two rRNA-depleted RNA-seq experiments in HepG2 (ENCODE accession ENCFF533XPJ, ENCFF321JIT) and K562 (ENCFF286GLL, ENCFF986DBN). The set of differentially expressed and bound genes was determined by taking all genes differentially expressed upon RBP KD that contained at least one IDR-filtered peak in the corresponding eCLIP experiment in the same cell type.

Differentially spliced events were defined as those meeting p-value < 0.05, FDR < 0.1, and change in Percent Spliced In (|∆Ψ|) > 0.05 from rMATS analysis (described above). The number of unique events was defined as the number of non-overlapping events upon combining all experiments for a given sampling. A differentially spliced event was considered bound if for any RBP in which the event was differentially included upon KD, there was an eCLIP peak for the same RBP in the same cell type between the start of the upstream flanking exon and the end of the downstream flanking exon for cassette exons and mutually exclusive exons, start of the upstream flanking exon and end of the common exon region for A3SS, start of the common exon and end of the common exon region for A5SS, and start of the upstream and stop of the downstream exons for retained introns).

To perform saturation of transcript regions, the highest expressed transcript for each gene was first identified using transcript-level quantitations from the same rRNA-depleted RNA-seq experiments described above. The following regions were then identified: the entire unspliced transcript (pre-mRNA), all exons (exon), 5’ untranslated regions (5’ UTR), coding sequence (CDS), 3' untranslated regions (3’UTR), all introns (intron), 100nt intronic regions flanking the 5' and 3' splice sites (splice site), proximal intronic regions extending from 100nt to 500nt from the 5' and 3' splice site (prox. intron), and distal intronic regions extending from 500nt and beyond from the 5' and 3' splice sites. Saturation calculations were then performed as described above for all genes (Extended Data Fig. 8b and Extended Data Fig. 8g-i) or only genes with TPM > 1 in both K562 and HepG2 (Fig. 2g and Extended Data Fig. 8f), and plotted as either the total number of bases covered (Extended Data Fig. 8e-f), or the fraction of covered bases divided by the total number of bases in that annotation across all genes (Fig. 2g). The ratio of bases covered was calculated by dividing the number of bases covered in subsampling of N+1 datasets divided by the number covered in subsampling N datasets.

Analysis of the fold-increase between one and two datasets (Extended Data Fig. 8i) was determined by first taking all 73 RBPs profiled in both HepG2 and K562, and calculating the fold-increase in covered bases by considering 146 comparisons including HepG2 followed by K562 and K562 followed by HepG2. Then, for each of the 146 comparisons, 10 random other datasets were chosen from the same cell type, and for each of the 10 the fold-increase in covered bases from adding that dataset to the first was calculated.

To compare the fold-increase between profiling new RBPs in additional cell lines, eCLIP datasets profiling RBFOX2, IGF2BP1, IGF2BP2, and IGF2BP3 in H9 human embryonic stem cells were obtained from the Gene Expression Omnibus (GSE78509)^76^, and added as the 224th dataset. These were compared against profiling a new RBP in K562 or HepG2 (calculated by adding each of the 150 profiled RBPs as the 222nd (if it was profiled in both cell types) or 223rd (if it was profiled in only one cell type) datasets for other RBPs), or a profiled RBP done in second cell type (calculated by sampling 222 datasets and adding the 223rd).

### Preservation of RBP regulation across cell types

To consider binding across cell types, first the highest expressed transcript for each gene was identified using transcript-level quantitations from the same rRNA-depleted RNA-seq experiments described above and used as representative for that gene. Next, genes were categorized based on their absolute fold-difference (FD) between K562 and HepG2: unchanged (FD ≤1.2), weakly (1.2 < FD ≤ 2), moderately (2 < FD ≤ 5) or strongly (FD > 5) differential (for each, requiring TPM ≥ 1 in both K562 and HepG2), cell-type specific genes (TPM < 0.1 in one cell type and TPM ≥ 1 in the other), or Other (all other genes in GENCODE v19). Peaks were then categorized based upon the expression change of their associated gene (Extended Data Fig. 8j).

Analysis of preservation of binding across cell types was considered in three ways. First, for each peak identified in one cell type, the fold-enrichment for that region in the other cell type was calculated and considered for each gene type (as in Fig. 2h). Two groups of peaks were then identified: those that were ≥ 4-fold enriched in the other cell type, and those that were not enriched in the other cell type. The fraction of peaks associated with a gene class that were either ≥ 4-fold or not enriched were then considered for each gene class separately (Fig. 2i). Second, the set of peaks ≥4-fold enriched (and the set not enriched) was compiled across all genes, and the fraction associated with each gene class were then reported (Extended Data Fig. 8l). Finally, peak overlap between cell types (Extended Data Fig. 8k) was calculated by determining the fraction of IDR peaks identified in one cell type that overlap (requiring at least 1nt overlap) IDR peaks identified in the second cell type. For all comparisons, significance between groups was determined by Kolmogorov-Smirnov test

### Motif comparisons between RBNS and eCLIP

eCLIP 6mer Z-scores in Fig. 3b were calculated as previously described^77^. Briefly, peaks and a shuffled background set of peaks that preserves the region of binding (3’UTR, 5’UTR, CDS, exon, proximal and distal intron) were generated. EMBOSS compseq [http://structure.usc.edu/emboss/compseq.html] was used on these two peak sets and the Z-scores of the difference between real and background 6mer frequencies was calculated.

To produce eCLIP logos in a similar manner for comparison with RBNS logos, an analogous procedure was carried out on the eCLIP peak sequences (for this analysis, eCLIP peaks with at least 2-fold enrichment were used): the two halves of the RBNS pulldown read set were replaced with the two eCLIP replicate peak sequence sets (each peak was extended 50 nt upstream of its 5’ end as some RBPs have motif enrichments symmetrically around or only upstream of the peak starts), and the input RBNS sequences were replaced by random regions within the same gene as each peak that preserved peak length and transcript region (5’ and 3’ UTR peaks were chosen randomly within that region; intronic and CDS peaks were shuffled to a position within the same gene that preserved the peak start’s distance to the closest intron/exon boundary to match sequence biases resulting from CDS and splicing constraints). The enrichment Z-score threshold for 5mers included in eCLIP logos was 2.8, as this threshold produced eCLIP logos containing the most similar number of 5mers to that of the Z≥3 5mer RBNS logos. Each eCLIP motif logo was filtered to include only 5mers that occurred in both of the corresponding eCLIP replicate logos. eCLIP motif logos were made separately for all eCLIP peaks, only 3’UTR peaks, only CDS peaks, and only intronic peaks, with the eCLIP logo of those 4 (or 8 if CLIP was performed in both cell types) with highest similarity score to the RBNS logo shown in Fig. 3a, where the similarity score was the same as previously described to cluster RBNS logos (eCLIP logos for all transcript regions shown in Extended Data Fig. 3a). To determine significance of overlap between RBNS and eCLIP, a hypergeometric test was performed with 5mers in all RBNS logos, eCLIP logo 5mers (for peaks in the region with highest similarity score to the RBNS logo), and 5mers in their intersection, relative to the background of all 1,024 5mers; overlap was deemed significant if P<0.05. The top ‘eCLIP-only’ logo in each region was the highest eCLIP logo, if any, comprised of 5mers that had no overlap with any RBNS Z≥3 5mers (always using at least the top 10 RBNS 5mers if there were fewer than 10 with Z≥3).

All eCLIP/RBNS comparisons were for the same RBP with the following exceptions in which the eCLIP RBP was compared to a closely related RBNS protein: KHDRBS2 eCLIP versus KHDRBS1 RBNS; PABPN1 eCLIP versus PABPN1L RBNS; PTBP1 eCLIP versus PTBP3 RBNS; PUM2 versus PUM1 RBNS; and RBM15 versus RBM15B RBNS.

### Splicing regulatory effects of RBNS+ and RBNS− eCLIP peaks

To assess the splicing regulatory effects of RBNS+ and RBNS− eCLIP peaks for Fig. 3c, only rMATS SEs with a Ψ between 0.05 and 0.95 in at least one of the control or KD were considered for each RBP. Each eCLIP peak (extended 50 nt 5’ of the peak start) was first checked if it overlapped the SE, and if not then if it overlapped the upstream or downstream flanking 250 nt. To compare the magnitude of splicing changes upon KD for eCLIP+ versus eCLIP− SEs while minimizing the confounding factors of different wildtype host gene expression level and SE Ψ values among these two sets of SEs, a matched set of eCLIP− SEs was created by selecting for each eCLIP+ SE an SE in the same decile of wildtype gene expression and wildtype Ψ for each corresponding SE with an eCLIP peak. A CDF of the ∆Ψ changes upon KD was compared for the eCLIP+ versus eCLIP– SEs in each of the 6 SE direction/eCLIP region combinations ([included, excluded SE] × [peak over SE, upstream intron, downstream intron]), with significance P<0.05 for a one-sided Wilcoxon rank-sum test that |∆Ψ|_SE, peak_ > |∆Ψ|_SE, no peak_. If the eCLIP+ versus eCLIP− comparison was significant, the eCLIP peaks were divided into those that did and did not contain the top RBNS 5mer. The ∆Ψ values for all RBPs in each of the 6 SE direction/eCLIP regions were combined for comparison in Fig. 3c; see Extended Data. Fig. 4a for RBPs that were significant in each region (12 included/4 excluded upon KD, upstream intron eCLIP peak; 11 included/2 excluded upon KD, SE eCLIP peak; 7 included/7 excluded upon KD, downstream intron eCLIP peak). To assess eCLIP peaks with or without the top ‘eCLIP-only’ *k*mer, the top *5*mer from the aforementioned ‘eCLIP-only’ logo was used from the first region with an eCLIP-only logo among: all peaks; CDS peaks; intron peaks; and 3’UTR peaks (the more highly enriched 5mer if eCLIP was performed in both cell types). The resulting ‘eCLIP-only’ 5mers for Extended Data Fig. 4b were: CELF1 (CUCUC), EIF4G2 (GUGUG), EWSR1 (CGCGG); FUBP3 (UUGUU); FUS (GUGUG); HNRNPC (GUCGC); HNRNPK (UCCCC); HNRNPL (none); IGF2BP1 (GUGUG); IGF2BP2 (CGCCG); KHDRBS2: (none); KHSRP (none); PABPN1L (CGCGG); PCBP2 (CGGCG); PTBP3 (GAAGA); PUM2 (UUUUU); RBFOX2 (GGGGG); RBM22 (GGUAA); SFPQ (UCCGG); SRSF5 (CGGCG); SRSF9 (CUGGA); TAF15 (AGGGA); TARDBP (GAAGA); TIA1 (CGCCG); TRA2A (GAGGG).

### Overlaps between RBP binding and gene expression perturbation upon KD/RNA-seq

To increase sensitivity for gene expression analysis, significant binding was determined at the level of transcript regions (including 5’UTR, CDS, 3’UTR, and introns) instead of using peaks. To identify significant enrichment between binding and expression changes, genes with significantly enriched eCLIP signal at regions (p ≤ 0.00001 and log_2_(fold-enrichment) ≥ 4, as described above) were overlapped with the set of genes with significantly altered expression in KD/RNA-seq (adjusted p-value < .05 between knockdown and control from DEseq analysis). Enrichment was calculated separately for knockdown-increased and knockdown-decreased genes, with significance determined by Fisher Exact test (or Yates’ Chi-Square test if all observed and expected values were above 5). Comparisons with either knockdown-increased or knockdown-decreased genes from knockdown RNA-seq were only performed if more than 10 genes showed significant changes. To avoid biases due to RNA abundance, for each comparison of a region type with each eCLIP dataset a background set of genes was created by identifying all genes for which the region type (5’UTR, CDS, 3’UTR) had at least 10 reads in one of IP or input, at least 10 reads would be expected in the opposite (IP or input) dataset given the total number of usable reads. For cumulative distribution plots, genes were separated based on their eCLIP fold-enrichment in IP versus input for the indicated transcript region.

To perform TIA1 motif enrichment analysis, first the fold-enrichment of each 5mer was calculated by comparing the frequency in 3’UTRs of genes increased or decreased upon TIA1 knockdown in K562 or HepG2 with the frequency in a set of control genes upon knockdown (changed genes upon KD: DEseq adjusted P-val < 0.05 and |Fold-Change| > 1.5; control genes: DEseq P-val > 0.5 and |Fold-Change| < 1.1, subsetted to match the starting expression of changing genes upon KD). The top 15 5mers in TIA1 RBNS were then highlighted among in the ranked ordering of all 1,024 5mers. For positional analysis, a meta-3’UTR was created by normalizing all 3’UTRs to a 100nt window. For each normalized position, the frequency of the top 10 TIA1 RBNS 5mers was calculated for each of the up-regulated, down-regulated, and control gene sets. Significance at each position was determined by P < 0.05 in a binomial test comparing the number of up- or down-regulated genes that have one of the top 10 RBNS 5mers at that position under the null frequency that it is equal to the corresponding frequency observed in control genes.

### RBP binding correlation with knockdown-perturbed splicing (splicing maps)

RBP binding/splicing maps were generated using eCLIP normalized (reads per million) read densities overlapped with alternatively spliced (AS) regions from rMATS JunctionCountsOnly files from the same cell type. First, the set of differentially alternatively spliced events of the desired type (cassette/skipped exons (SE), alternative 5’ splice site (A5SS), or alternative 3’ splice site (A3SS) events were identified (Extended Data Fig. 18a), requiring rMATS p-value < 0.05, FDR < 0.1, and |∆Ψ| > 0.05 in knockdown versus control RNA-seq. To eliminate potential double counting of CLIP densities, overlapping AS events were additionally filtered to choose only the events containing the highest average inclusion junction count (IJC) among all replicates (using the bedtools v2.26 command merge (-o collapse -c 4) and pybedtools 0.7.9).

Next, for each splicing event, per-position input probability densities were subtracted from IP probability densities to attain position-level enrichment or depletion, for regions extending 50nt into each exon and 300nt into each intron composing the event. Subtracted read densities were then normalized to sum to 1 across each event in order to equally weigh each event, creating tracks referred to as ‘Normalized eCLIP enrichment’ (Extended Data Fig. 18b). For shorter exons (<100 nt) and introns (<600nt), densities were only counted until the boundary of the neighboring feature. Skipped exon (SE) maps were plotted using eCLIP densities overlapping the following 4 regions around AS events: 3' end of the upstream exon, 5' end of the cassette, 3' end of the cassette, and 5' end of the downstream exon. A lternative 3' splice site (A3SS) maps were defined with three regions: 3' end of the upstream exon, 5' end of the longer transcript, and the 5' end of the shorter transcript. Alternative 5' splice site (A5SS) maps were defined with three regions: 3' end of the shorter transcript, 3' end of the longer transcript, and the 5' end of the downstream exon.

Plots of eCLIP signal enrichment (referred to as ‘splicing maps’) were then created by calculating the mean and standard error of the mean over all events after removing the highest (2.5%) and lowest (2.5%) outlying signal at each position, referred to as ‘Average eCLIP enrichment’ (Extended Data Fig. 18c). Splicing maps were only considered for RBPs with 100 or more altered cassette exon events, or 50 or more alternative 5’ or 3’ splice site events, considering knockdown-included and knockdown-excluded events separately. Out of a total of 203 pairings of eCLIP and knockdown/RNA-seq in the same cell type (covering 139 RBPs), this left 92 pairings (72 RBPs) for cassette exons, 27 pairings (22 RBPs) for A3SS, and 20 pairings (18 RBPs) for A5SS. As a background reference for cassette exon comparisons, sets of 1,805 (HepG2) and 2,222 (K562) ‘native’ cassette exons were identified which had 0.05 < Ψ < 0.95 in at least half of control shRNA RNA-seq datasets for that cell type. Similar sets of 202 (K562) and 159 (HepG2) native alternative 5’ splice site and 389 (K562) and 352 (HepG2) native alternative 3’ splice site events were identified that had 0.05 < Ψ < 0.95 in at least half of control shRNA RNA-seq datasets for that cell type. RBP-responsive event eCLIP enrichment was then calculated as eCLIP signal enrichment at RBP-regulated events minus eCLIP signal enrichment at native control events, referred to as ‘Enrichment relative to control events’ (Extended Data Fig. 18d). To calculate significance, 1000 random samplings were performed from the native cassette exon set using the number of events in the knockdown-included (or excluded), and significance was set as being either lower than the 0.5^th^ or higher than the 99.5^th^ percentile for each position.

Correlation between splicing maps was defined as the Pearson correlation (R) between a vector containing both included-upon knockdown and excluded-upon knockdown RBP-responsive event eCLIP enrichment for each RBP. If an RBP had less than the minimum required number of events (100 for cassette exons or 50 for alternative 5’ or 3' splice site events) for either knockdown-included or knockdown-excluded events, the correlation was only calculated using the other event type.

To generate cross-RBP splicing maps, the above approach was modified by taking the set of differentially included (or excluded) cassette exons identified in knockdown of RBP A and calculating the eCLIP splicing map separately for every other RBP within the same binding class (determined in Fig. 2b) as RBP A, including the normalization against a background of eCLIP signal for native SE events (as shown for HNRNPC knockdown-included, RBFOX2 knockdown-excluded, and TIA1 knockdown-included cassette exons in Extended Data Fig. 20b,c,e respectively). The average across all RBPs was then used to calculate the average cross-RBP enrichment (Extended Data Fig. 20a).

To calculate the number of RBPs bound per exon, the set of spliceosomal RBPs was taken from manual annotation of RBP functions (described above and listed in Supplementary Data 1). The number of reproducible (IDR) peaks at each position relative to splice sites was summed across all RBPs and divided by the total number of cassette or constitutive exons respectively.

### Comparison of DNA- and RNA-binding properties of RBPs

For integrative analyses, DNaseI HS data (http://genome.ucsc.edu/cgi-bin/hgFileUi?db=hg19&g=wgEncodeOpenChromSynth), histone modifications by ChIP-seq from ENCODE/ Broad Institute (http://genome.ucsc.edu/cgi-bin/hgFileUi?db=hg19&g=wgEncodeBroadHistone) and eCLIP-seq data from ENCODE (https://www.encodeproject.org) were downloaded and compared with RBP ChIP-seq data.

To explore the possibility that some RBP chromatin association events might be coupled with their direct RNA binding activities in cells, RNA binding peaks were compared with DNA binding signals as assayed by ChIP-seq to quantify enrichment. Only eCLIP peaks in gene body regions (excluding promoter and terminator regions, defined as the 1kb surrounding regions of TSS and TTS) were considered. The ChIP-seq signals were calculated for each eCLIP peak, together with surrounding regions that are 10 times the length of eCLIP peak on each side. Wilcoxon rank-sum tests were then performed to see whether ChIP-seq signal were enriched at the middle third regions.

To see whether those differentially-expressed genes after RBP knockdown were enriched in RBP binding at chromatin level, equal numbers of genes with similar expression level either with or without binding to the TSS region were randomly sampled, the number of differentially-expressed genes after knockdown of the RBP were counted (fold change > 1.5 or < 2/3, adjusted p-value <0.05 by DESeq2), and one-tailed Fisher’s exact tests were then performed to test the dependence of RBP binding and differential expression. Odds ratio was defined as (a/b) / (c/d), where a = the number of genes with RBP ChIP-seq peaks and differential expression (or splicing) upon RBP knockdown, b = genes with RBP ChIP-seq peaks but no differential expression, c = genes without ChIP-seq peaks but with differential expression, and d = genes without ChIP-seq peaks or differential expression. The above procedure was performed 100 times to give the distribution of the odds ratio. A significant dependence was defined when the null hypothesis was rejected at level of 0.05 for at least 95 times. The correlation between RBP association and genes with regulated alternative splicing events (A3SS, A5SS, RI, MXE and SE events) were investigated similarly.

### Analysis of RBP regulatory features in subcellular space

Localization annotations and calculation of nuclear versus cytoplasmic ratio were generated from immunofluorescence imaging as described above. “Nuclear RBPs” were defined as those with nuclear/cytoplasmic ratio ≥ 2, and “Cytoplasmic RBPs” were defined as those with nuclear / cytoplasmic ratio ≤ 0.5. Spliced reads were defined as reads mapping across an annotated GENCODE v19 splice junction (extending at least 10 bases into each exon) and unspliced reads were defined as reads that overlapped an exon-intron junction (extending at least 10 bases into both the exon and intron regions). Significance between groups was determined by Wilcoxon rank sum test. Prediction of RNA secondary structure was performed using the RNAfold webserver (http://rna.tbi.univie.ac.at//cgi-bin/RNAWebSuite/RNAfold.cgi)^78^ with default parameters. Shown is the MFE secondary structure prediction.

### RBP expression in tissues

Tissue specificity was measured as the entropy deviation from a uniform distribution among all tissues as in^1^. For each RBP, the log_2_(TPM+1) was calculated for each of the 42 samples (HepG2, K562, and 40 tissues profiled by the GTEx consortium^79^), and the tissue specificity was computed as the difference between the logarithm of the total number of samples (*N*=42) and the Shannon entropy of the expression values for an RBP:

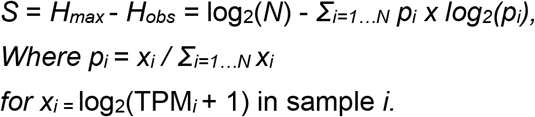

The data used for the analyses were obtained from dbGaP accession number phs000424.v2.p1 in Jan. 2015. TPMs were measured using kallisto^80^ on the following samples: Adipose-Subcutaneous: SRR1081567; AdrenalGland: SRR1120913; Artery-Tibial: SRR817094; Bladder: SRR1086236; Brain-Amygdala: SRR1085015; Brain-AnteriorCingulateCortex: SRR814989; Brain-CaudateBasalGanglia: SRR657731; Brain-CerebellarHemisphere: SRR1098519; Brain-Cerebellum: SRR627299; Brain-Cortex: SRR816770; Brain-FrontalCortex: SRR657777; Brain-Hippocampus: SRR614814; Brain-Hypothalamus: SRR661179; Brain-NucleusAccumben: SRR602808; Brain-SpinalCord: SRR613807; Brain-SubstantiaNigra: SRR662138; Breast-MammaryTissue: SRR1084674; Cervix: SRR1096057; Colon: SRR1091524; Esophagus: SRR1085211; FallopianTube: SRR1082520; Heart-LeftVentricle: SRR815517; Kidney-Cortex: SRR809943; Liver: SRR1090556; Lung: SRR1081283; MinorSalivaryGland: SRR1081589; Muscle-Skeletal: SRR820907; Nerve-Tibial: SRR612911; Ovary: SRR1102005; Pancreas: SRR1081259; Pituitary: SRR1077968; Prostate: SRR1099402; Skin: SRR807775; SmallIntestine: SRR1093314; Spleen: SRR1085087; Stomach: SRR814268; Testis: SRR1081449; Thyroid: SRR808886; Uterus: SRR820026; Vagina: SRR1095599.

## Acknowledgments

The authors would like to thank Elise Feingold, Mike Pazin, Dan Gilchrist and Grace Xiao for helpful discussions, and Cricket Sloan, Jean Davidson, Eurie Hong, and Mike Cherry for assistance with data deposition and distribution to the public. This work was funded by the National Human Genome Research Institute ENCODE Project, contract U54HG007005, to BRG (principal investigator) and X-DF, CBB and GWY (co-principal investigators), U41HG009889 to BRG (PI), GWY (PI), CB (coPI) and EL (coPI), as well as a grant from the Fonds de Recherche du Québec-Santé to EL. ELVN is a Merck Fellow of the Damon Runyon Cancer Research Foundation (DRG-2172-13) and is supported by a K99 grant from the NHGRI (HG009530). GAP is supported by the NSF graduate research fellowship. XDF and GWY were partially supported by grants from the NIH (HG007005, NS075449).

## Author Contributions

Work on the paper was divided between data production and analysis. The analysts were ELVN, PF, GAP, XW, RX, JC, DD, LPBB, MOD, NJL, SS, BAY, BZ. The data producers were ELVN, XW, RX, SMB, NALC, DD, SO, BS, LZ, CB, JB, KG, CG-B, MH, NJL, HL, TBN, TP, IR, RS, AS, RW, ALL, and SA. Substantially larger contributions were made by the joint first authors. Overall project management was carried out by the senior authors X-DF, EL, CBB, BRG and GWY.

## Author Information

Data sets described here can be obtained from the ENCODE project website at http://www.encodeproject.org via accession numbers in Supplementary Data 2. ELVN and GWY are co-founders and consultants for Eclipse BioInnovations Inc. The terms of this arrangement have been reviewed and approved by the University of California, San Diego in accordance with its conflict of interest policies. The authors declare no other competing financial interests.

---

**Extended Data Figure 1.**
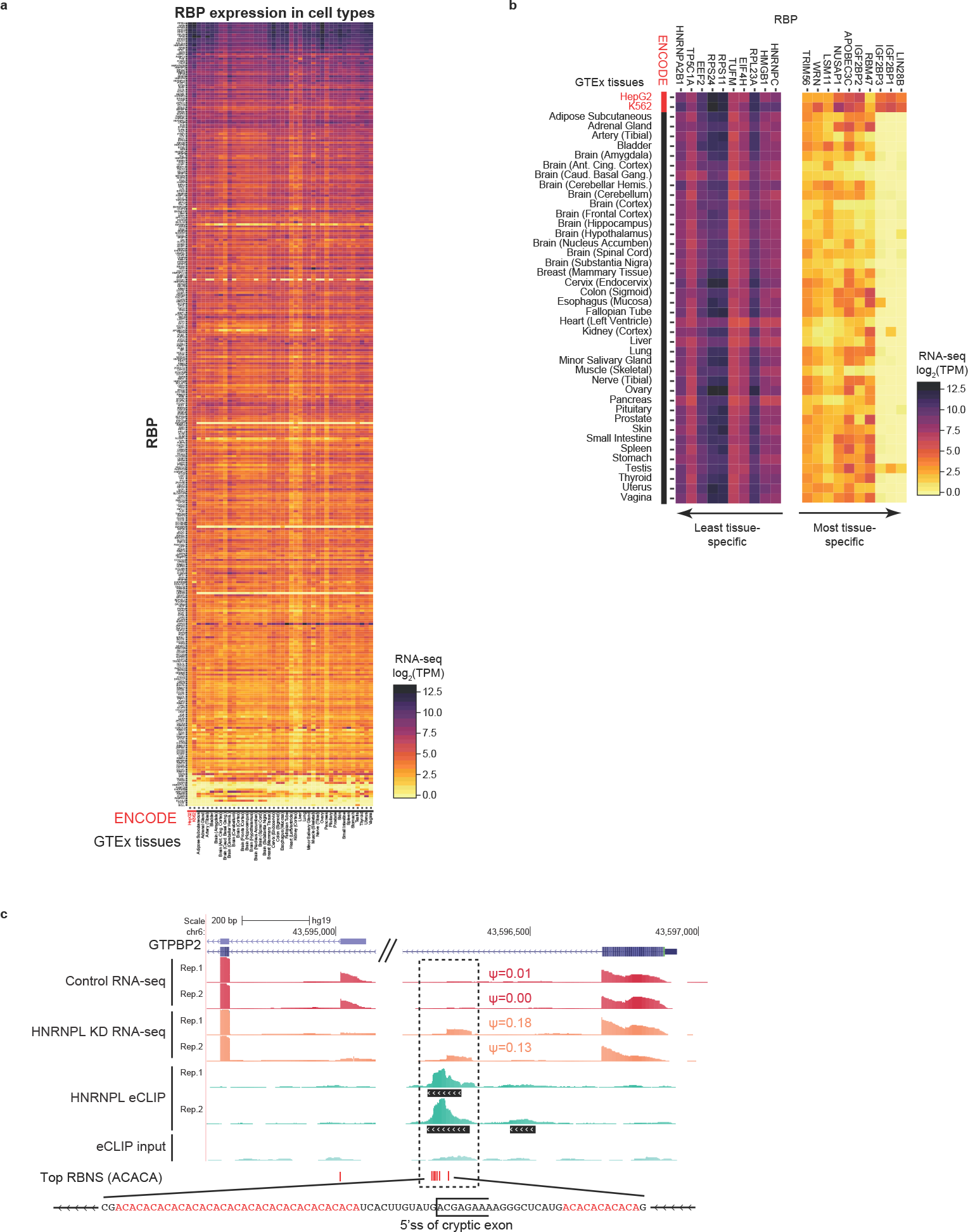
Expression of RBPs across tissues and cell types. (a) Expression of the 356 RBPs (in Transcripts Per Million) investigated in this study in ENCODE cell lines HepG2 and K562 as well as 40 human tissues measured by the GTEx project. RBPs sorted by decreasing expression in HepG2. (b) Expression of the 10 RBPs with the highest and lowest tissue-specificity across the two ENCODE cell lines and 40 human tissues. (c) Shown is RNA-seq read density (reads per million), eCLIP read density (reads per million), and RBNS motif presence proximal to a 73-nt cryptic exon expected to induce nonsense-mediated decay (NMD) of GTP Binding Protein 2 (a ribosome rescue factor whose loss induces neurodegeneration in certain genetic backgrounds^81^). eCLIP indicates that HNRNPL binds over the cryptic exon 5’ splice site in a sequence-specific manner to a region rich in the top RBNS 5mer, ACACA.

**Extended Data Figure 2.**
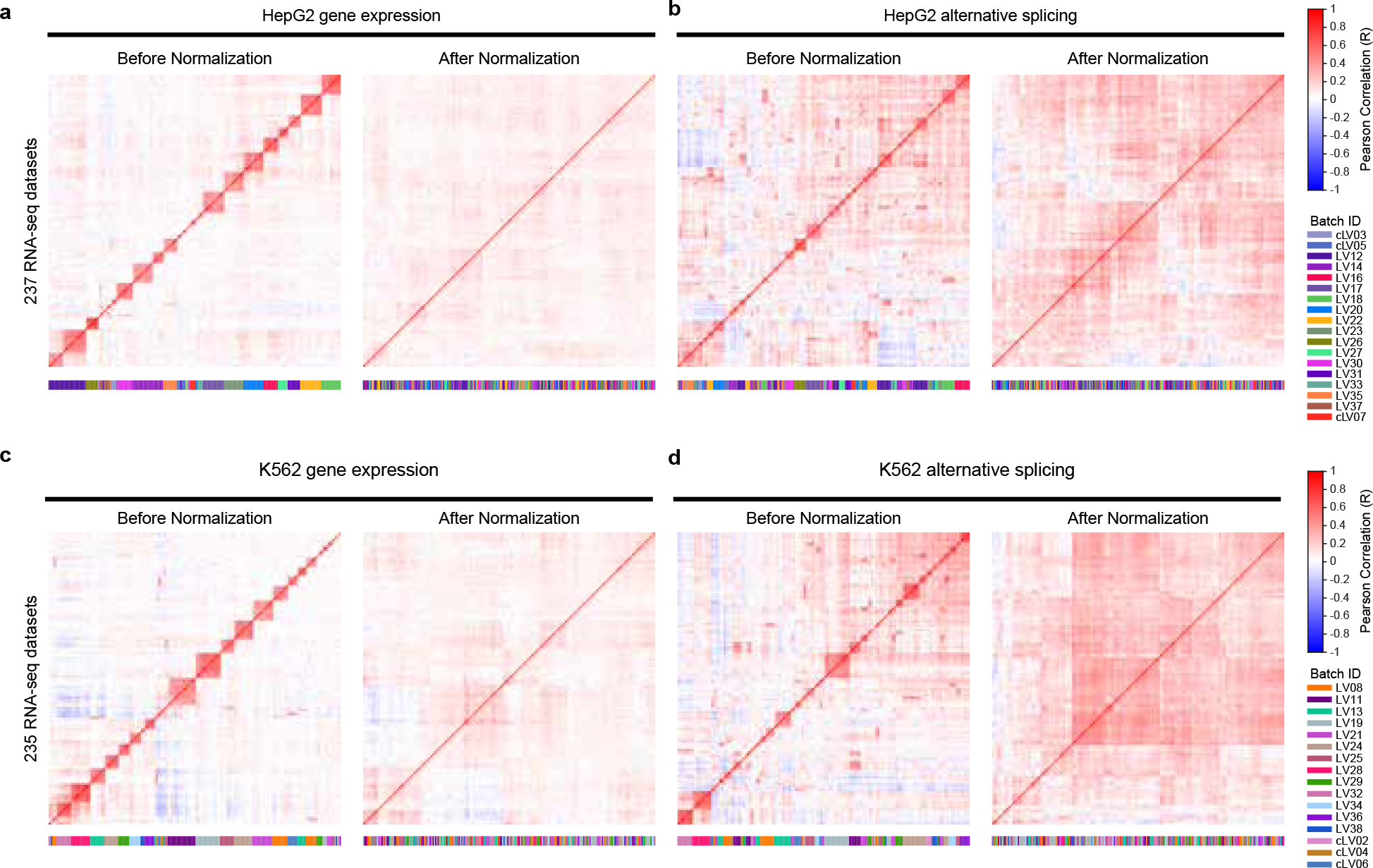
Batch correction of RBP knockdown RNA-seq datasets. (a-d) Heatmaps show Pearson correlation between all RNA-seq datasets before and after normalization to remove batch effects, followed by hierarchical clustering. Analysis was performed separately for (a-b) HepG2 and (c-d) K562 cell lines. (a,c) For gene expression, correlation was determined between gene expression fold-change values (log_2_) from comparison of (left) RBP knockdown versus paired control or (right) performing batch correction on all datasets (as described in Methods) followed by comparing RBP knockdown replicates versus a ‘virtual control’ defined as the average of all replicate 1 or replicate 2 control experiments respectively. (b,d) For splicing, correlation was calculated between change in exon inclusion values between (left) RBP knockdown and within-batch control experiments, and (right) RBP knockdown versus a ‘virtual control’ defined as the average of all replicate 1 or replicate 2 control samples respectively following batch correction of junction read counts as described in Methods. For all, colors below indicate experimental batches.

**Extended Data Figure 3.**
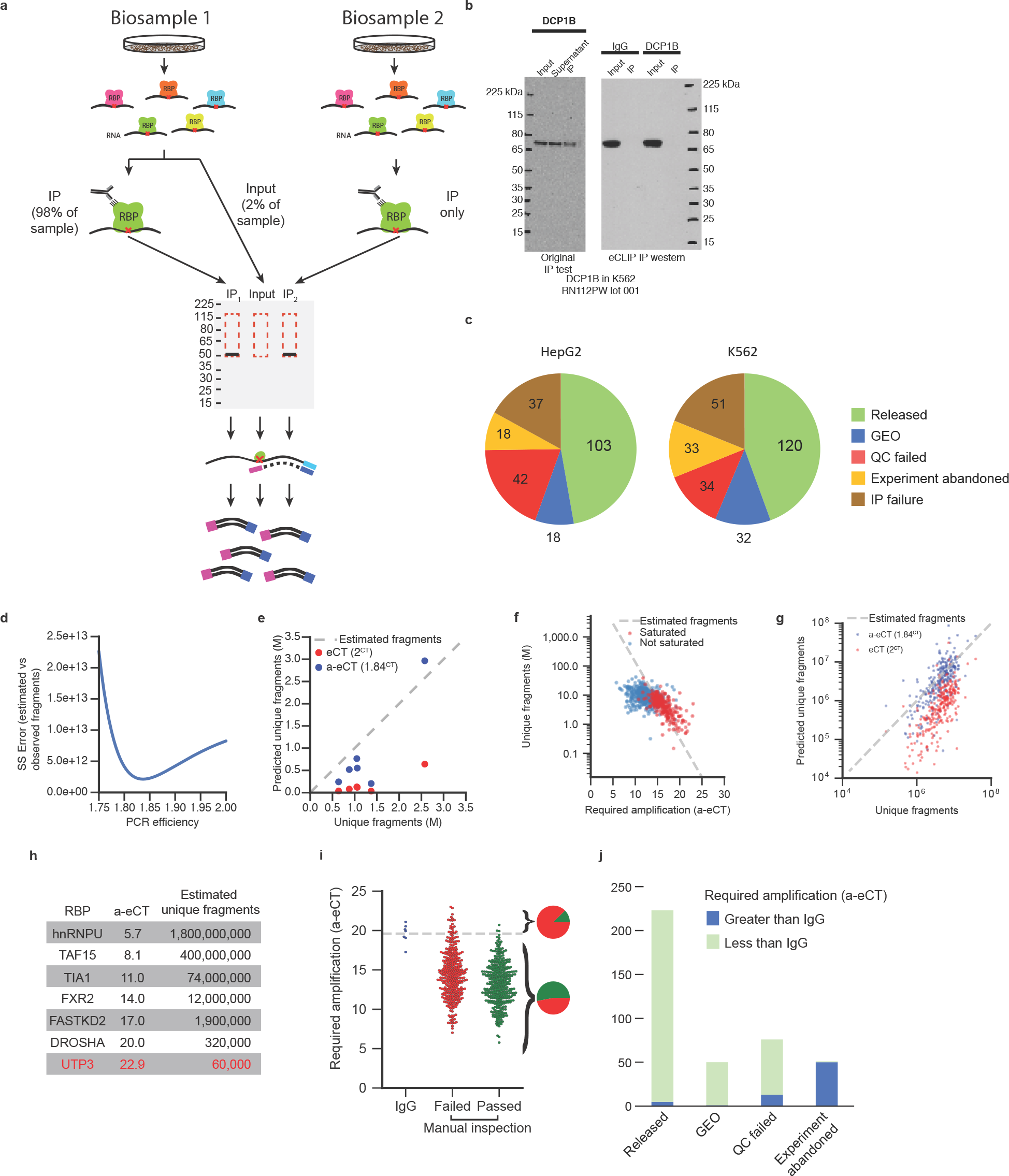
Experimental quality assessment of eCLIP assays. (a) Model of ENCODE eCLIP experiments. Inputs were taken by sampling 2% of one of the two biosamples prior to immunoprecipitation (IP). (b) Example IP-western image for DCP1B (left) during initial IP tests performed without enzymatic steps and (right) during eCLIP experiments. (c) Pie charts indicate the number of eCLIP experiments that fell into the following categories: failure to successfully immunoprecipitated during eCLIP (IP failure), failure to yield amplifiable library in less than 20 PCR cycles (Experiment abandoned), experiments which yielded immunoprecipitated library and were sequenced but failed quality assessment (QC failed), successful experiments which did not meet ENCODE standards but contained reproducible signal and have been released on the Gene Expression Omnibus (GEO), and successful experiments which met ENCODE standards and are available at the ENCODE Data Coordination Center (Released). (d) Plot indicates sum of squared error for varying PCR efficiency when comparing true observed number of unique molecules to estimated number of unique molecules for six highly saturated (>90% PCR duplicated) experiments. (e) Scatter plot of estimated unique molecules at two estimates of PCR efficiency, (red) 2 and (blue) 1.84 versus unique fragments obtained after sequencing for six highly saturated (>90% PCR duplicated) experiments. (f) Scatter plot indicates accurate-eCT (a-eCT) (see Methods) versus unique fragments observed (including non-PCR duplicate reads mapped either to unique genomic loci or repetitive elements, in millions of reads mapped) for all ENCODE eCLIP experiments. Non-saturated (<60% PCR duplicates) datasets are indicated in blue, and saturated (>60% PCR duplicates) datasets are indicated in red. Dashed line indicates the number of unique molecules expected based on a-eCT. (g) Scatter plot of estimated unique molecules at two estimates of PCR efficiency, (red) 2 and (blue) 1.84 versus unique fragments obtained after sequencing. Shown are 276 moderately saturated experiments (>60% PCR duplicated). (h) Representative RBPs are listed along with their a-eCT and corresponding estimate of the number of unique RNA molecules isolated in eCLIP. UTP3 (in red) did not pass quality control metrics. (i) Points indicate the a-eCT value of all ENCODE eCLIP experiments, separated into (blue) IgG controls, (red) datasets that failed manual quality assessment, and (green) datasets passing manual assessment. Dotted line indicates average a-eCT of IgG control experiments (19.6). (j) Bars indicate the distribution of eCLIP datasets (separated into classes as described in (c)) with respect to required amplification (a-eCT) relative to IgG controls.

**Extended Data Figure 4.**
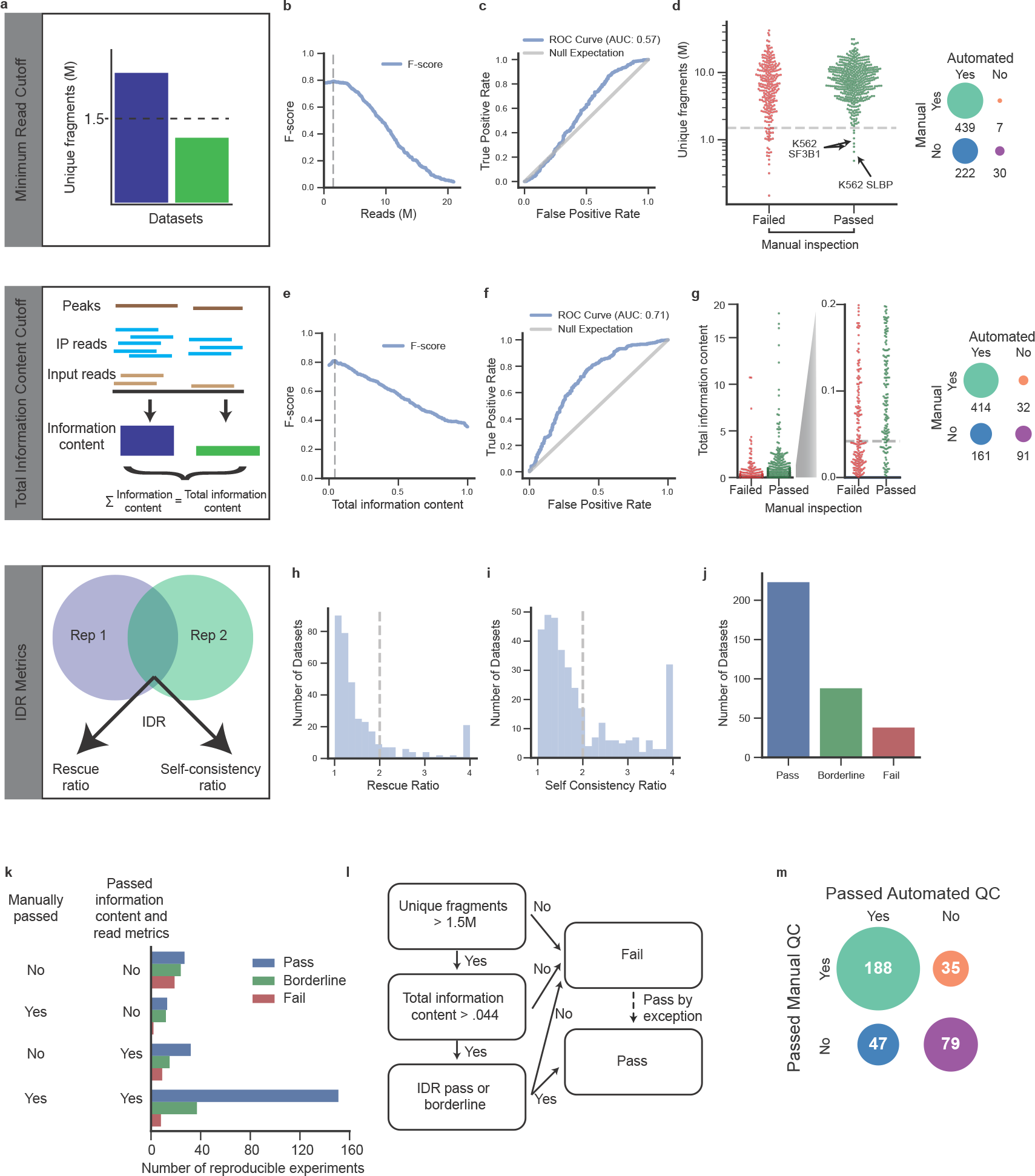
Quality metrics to assay eCLIP data quality and reproducibility. (a) Schematic of eCLIP data quality standards. (b) Plot indicates f-score for classification of datasets relative to manual quality assessment based on unique fragments present. Maximal classification of datasets was obtained at a cutoff of 1.5 million unique fragments. (c) ROC curve for classifying datasets based upon varying minimum unique fragment thresholds. (d) Swarm plot indicates number of unique fragments observed in each eCLIP dataset separated by (red) failing or (green) passing manual quality inspection. Dashed line indicates 1.5 million read quality threshold that maximizes predictive power on manual classification (as shown in (b)) and inset indicates confusion matrix for this threshold versus manual inspection. Three datasets judged to be high quality despite low unique fragment number are indicated. (e) Plot indicates f-score for classification of datasets based on the total information content in all significantly enriched peaks. Only datasets passing the unique fragment cutoff in (a-d) were considered. (f) ROC curve for classifying datasets based upon varying total information in peak cutoff. (g) Swarm plot indicates the total information content across all peaks in each eCLIP dataset that passes the unique fragment threshold in (d), separated by (red) failing or (green) passing manual quality inspection. Dashed line indicates the information content threshold that maximizes predictive power on manual classification (as shown in (e)) and inset indicates confusion matrix for this threshold versus manual inspection. (h) Bar plot indicates IDR rescue ratio for all ENCODE eCLIP experiments. (i) Bar plot indicates IDR self-consistency ratio for all ENCODE eCLIP experiments. Dashed line indicates a cutoff of 2 previously used for ChIP-seq analysis. (j) Bars indicate the number of ENCODE eCLIP experiments that either (pass, in blue) pass both rescue ratio and self-consistency ratio, (borderline, in green) passed just one of the two tests, or (fail, in red) failed both tests. (k) Bar chart indicates the count of all ENCODE experiments that pass or fail manual or automated QC approaches, broken into three groups based on their IDR thresholding metric status: (blue) passed, (green) borderline, and (red) failed. (l) Schematic detailing final recommended quality assessment decision flowchart. (m) Confusion matrix of final classification scheme versus manual quality assessment.

**Extended Data Figure 5.**
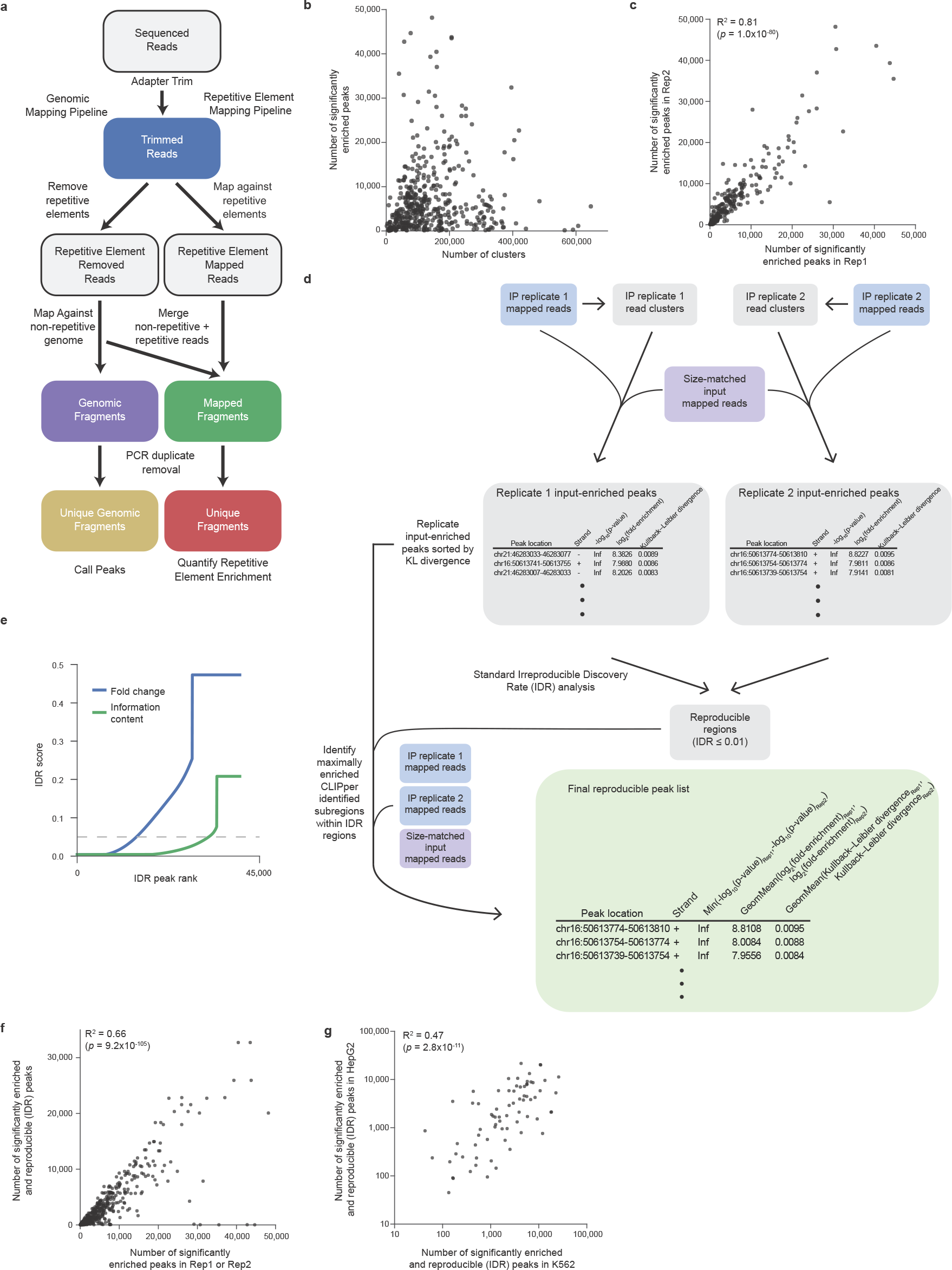
Identification of reproducible eCLIP peaks. (a) Schematic of eCLIP processing for both unique genomic mapping and repetitive element mapping. (b) Points indicate (x-axis) the number of CLIPper-identified clusters versus (y-axis) the number of significantly enriched peaks (fold-enrichment ≥ 8 and *p* ≤ 0.001) identified for each eCLIP experimental replicate. (c) Points indicate the number of significantly enriched peaks (fold-enrichment ≥ 8 and *p* ≤ 0.001) identified in replicate 1 versus replicate 2 for each of 223 high-quality eCLIP experiments. (d) Schematic of adaption of Irreproducible Discovery Rate (IDR) analysis to identification of reproducible eCLIP peaks. First, input-normalized clusters are identified separately for two biological replicates. Next, these peaks are ranked by relative information content, defined as 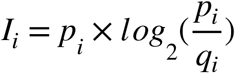, for proportion of IP reads within peak *i* represented by *p*_*i*_ and fraction of input reads within the peak as *q*_*i*_. Next, standard IDR analysis is performed on the ranked peak lists to identify reproducible regions at IDR cutoff of 0.01. Next, we considered all CLIPper-identified subregions within these IDR regions, and calculated the fold-enrichment in IP versus input for each subregion in each replicate. Subregions were ranked by the geometric mean of fold-enrichment between the two replicates, and the set of non-overlapping subregions that were significantly enriched (*p* ≤ 0.001 in both replicates) with geometric mean of fold-enrichment ≥ 8 in both replicates were obtained as the set of reproducible peaks (e) Plot indicates each peak ranked by IDR score, when IDR score is calculated by ranking peaks based on (blue) fold-enrichment above input or (green) information content. (f) Points indicate the number of significantly enriched peaks (fold-enrichment ≥ 8 and *p* ≤ 0.001) identified in each of replicate 1 and replicate 2 versus the number of reproducible peaks identified from IDR analysis (as shown in (b)). (g) Points indicate the number of significant and reproducible peaks identified in (x-axis) K562 versus (y-axis) HepG2, for all RBPs with eCLIP in both cell types.

**Extended Data Figure 6.**
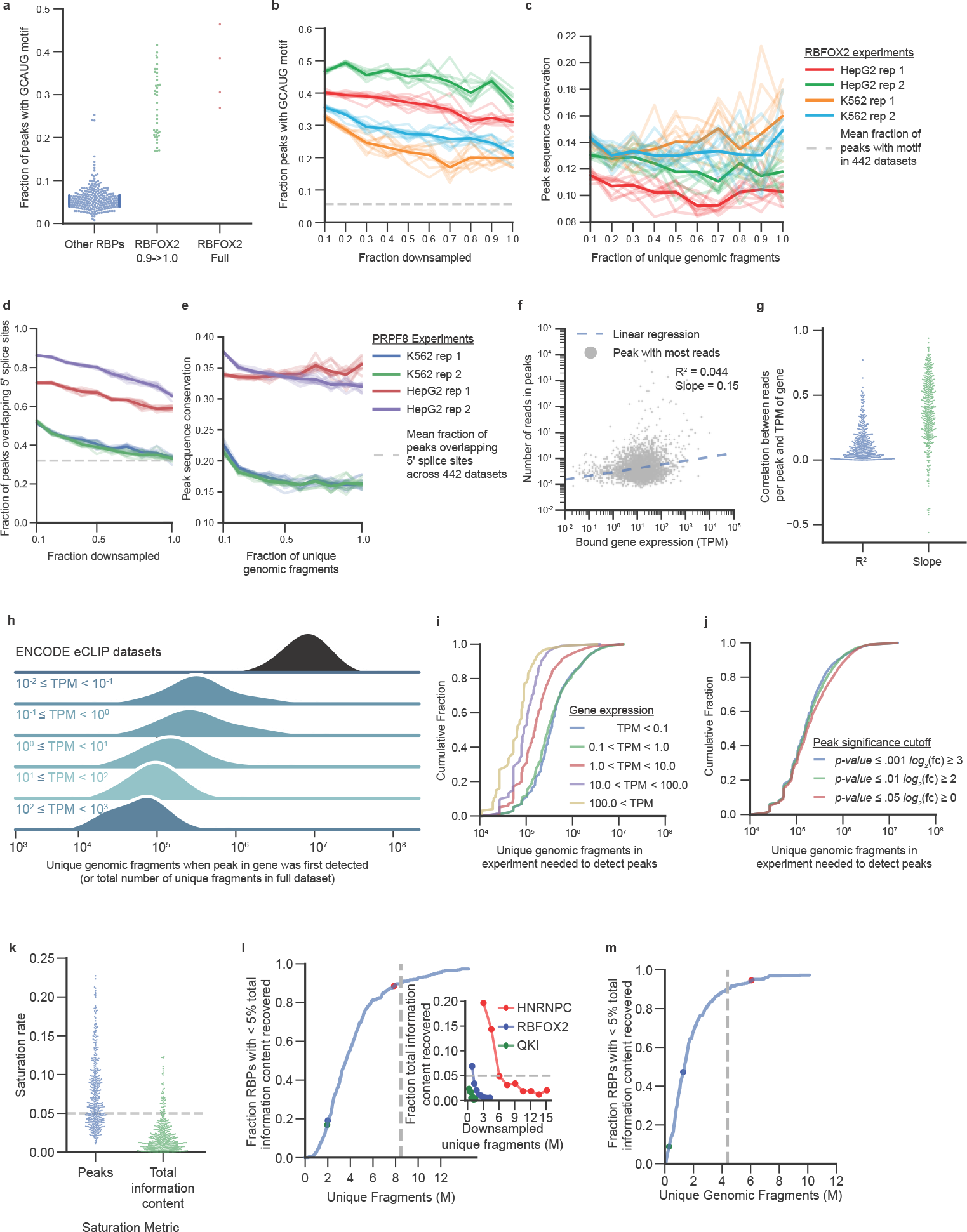
Analysis of eCLIP signal detection based on sequencing depth. (a) Points indicate the fraction of significant peaks that contain a GCAUG motif. Shown are (blue) all RBPs except RBFOX2, (green) peaks newly identified when comparing the 90% subsampled to full RBFOX2 dataset, and (red) the full RBFOX2 dataset. (b) Plot indicates the fraction of peaks containing a GCAUG motif for peaks identified in a series of subsamples of eCLIP unique genomic fragments for RBFOX2 in HepG2 and K562. Shown are RBFOX2 HepG2 (red) replicate 1 and (green) replicate 2, and RBFOX2 K562 (orange) replicate 1 and (blue) replicate 2. The dashed grey line indicates the mean fraction of GCAUG-containing peaks observed across all released eCLIP datasets. (c) Plot indicates mean mammalian phastons conservation for all peaks newly discovered in each downsampled subsample for RBFOX2 eCLIP in HepG2 (red) replicate 1 and (green) replicate 2 and K562 (orange) replicate 1 and (blue) replicate 2. (d-e) Downsampling analysis for PRPF8 eCLIP in HepG2 (blue) replicate 1 and (green)replicate 2, and K562 (red) replicate 1 and (purple) replicate 2. (d) Plot indicates the fraction of peaks newly discovered in each downsampled subsample that overlap the 5’ splice site. The dashed grey line indicates mean fraction overlap with 5’ splice sites for all 512 non-PRPF8 released ENCODE datasets. (e) Lines indicate the average conservation for newly discovered peaks at the indicated downsampling fraction. (f) One point for each gene indicates the TPM (Transcripts Per Million reads) of the gene (x-axis) and the number of reads (normalized by peak size) in the peak with the highest number of reads for RBFOX2 in HepG2. Dashed line indicates simple linear regression. (g) Points indicate the Pearson correlation coefficient (*R*^2^) (blue) and slope (green) for the linear regression between gene TPM and maximum peak read density for all released ENCODE eCLIP experiments. Each point represents an individual dataset as shown in (f). (h) Joy plot indicates (top) the distribution of unique genomic fragment values for released ENCODE eCLIP experiments, versus (bottom) the distribution of total eCLIP unique genomic fragments in the downsampled subsample where the first peak was identified in each gene, separated into bins by gene TPM. (i) Cumulative distribution function plot indicates the number of reads needed to first detect peaks for the set of genes in indicated bins separated by gene TPM: (blue) TPM < .01, (green) . 01 ≤ TPM < 1.0, (red) 1.0 ≤ TPM < 10.0, (purple) 10.0 ≤ TPM < 100.0, and (gold) 100.0 ≤ TPM. (j) Plots indicate the cumulative fraction of genes with peaks discovered at given experimental sequencing depth, for the indicated cutoffs for peak enrichment in IP versus input (p-value ≤ . 001 and fold-enrichment ≥ 8 (blue), p-value ≤ .01 and fold-enrichment ≥ 4 (green), p-value ≤ .05 and fold-enrichment ≥ 0 (red). (k) Points indicate saturation rate for peak or total information content between the 90% subsampled fraction retained and 100% (full dataset) for all 223 high quality ENCODE eCLIP experiments. Grey dashed line is 5% saturation cutoff. (l) (right) Lines show percent of additional information recovered when adding 10% additional reads for (red) HNRNPC, (blue) RBFOX2, and (green) QKI in HepG2, with number of unique (non-PCR duplicate) fragments indicated by the x-axis. Dotted line indicates the ‘saturation’ point at which less that 5% additional information is gained. (left) Cumulative fraction plot indicates the distribution of unique fragments when each eCLIP dataset reaches saturation. Colored points indicate depth of sequencing when HNRNPC, RBFOX2 and QKI saturate. As in (l), but points are now plotted relative to unique genomic-mapped non-PCR duplicate fragments only.

**Extended Data Figure 7.**
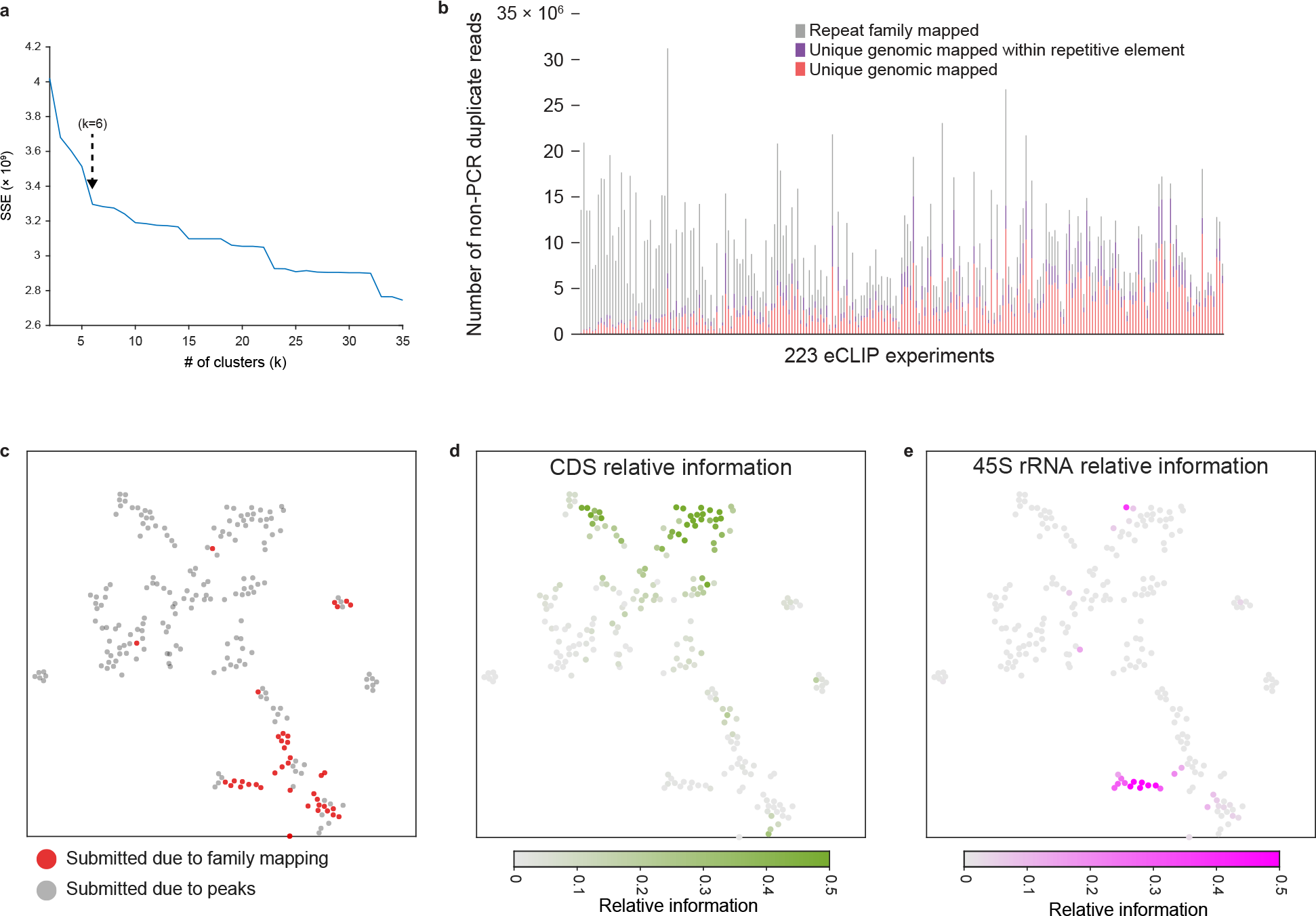
Integrated analysis of 223 eCLIP datasets identifies RBP clusters based on binding patterns. (a) Plot shows the effect of cluster number on hierarchical clustering on the Euclidean distance between RBPs for the fraction of peaks overlapping each of the RNA region types as shown in Figure 2b. For each number of clusters *k* between 2 to 35, the sum of squared error was calculated between the number of peaks annotated for each region versus the mean of all RBPs in that RBP’s cluster and summed across all RBPs. An inflection point was identified at *k*=6 (indicated). (b) Stacked bars indicate the number of reads from replicate 1 of all 223 eCLIP experiments, separated by whether they map (red) uniquely to the genome, (purple) uniquely to the genome but within a repetitive element identified by RepeatMasker, or (grey) to repetitive element families. Datasets are sorted by the fraction of unique genomic reads. (c-e) Each eCLIP dataset is displayed as a point based on tSNE clustering shown in Figure 2e, with color indicating (c) whether the dataset passed peak-based or family-mapping based quality assessment, (d) the relative information at coding sequence (CDS), or (e) relative information at the 45S ribosomal RNA precursor.

**Extended Data Figure 8.**
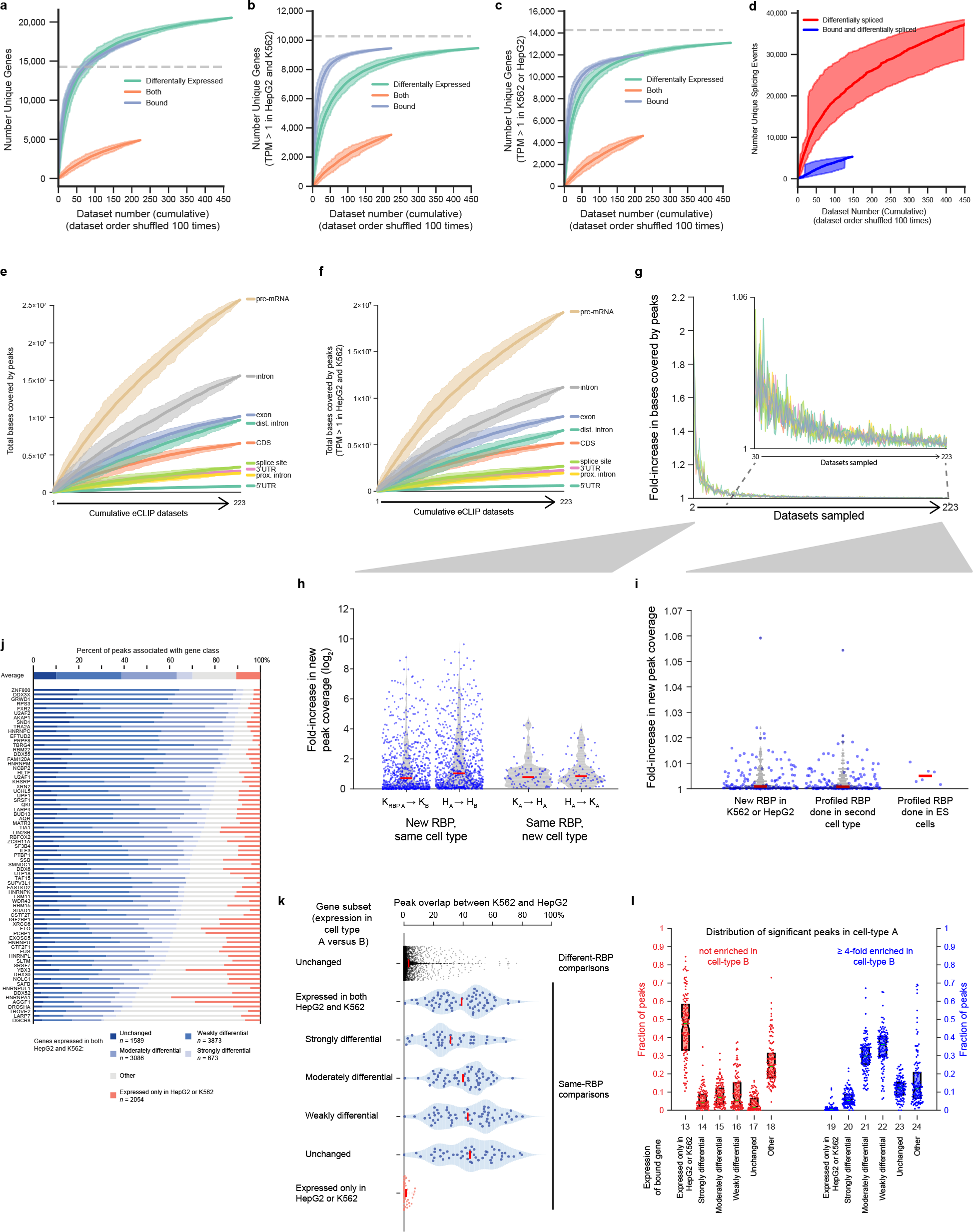
Saturation of RBP binding and regulation in the transcriptome. (a-c) Lines indicate the mean of 100 random orderings of each data type for the number of genes that are (green) differentially expressed upon RBP knockdown and RNA-seq (requiring FDR < 0.05 and *p*-value < 0.05,), (blue) bound in eCLIP (overlapped by a IDR-reproducible peak with *p* ≤ 10-3 and fold-enrichment ≥ 8 in IP versus input), or (orange) both bound and differentially expressed in the same cell type. The set of genes considered was (a) all genes (in GENCODE v19), (b) genes with TPM > 1 in both HepG2 and K562, or (c) TPM > 1 in either K562 or HepG2. Grey dotted line indicates the total number of expressed genes, defined as (a,c) TPM > 1 in either K562 or HepG2 or (b) TPM > 1 in both HepG2 and K562. Shaded regions indicate tenth to ninetieth percentile. (d) Lines indicate the mean of 100 random orderings of datasets for the number of (red) differential splicing events upon RBP knockdown (including cassette exons, alternative 5′ and 3′ splice sites, retained introns, and mutually exclusive exons; requiring FDR < 0.05, *p*-value < 0.05, and absolute value of change in percent spliced in (|∆Ψ|) > 0.05), and (blue) exons both bound by an RBP and differentially spliced upon RBP knockdown in the same cell type (with binding defined as a peak located anywhere between the upstream intron 5′ splice site and downstream intron 3′ splice site). Shaded regions indicate tenth to ninetieth percentile. (e-f) Lines indicate the mean cumulative fraction of bases covered by peaks for 100 random orderings of the 223 eCLIP datasets, separated by transcript regions as indicated, with shaded region indicating tenth and ninetieth percentiles. The set of genes considered was (e) all genes, or (f) only genes with TPM > 1 in both K562 and HepG2. (g) Data and colors as in (e), represented as fold-increase in mean bases covered by peaks from *n* to *n+1* eCLIP datasets. (h) Points indicate the fold-increase in bases covered by peaks between sampling one or two datasets, separated by whether the second is the same RBP in a new cell type (K_A_ -< H_A_ or H_A_ -> K_A_ for RBP A profiled in K562 and then HepG2 or HepG2 and then K562 respectively) or a different RBP in the same cell type (K_A_ -< K_B_ or H_A_ -< H_B_ for RBP A followed by RBP B in either K562 or HepG2 respectively), with kernel smoothed density indicated by the shaded area. Red line indicates median. (i) Points indicate the fold-increase in bases covered by peaks between sampling all versus leaving one dataset out, separated by whether the RBP is (left) a newly profiled RBP or (center) a previously profiled RBP profiled in a second cell type (of either K562 or HepG2). (right) The fold-increase observed if an independent eCLIP experiment performed in H1 or H9 human embryonic stem cells is added (including RBFOX2, IGF2BP3, and two replicates each for IGF2BP1 and IGF2BP2). Red line indicates median. (j) Bars indicate the fraction of peaks observed for each RBP within sets of genes separated by their relative expression change between K562 and HepG2: unchanged (fold-difference ≤ 1.2), weakly (1.2 < fold-difference ≤ 2), moderately (2 < fold-difference ≤ 5) or strongly (fold-difference > 5) differential, or cell-type specific genes (TPM < 0.1 in one cell type and TPM ≥ 1 in the other). *n* indicates the number of genes meeting each criteria. For each RBP, the results shown are for the cell type with fewer total peaks. (k) Points indicate the fraction of overlapping peaks identified from our standard eCLIP processing pipeline between K562 and HepG2 for RBPs profiled (blue or red) in both cell types, or (black) between one RBP in K562 and a second in HepG2, for sets of genes separated by their relative expression change between K562 and HepG2 as in (j). Red line indicates mean. (i) Each point represents one eCLIP dataset compared with the same RBP profiled in the second cell type. For the set of peaks from the first cell type that are not enriched (fold-enrichment < 1) in the second cell type, red points indicate the fraction occurring in genes with the indicated expression difference between HepG2 and K562. Blue points similarly indicate the gene distribution of peaks four-fold enriched in the opposite cell type. Boxes indicate quartiles, with median indicated by the central green line.

**Extended Data Figure 9.**
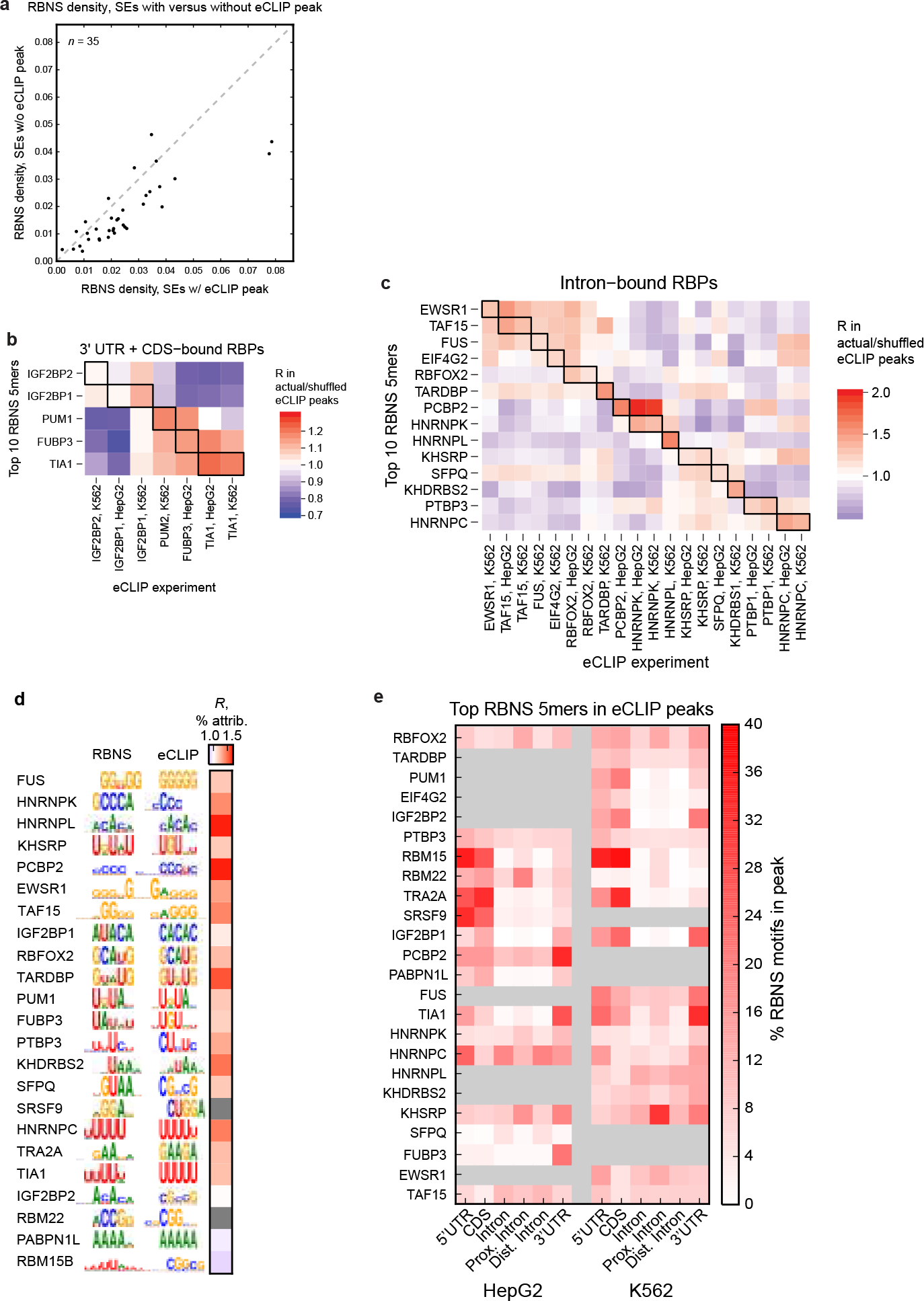
Enrichment of *in vitro* motifs in eCLIP peaks for different RNA types. (a) Comparison of RBNS density (proportion of 5mers that are 10 RBNS 5mers, in the SE and flanking intronic up/downstream 100 nt), for changed SEs with (x-axis) vs. without (y-axis) an eCLIP peak. All experiments with RBNS, eCLIP, and KD/RNA-seq are shown, with 28/35 experiments having greater RBNS density in SEs with an eCLIP peak. (b-c) The average enrichment (geometric mean) of the top 10 RBNS 5mers for a given RBP in the peaks of an eCLIP experiment compared to shuffled eCLIP peaks, among all RBPs predominantly bound to (b) introns or (c) 3’ UTR + CDS by eCLIP. RBPs arranged by RBNS motif similarity along the y-axis, with corresponding RBPs between RBNS and eCLIP boxed along the diagonals. (d) RBP order and RBNS and eCLIP motifs are as in Figure 3a. At right is shown the ratio of the % eCLIP peaks attributable to the top ten RBNS 5mers for each RBP compared to the % of eCLIP peaks attributable to the same ten 5mers, averaged over all other eCLIP experiments in the same RNA type class (from panels b and c above). For 18 out of 21 RBPs the RBNS motifs explain more (R > 1) of the corresponding eCLIP peaks than eCLIP peaks of proteins binding similar transcript regions (SRSF9 and RBM22, shown in gray, were excluded because of insufficient numbers of RBPs in their type class to perform this analysis). (e) The proportion of the top 10 RBNS 5mers that fall within an eCLIP peak, separated by transcript region. RBPs arranged from top to bottom according to the proportion falling within an eCLIP peak over all transcript regions (all motif occurrences in expressed transcripts).

**Extended Data Figure 10.**
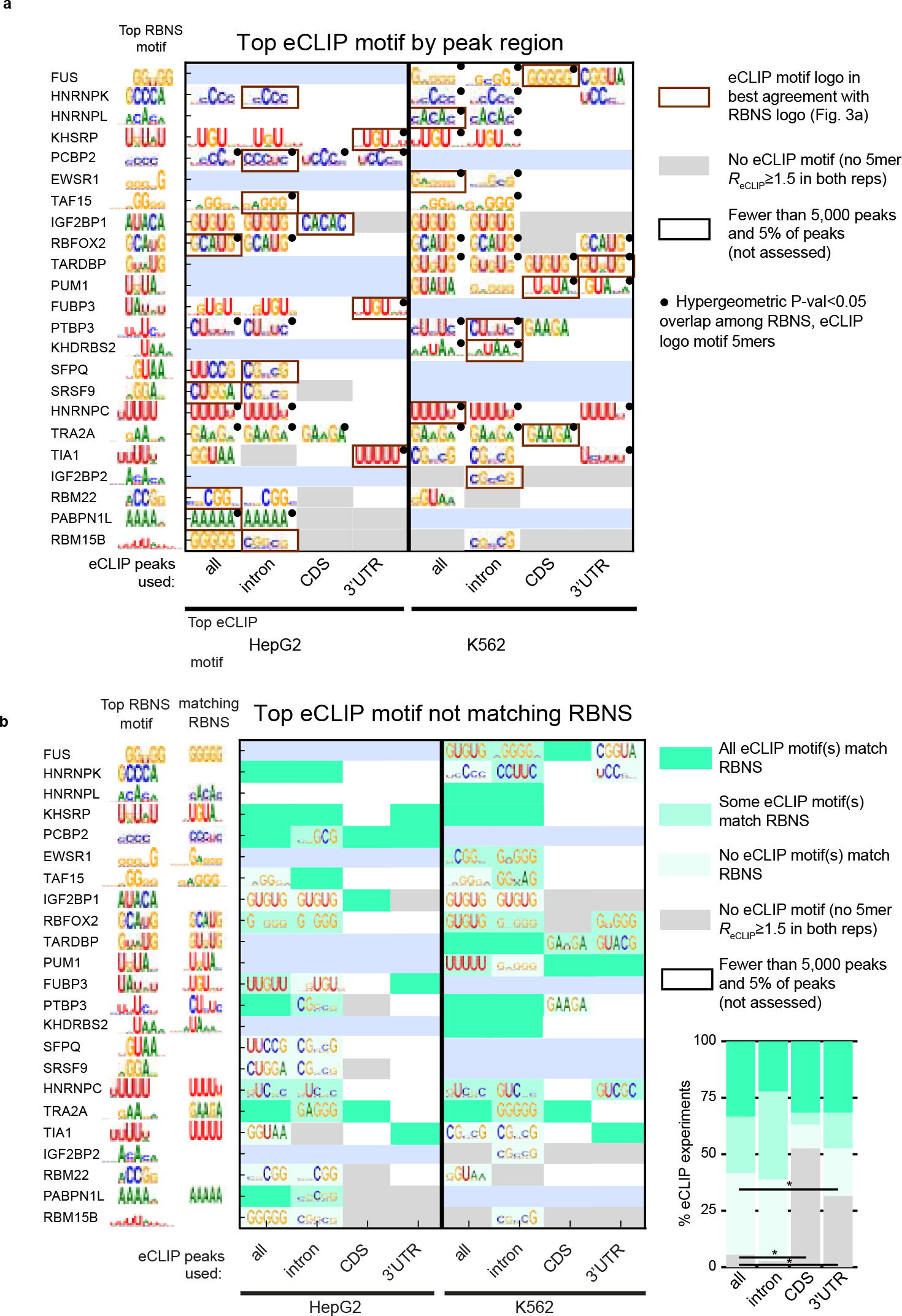
Comparison of *in vitro* RBNS-derived motifs with *in vivo* eCLIP-derived motifs. (a) Top motif derived from all eCLIP peaks as well as eCLIP peaks within intronic, CDS, and 3′UTR regions. Motifs were only derived if there were at least 5,000 peaks or 5% of total peaks in that region, averaged over the two eCLIP replicates. Dashed lines indicate eCLIP was not performed in that cell line. Filled circles indicate significant overlap (P < 0.05 by hypergeometric test) between RBNS and eCLIP motifs. (b) The top eCLIP motif that does not match RBNS for the corresponding RBP (if any). The eCLIP motif was considered as matching RBNS if any of its constituent 5mers were among the RBNS Z≥3 5mers (always using at least 10 RBNS 5mers if there were fewer with Z≥3). Dashed lines indicate eCLIP was not performed in that cell line. (right) The percentage of eCLIP experiments aggregated over all RBP/cell types in each category of agreement with RBNS. Horizontal line indicates a significant difference in the proportion of a particular eCLIP/RBNS agreement category between eCLIP analysis of all peaks versus eCLIP analysis of intron, CDS, or 3′UTR peaks (P < 0.05 by Fisher’s Exact Test).

**Extended Data Figure 11.**
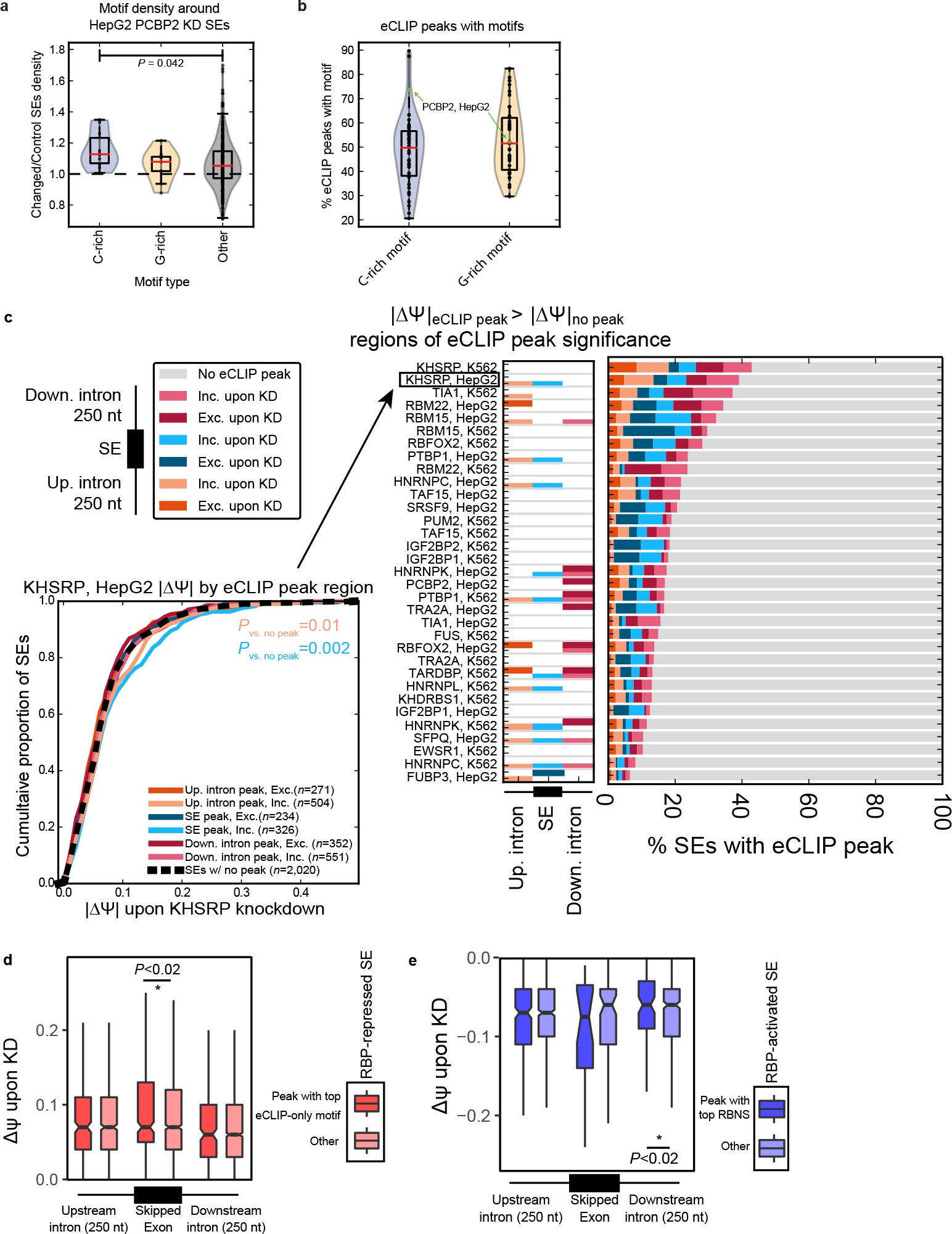
Splicing regulatory activity of RBNS+ and RBNS− eCLIP peaks. (a) Density of 5mers in cassette / skipped exons (SEs) and their flanking intronic up/ downstream 100 nt in changed versus control SEs upon PCBP2 KD in HepG2 cells. The ratio of changed and control frequency was computed for each 5mer with the ratio plotted as density on the y-axis, and 5mers were separated by C-rich (contain 4-5 C’s), G-rich (contain 4-5 G’s), or “Other”. Significance determined by Kolmogorov-Smirnov test. (b) Percentage of eCLIP peaks that contain a C- or G-rich motif (5mer with 4+ of the respective base) among all eCLIP experiments that have corresponding RBNS data. PCBP2 eCLIP in HepG2 cells demarcated (eCLIP with 3rd highest proportion of peaks with C-rich motifs; median for peaks containing G-rich motifs). (c) Left: The distribution of ∆Ψ changes upon KD in each of the 6 eCLIP+ peak region/SE splicing change types compared to that of eCLIP-SEs for KHSRP in HepG2 cells (significant if P<0.05 by Wilcoxon rank-sum test). Center: Regions of significance for eCLIP+ vs. eCLIP-SEs for each eCLIP experiment. Right: Proportion of SEs in each of the six eCLIP+ types for each eCLIP experiment. Bottom: Classification of eCLIP+ peaks into RBNS+ and RBNS− based on the presence of the top RBNS 5mer, shown here for two of the KHSRP peaks in HepG2 cells. (d) Same set of RBPs and corresponding eCLIP+ peak region/SE splicing change types as used in Fig. 3c, but separating eCLIP peaks on whether they contain the top ‘eCLIP-only’ 5mer (based on the motifs from Extended Data Fig. 10b) instead of the top RBNS 5mer. (e) As in Fig. 3c, but shown for RBP-activated SEs (decreased inclusion upon RBP knockdown).

**Extended Data Figure 12.**
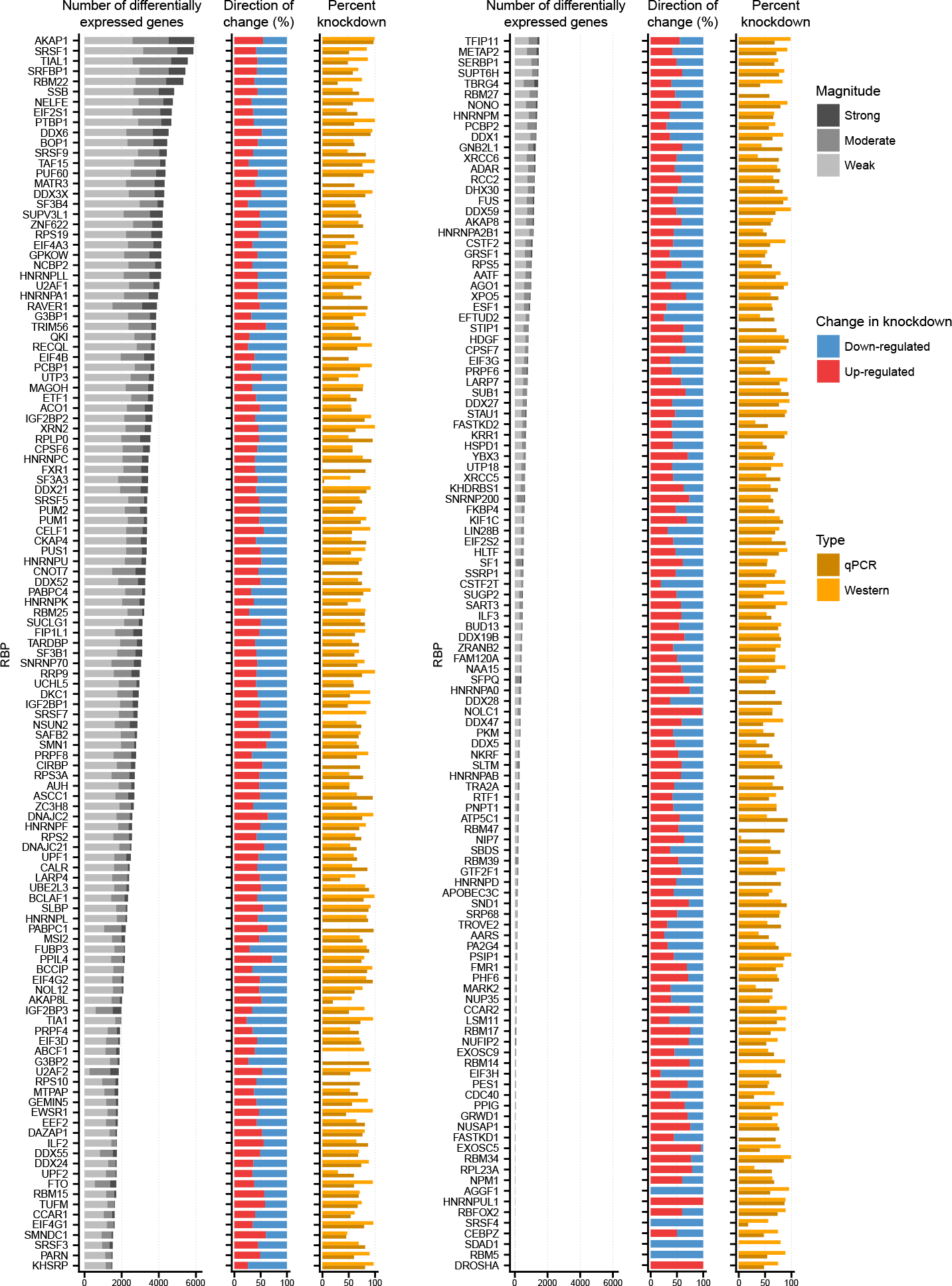
Gene expression changes upon RBP knockdown in HepG2 cells. Each row indicates summary statistics for an RBP knockdown followed by RNA-seq dataset in HepG2 cells. Bars indicate (left) the number and magnitude of differentially expressed genes, (center) the type of regulation and (right) the knockdown level of the targeted RBP protein and/or mRNA. The magnitudes of differential expression were defined as strong (fold-change ≥ 4), Moderate (2 < fold-change < 4) and weak (fold-change ≤ 2). (center) Bars indicate the fraction of differentially expressed genes (red) increased or (blue) decreased upon RBP knockdown. (right) Bars indicate the percent knockdown of the RBP mRNA observed by qPCR and protein observed by Western blot analysis.

**Extended Data Figure 13.**
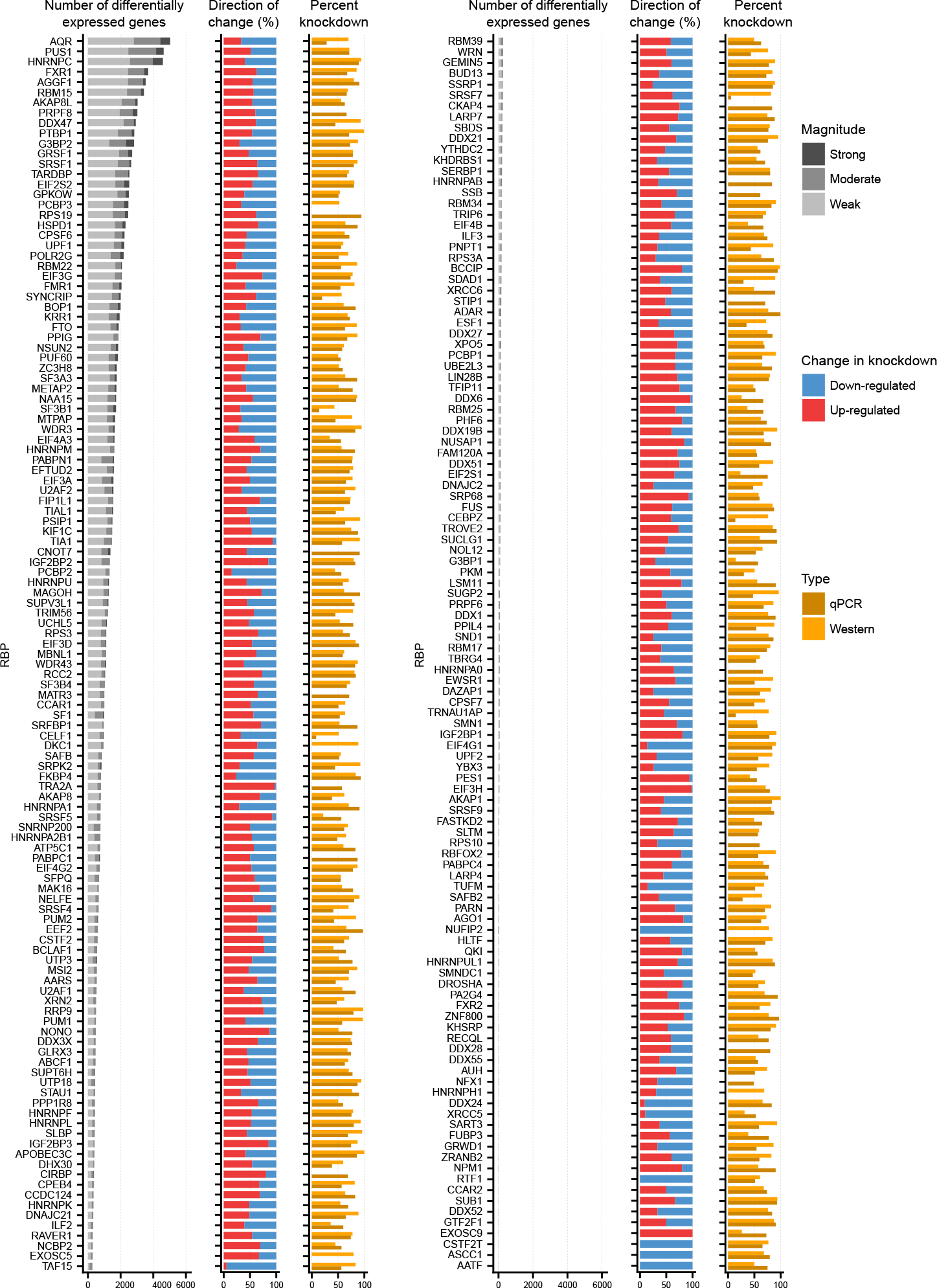
Gene expression changes upon RBP knockdown in K562 cells. Each row indicates summary statistics for an RBP knockdown followed by RNA-seq dataset in K562 cells. Bars indicate (left) the number and magnitude of differentially expressed genes, (center) the type of regulation and (right) the knockdown level of the targeted RBP protein and/ or mRNA. The magnitudes of differential expression were defined as strong (fold-change ≥ 4), Moderate (2 < fold-change < 4) and weak (fold-change ≤ 2). (center) Bars indicate the fraction of differentially expressed genes (red) increased or (blue) decreased upon RBP knockdown. (right) Bars indicate the percent knockdown of the RBP mRNA observed by qPCR and protein observed by Western blot analysis.

**Extended Data Figure 14.**
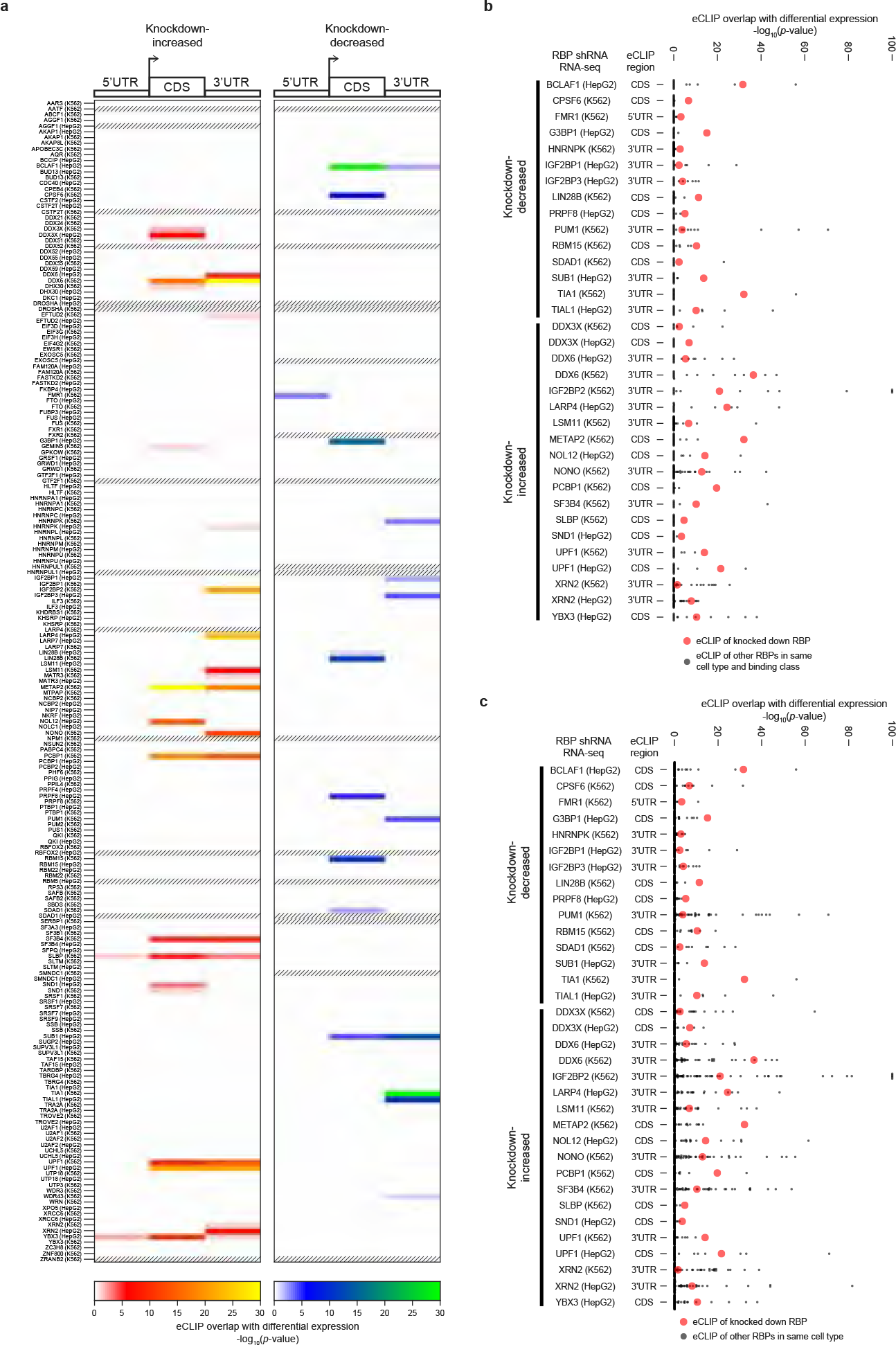
Association between RBP binding and RNA expression upon knockdown. (a) Color indicates the significance of overlap between genes differentially expressed upon knockdown of an RBP and target genes with significant enrichment for 5’UTR, CDS, or 3’UTR regions in eCLIP of the same RBP in the same cell type. Shown are all 203 pairings of knockdown-RNA-seq and eCLIP performed in the same cell type. Dashed boxes indicate comparisons with less than 10 genes altered in RNA-seq. The background gene set for each comparison was chosen by taking genes with at least 10 reads in one of IP or input, and where at least 10 reads would be expected in the comparison dataset given the total number of usable reads. (b-c) Red points indicate significance of overlap between eCLIP and knockdown RNA-seq for the 34 significant overlaps (multiple hypothesis corrected *p*-value ≤ 0.05), showing only the most significantly enriched region from (a). Black points indicate knockdown RNA-seq datasets compared against enrichments for the same transcript region for (b) eCLIP datasets for RBPs within the same binding type class (as identified in Fig. 2b), or (c) all eCLIP datasets in the same cell type.

**Extended Data Figure 15.**
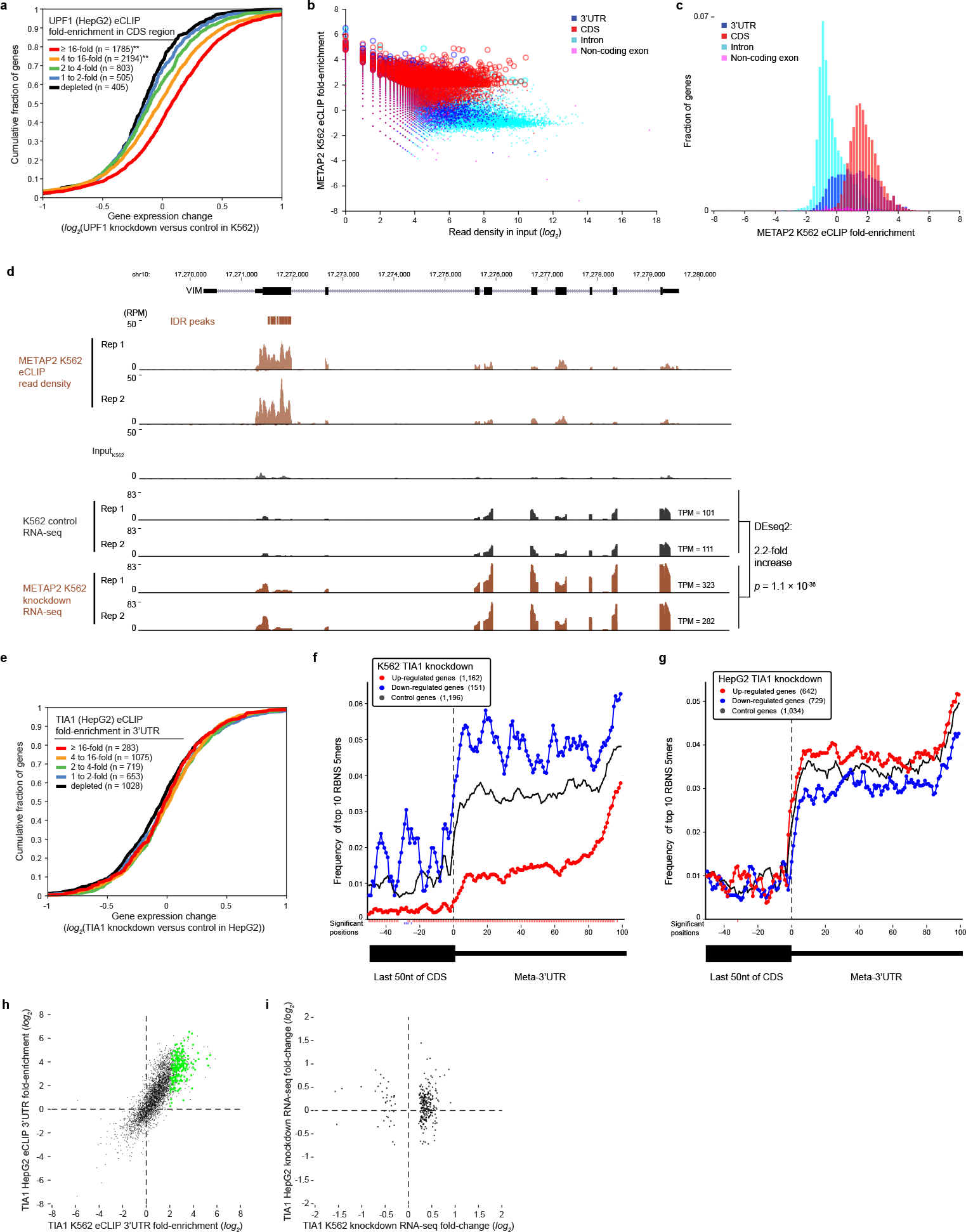
Association between RBP binding and RNA expression upon knockdown. (a) Lines indicate cumulative distribution plots of gene expression fold-change (UPF1 knockdown versus control) for indicated categories of UPF1 eCLIP enrichment in K562 cells. (b-c) METAP2 K562 eCLIP region-level enrichment at 3’UTR, CDS, intronic, and non-coding exonic regions. (b) Points indicate read density in (x-axis) input versus (y-axis) fold-enrichment in METAP2 eCLIP for indicated transcript regions of all GENCODE v19 genes. Significantly enriched regions (p ≤ 10^−5^ and fold-enrichment ≥ 4) are indicated by open circles. (c) Histogram of METAP2 eCLIP fold-enrichment for the indicated transcript regions. (d) Genome browser view of eCLIP and knockdown RNA-seq read density for METAP2 experiments in K562 cells for *VIM*. Read density is shown in reads per million (RPM). (e) Cumulative distribution plots of gene expression fold-change (TIA1 knockdown versus control in HepG2 cells) for indicated categories of 3’UTR TIA eCLIP enrichment. (f-g) Position-specific frequency of the top 10 TIA1 RBNS 5mers in the last 50 positions of the CDS and in a meta-3’UTR of (red) up-regulated, (blue) down-regulated, and (black) control genes upon TIA1 knockdown in (f) K562 and (g) HepG2 cells. Positions of motif density significantly different in up- or down-regulated genes relative to control genes are indicated below the x-axis (calculated using a binomial test comparing the number of regulated genes that do versus do not have one of the top 10 RBNS 5mers at that position versus the frequency observed in control genes). (h) Points indicate fold-enrichment (log_2_) between IP and input for 3’UTR regions of all genes meeting minimal read depth requirements (at least 10 reads in one of IP or input, and where at least 10 reads would be expected in the comparison dataset given the total number of usable reads, were considered) in (x-axis) K562 and (y-axis) HepG2 cells. Points in green indicate genes that had both significant eCLIP enrichment in K562 cells and differential expression upon TIA1 knockdown in K562 cells. (i) Points indicate fold-change (log2) in expression between TIA1 knockdown and control RNA-seq for the set of genes with both significant eCLIP enrichment in K562 cells and differential expression upon TIA1 knockdown in K562 cells.

**Extended Data Figure 16.**
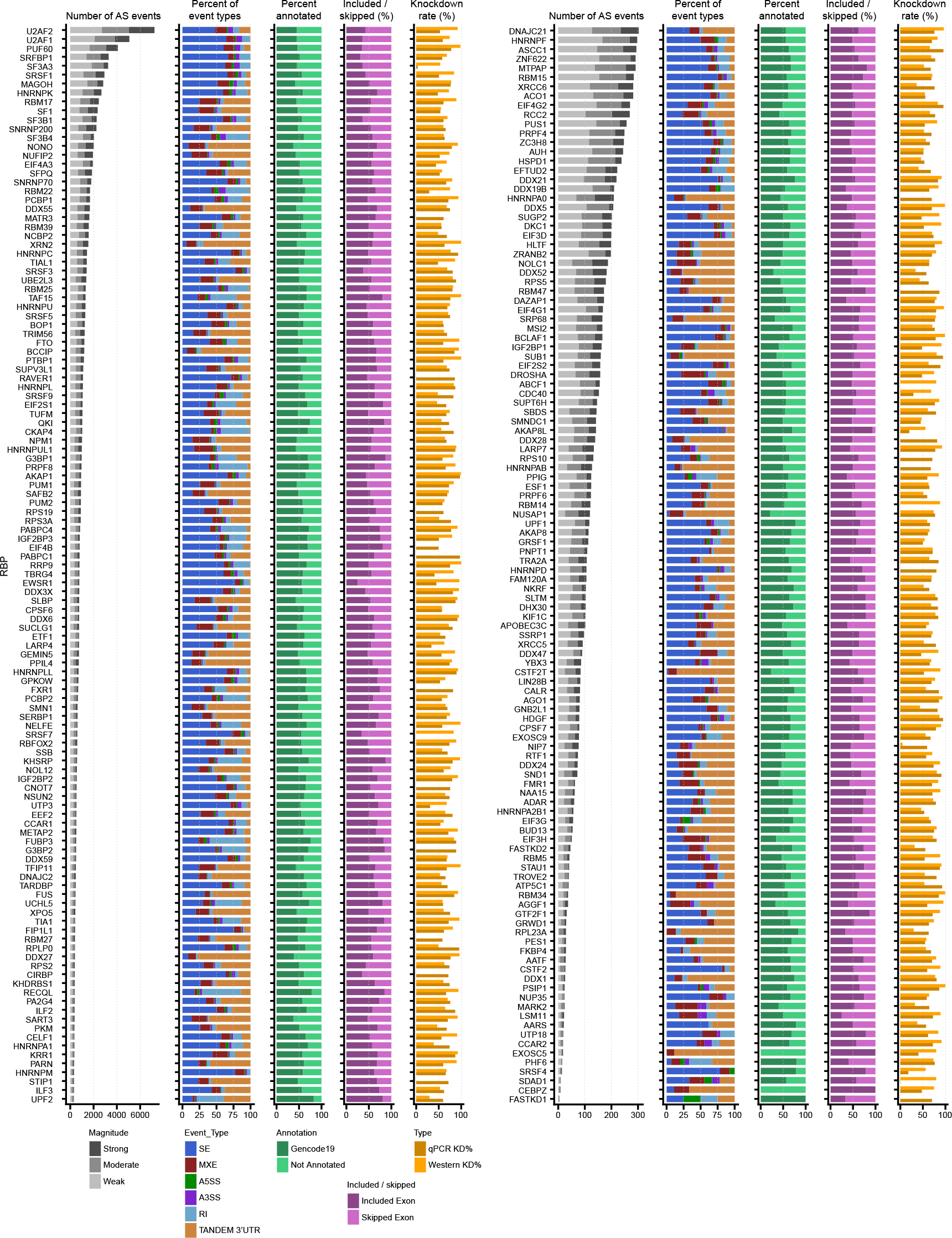
Alternative splicing changes upon RBP knockdown RNA-seq in HepG2 cells. Each row indicates summary alternative splicing statistics for an RBP knockdown followed by RNA-seq dataset in HepG2 cells. Bars indicate (left) the number and magnitude of differentially spliced events, (center-left) the fraction of each type of alternative splicing event, (center) the percent of events observed that are present in GENCODE v19, (center-right) the fraction of cassette exons that are either included or excluded, and (right) the knockdown level of the targeted RBP mRNA and protein by qPCR and Western blot analysis. The magnitudes of differential splicing were defined as weak (|d-PSI| = 5% - 15%), moderate (|d-PSI| = 15% - 30%) or strong (|d-PSI| >= 30. The affected alternative event types are SE (skipped exon), MXE (mutually exclusive exons), A5SS (alternative 5’ splice site), A3SS (alternative 3’ splice site), RI (retained intron) and TANDEMUTR (tandem 3’ UTR).

**Extended Data Figure 17.**
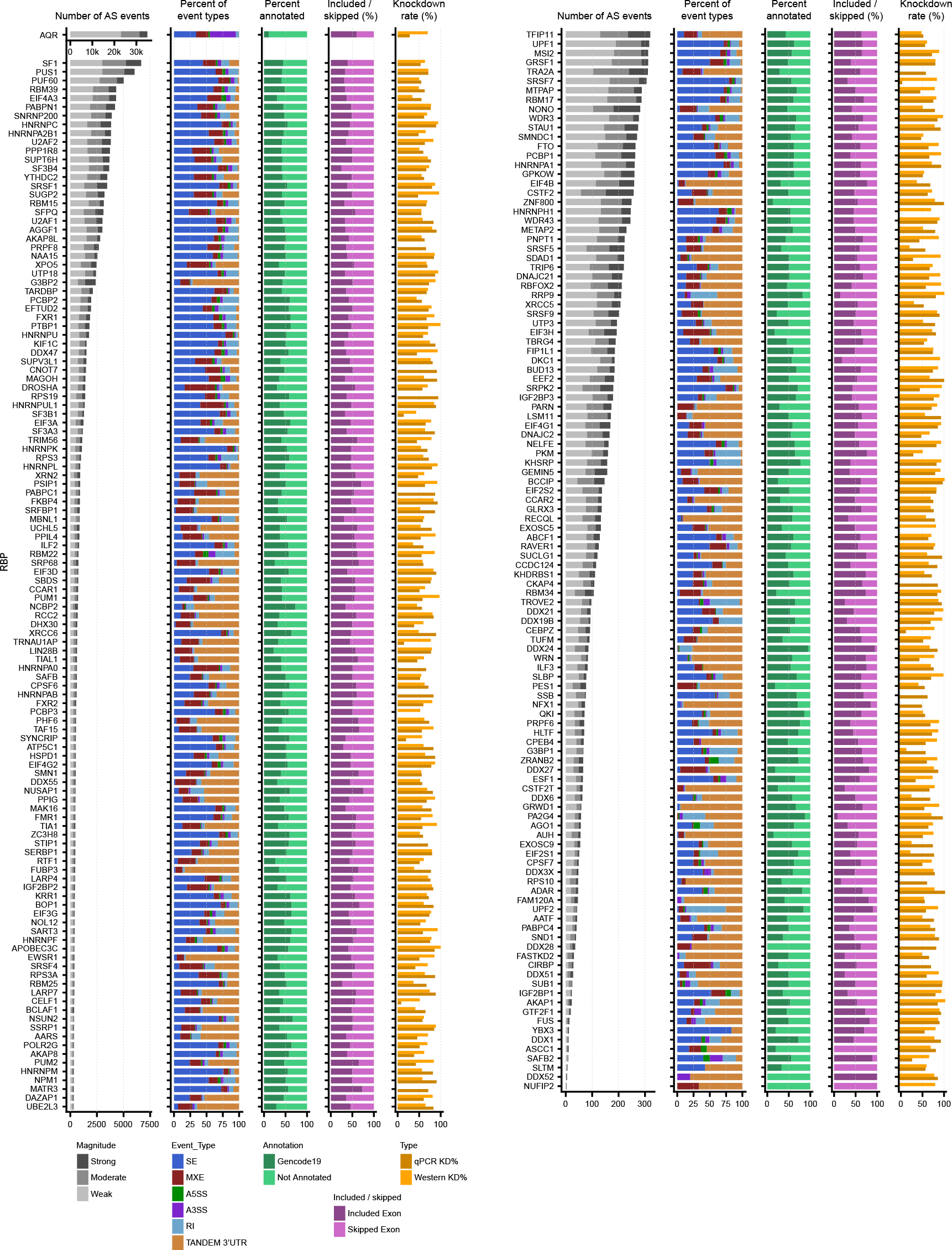
Alternative splicing changes upon RBP knockdown RNA-seq in K562 cells. Each row indicates summary alternative splicing statistics for an RBP knockdown followed by RNA-seq dataset in K562 cells. Bars indicate (left) the number and magnitude of differentially spliced events, (center-left) the fraction of each type of alternative splicing event, (center) the percent of events observed that are present in GENCODE v19, (center-right) the fraction of cassette exons that are either included or excluded, and (right) the knockdown level of the targeted RBP mRNA and protein by qPCR and Western blot analysis. The magnitudes of differential splicing were defined as weak (|d-PSI| = 5% - 15%), moderate (|d-PSI| = 15% - 30%) or strong (|d-PSI| >= 30. The affected alternative event types are SE (skipped exon), MXE (mutually exclusive exons), A5SS (alternative 5’ splice site), A3SS (alternative 3’ splice site), RI (retained intron) and TANDEMUTR (tandem 3’ UTR).

**Extended Data Figure 18.**
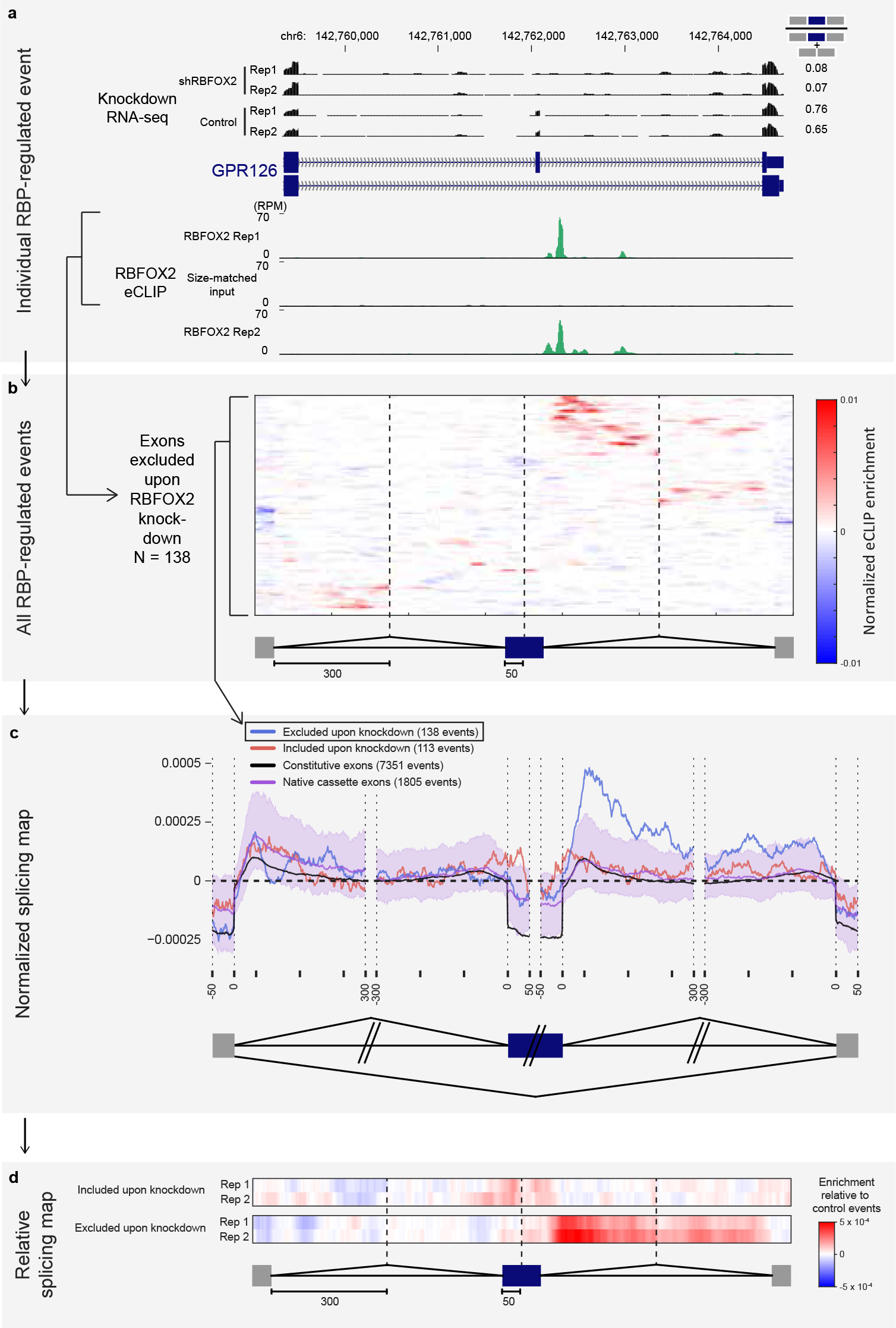
Generation of splicing maps for RBFOX2. (a) First, individual RBP-regulated splicing events are identified from significant changes in knockdown RNA-seq. Genome browser tracks indicate RNA-seq read density (as reads per million (RPM)) and eCLIP read density (RPM) of RBFOX2 in the same cell type, as well as its paired size-matched input. (b) Next, each exon is normalized between IP versus input to obtain ‘Normalized eCLIP enrichment’. The heatmap indicates normalized eCLIP enrichment for all exons significantly excluded upon RBFOX2 knockdown. (c) Next, a ‘splicing map’ is created by calculating the mean and standard error of the mean of normalized eCLIP enrichment for each position across the region, removing the top and bottom 5% outlier values at each position. Lines in splicing map indicate ‘Average eCLIP enrichment’, defined as the mean normalized eCLIP enrichment for exons (red) included or (blue) excluded upon RBFOX2 knockdown. Also plotted are (purple) a control set of cassette exons (referred to as ‘native’ cassette exons) in wild-type HepG2 cells and (black) constitutive exons. Shaded area indicates 0.5^th^ to 99.5^th^ confidence interval obtained by 1000 random samplings of the native cassette exon control set (performed independently using the number of events in either excluded or included sets, and plotting the larger of the two confidence intervals). (d) A final simplified splicing map vector was calculated by subtracting the normalized eCLIP enrichment of control native cassette exons from that of either included or excluded exons at each position to calculate ‘Enrichment relative to control events’.

**Extended Data Figure 19.**
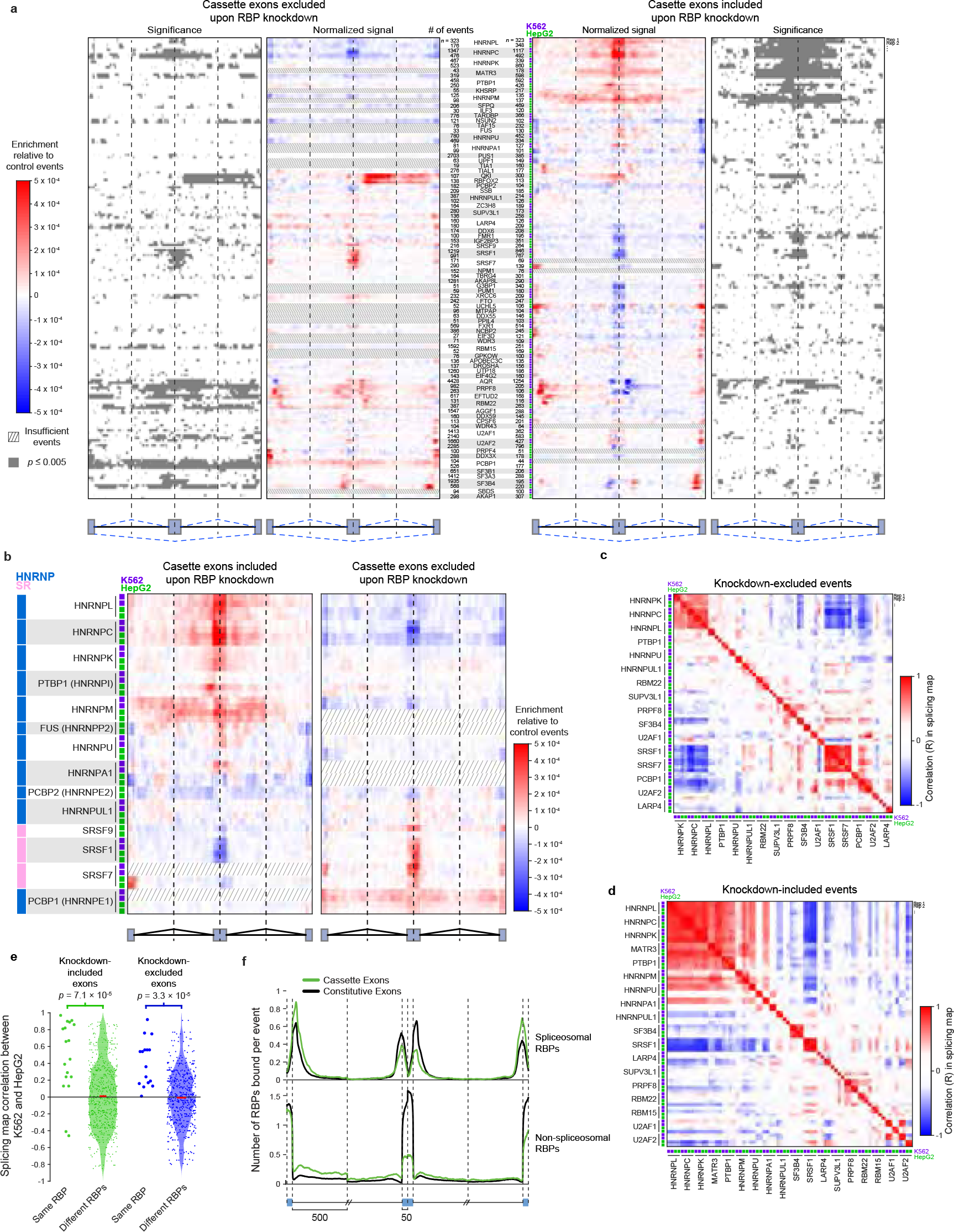
Integration of eCLIP and knockdown RNA-seq to identify splicing regulatory patterns. (a) As in Figure 5b, heatmap indicates the difference between normalized eCLIP read density at cassette exons excluded (left) or included (right) upon RBP knockdown, versus native cassette exons. Out of 203 pairings of eCLIP and knockdown/RNA-seq in the same cell type (139 RBPs total), shown are 92 pairings (72 RBPs) with at least 100 significantly included or excluded events. Outer heatmap indicates positions at which the signal exceeds the 0.5th to 99.5th confidence interval obtained by 1000 random samplings of the same number of events from the native cassette exon control set. Bar graphs indicate the number of RBP knockdown-altered cassette exons for each comparison. Datasets were hierarchically clustered at the RBP-level, and datasets with less than 100 events are indicated by slashed lines. (b) Relative splicing maps for cassette exons included (left) and excluded (right) upon knockdown (as described in Fig. 5b) are shown for all profiled SR and hnRNP proteins. Datasets were hierarchically clustered at the RBP-level, and datasets with less than 100 events are indicated by slashed lines. (c-d) Heatmap indicates correlation (Pearson R) between splicing maps for (c) knockdown excluded or (d) knockdown-included exons for RBPs profiled in both K562 and HepG2 cells, hierarchically clustered at the RBP level. (e) Plot represents the distribution of Pearson correlations between splicing maps as shown in (c-d), separated by whether the comparison is between the same RBP or different RBPs profiled in two different cell types. Different RBPs are shown as smoothed histogram using a Normal kernel, and red line indicates mean. Significance was determined by Kolmogorov-Smirnov test. (f) Lines indicate the average number of RBPs with reproducible eCLIP peaks (out of 223 total datasets) in 50nt exonic and 500nt intronic regions flanking splice sites, separated by whether the RBP is (top) annotated as a spliceosome component or (bottom) all other RBPs.

**Extended Data Figure 20.**
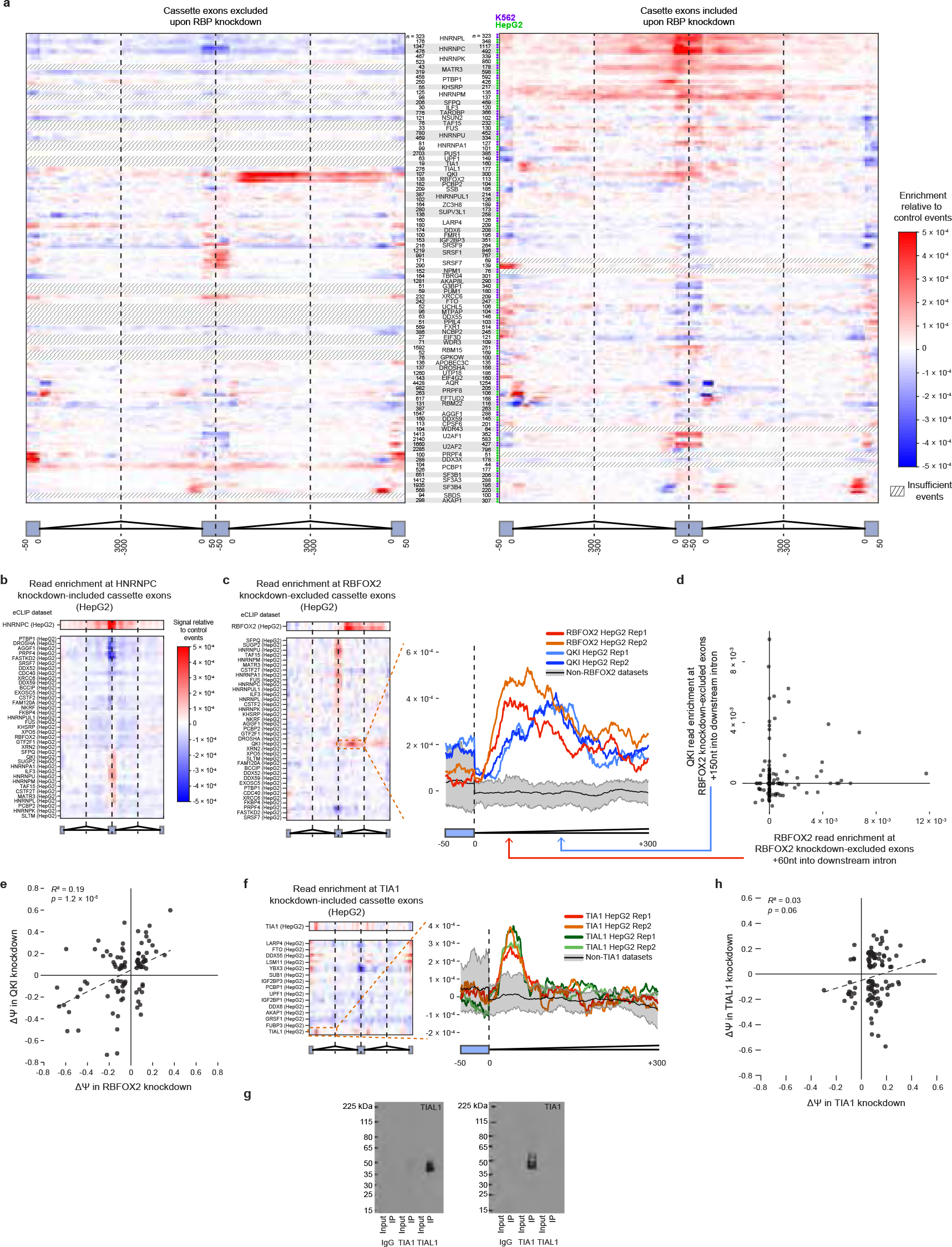
Cross-RBP splicing maps. (a) Similar to Fig. 5b, knockdown-altered cassette exons were identified for each RNA-seq experiment. However, for this analysis normalized eCLIP read density at cassette exons (left) excluded or (right) included upon RBP knockdown versus native cassette exons was calculated separately for all RBPs within the same RBP class (identified in Fig. 2b). The heatmap then indicates the difference between the normalized eCLIP signal for the shRNA-targeted RBP versus the mean of normalized eCLIP signal for all other RBPs within that class. Shown are all 92 pairings of RBPs with eCLIP and knockdown RNA-seq data and at least 100 included or excluded altered events, with dashed lines indicating datasets with less than 100 significantly altered events. (b) Heatmap indicates normalized eCLIP signal at HNRNPC knockdown-induced exons in HepG2 cells relative to native cassette exons for HNRNPC (top) and all other RBPs within the same binding class and cell type (bottom). (c) (left) as in (b) for RBFOX2 knockdown-excluded exons in HepG2 cells. (right) lines indicate normalized signal tracks for eCLIP replicates of RBFOX2 and QKI. Black indicates mean of all non-RBFOX2 datasets, with the 10^th^ to 90^th^ percentile indicated in grey. (d) For each of 138 RBFOX2 knockdown-excluded cassette exons in HepG2 cells, points indicate (x-axis) normalized RBFOX2 eCLIP enrichment at the +60nt position of the downstream intron versus (y-axis) normalized QKI eCLIP enrichment at the +150nt position of the downstream intron (as indicated by arrows in (c)). (e) Points indicate average change in percent exon inclusion (∆Ψ) in two replicates of RBFOX2 knockdown (x-axis) and QKI knockdown (y-axis) in HepG2 cells. Shown are all exons which were significantly altered (*p*-value < 0.05, FDR < 0.1, and |∆Ψ| > 0.05) from rMATS analysis of either RBFOX2 or QKI, and then were further required to have at least 30 inclusion or exclusion reads in both replicates and average |∆Ψ| > 0.05 for both RBFOX2 and QKI knockdown. Significance was determined from correlation in MATLAB. (f) as in (b) for TIA1 knockdown-included exons in HepG2 cells. (g) Western blot for (left) TIAL1 and (right) TIA1 of immunoprecipitation performed with IgG, TIA1 (RN014P, MBLI), and TIAL1 (RN059PW, MBNL) primary antibody. (h) as in (e) for TIA1 and TIAL1 in HepG2 cells.

**Extended Data Figure 21.**
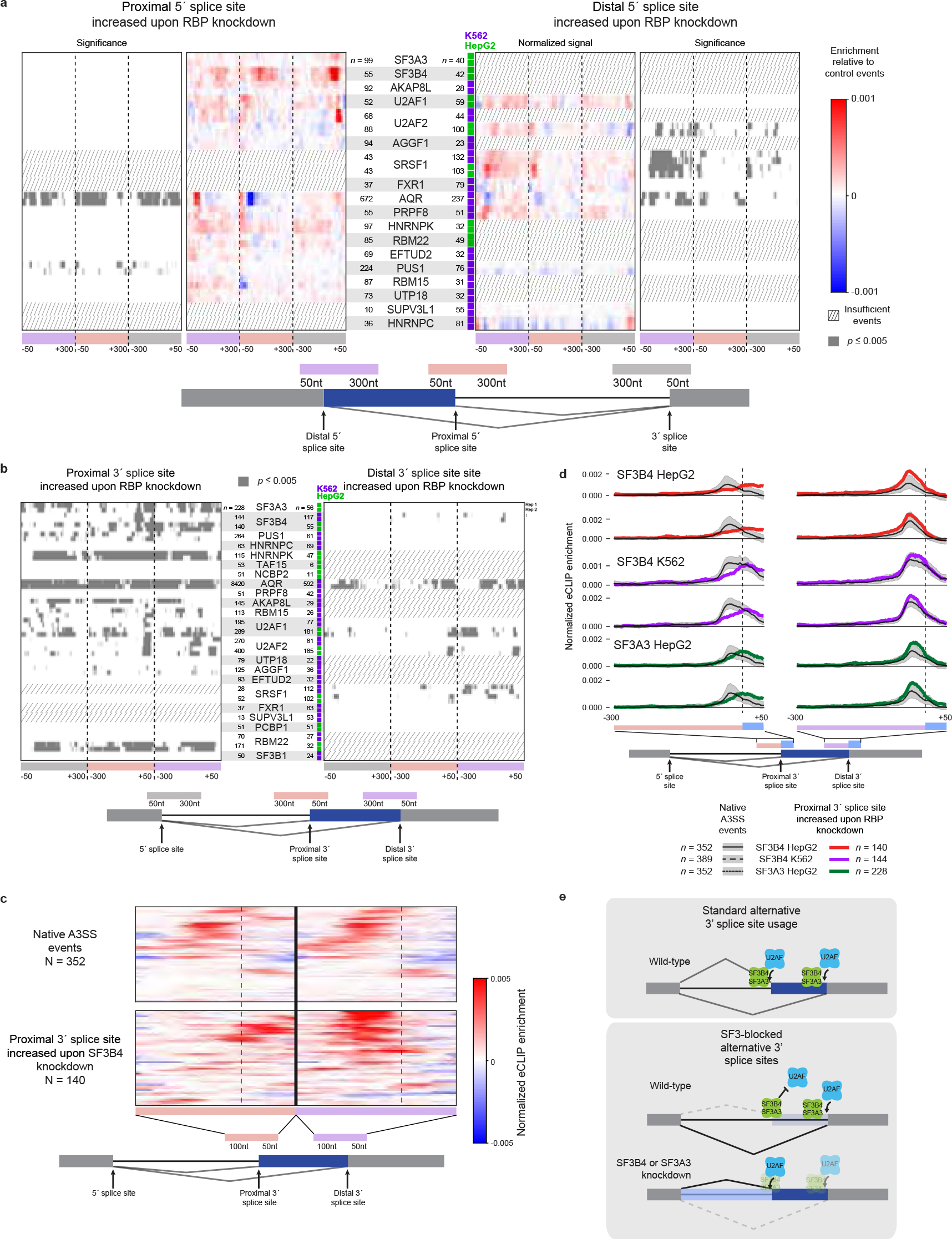
RNA maps for alternative 5’ and 3’ splice sites. (a) Heatmap indicates enrichment at RBP-responsive alternative 5’ splice site events relative to native alternative 5’ splice site events for all RBPs with eCLIP and knockdown RNA-seq data that showed a minimum of 50 significantly changing events upon knockdown. The region shown extends 50 nt into exons and 300 nt into introns. Outer heatmap indicates positions at which the signal exceeds the 0.5^th^ to 99.5^th^ confidence interval obtained by 1000 random samplings of the same number of events from the native alternative 5’ splice site control set. (b) Heatmap indicates positions at which the signal exceeds the 0.5^th^ to 99.5^th^ confidence interval obtained by 1000 random samplings of the same number of events from the native alternative 3’ splice site control set. (c) Heatmap indicates normalized eCLIP signal for SF3B4 in HepG2 cells at alternative 3’ splice site events either (top) alternatively spliced in wild-type cells or (bottom) events with increased usage of the extended 3’ splice site upon SF3B4 knockdown. The region shown extends 50 nt into exons and 100 nt into introns. (d) Lines indicate mean normalized eCLIP enrichment in IP versus input for SF3B4 and SF3A3 at (red/purple/green) alternative 3’ splice site extensions in RBP knockdown or (black) alternative 3’ splice site events in control HepG2 or K562 cells. The region shown extends 50 nt into exons and 100 nt into introns. (e) Model for SF3B4 and SF3A3 blockage of 3’ splice site recognition by U2AF. At SF3-blocked alternative 3’ splice site events, knockdown of SF3 components leads to either usage of the upstream (proximal) 3’ splice site, or retention of the intron.

**Extended Data Figure 22.**
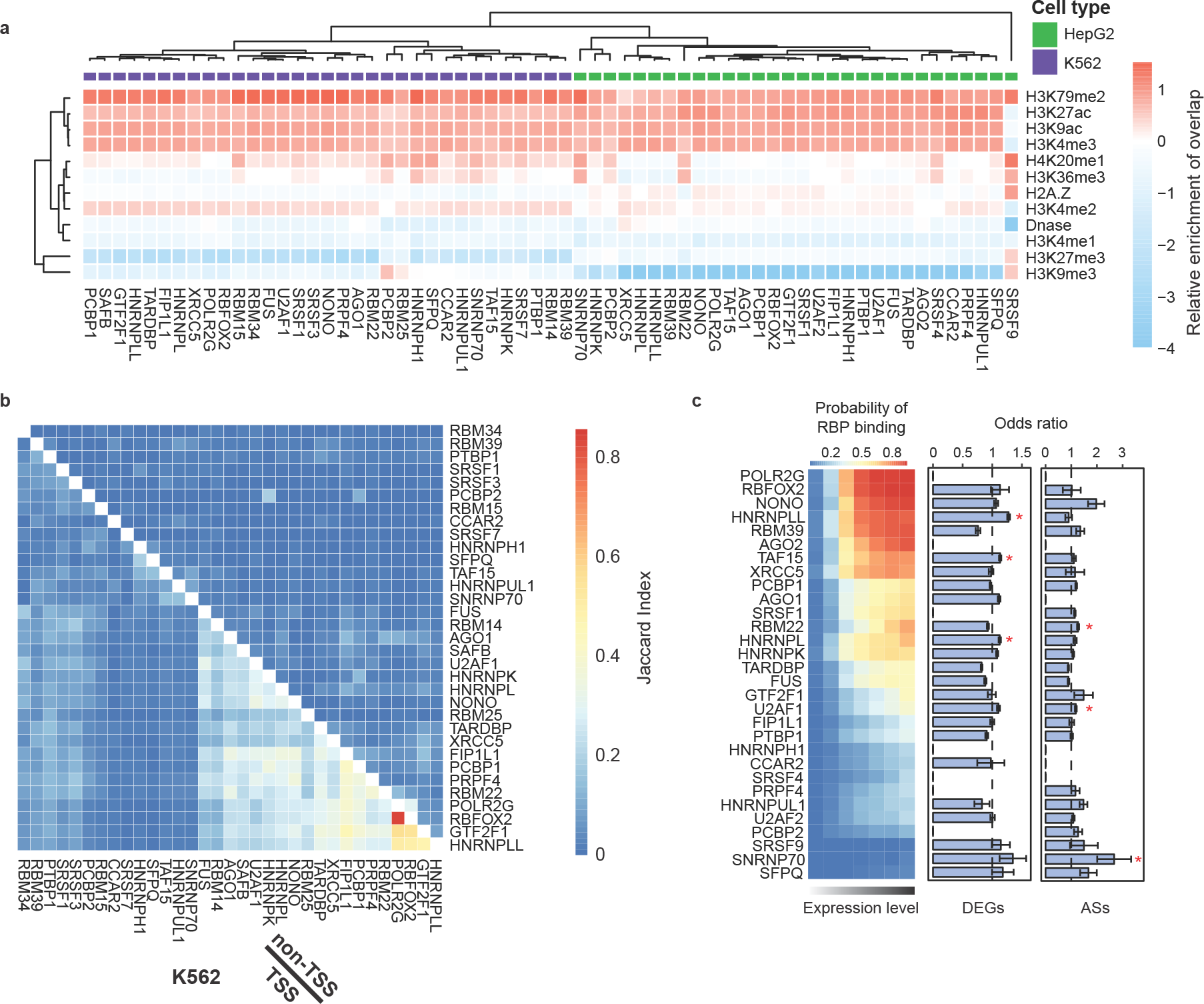
Comparison between RBP DNA and RNA association. (a) Heatmap indicates the relative enrichment of overlap between RBP ChIP-seq peaks and peaks for indicated histone modifications, column-normalized by ‘scale’ in the R heatmap function. (b) Heatmap indicates jaccard indexes between ChIP-seq peaks of different RBPs at promoter regions (bottom left) or non-promoter regions (top right) are displayed as heatmap for K562 cells. (c) (left) Heatmap indicates the fraction of genes (extended 500nt upstream of the TSS and 500nt downstream of the TTS) overlapped by a ChIP-seq peak for each RBP for the set of genes in (x-axis) seven bins of increasing gene expression from RNA-seq in HepG2 cells. (center and right): Bars indicate the odds ratio for overlap between RBP ChIP-seq peak presence and (center) differentially-expressed genes or (right) significant alternative splicing changes upon knockdown of the same RBP. * indicates p-value<0.05 as determined by 100 random samplings of genes with similar expression levels.

**Extended Data Figure 23.**
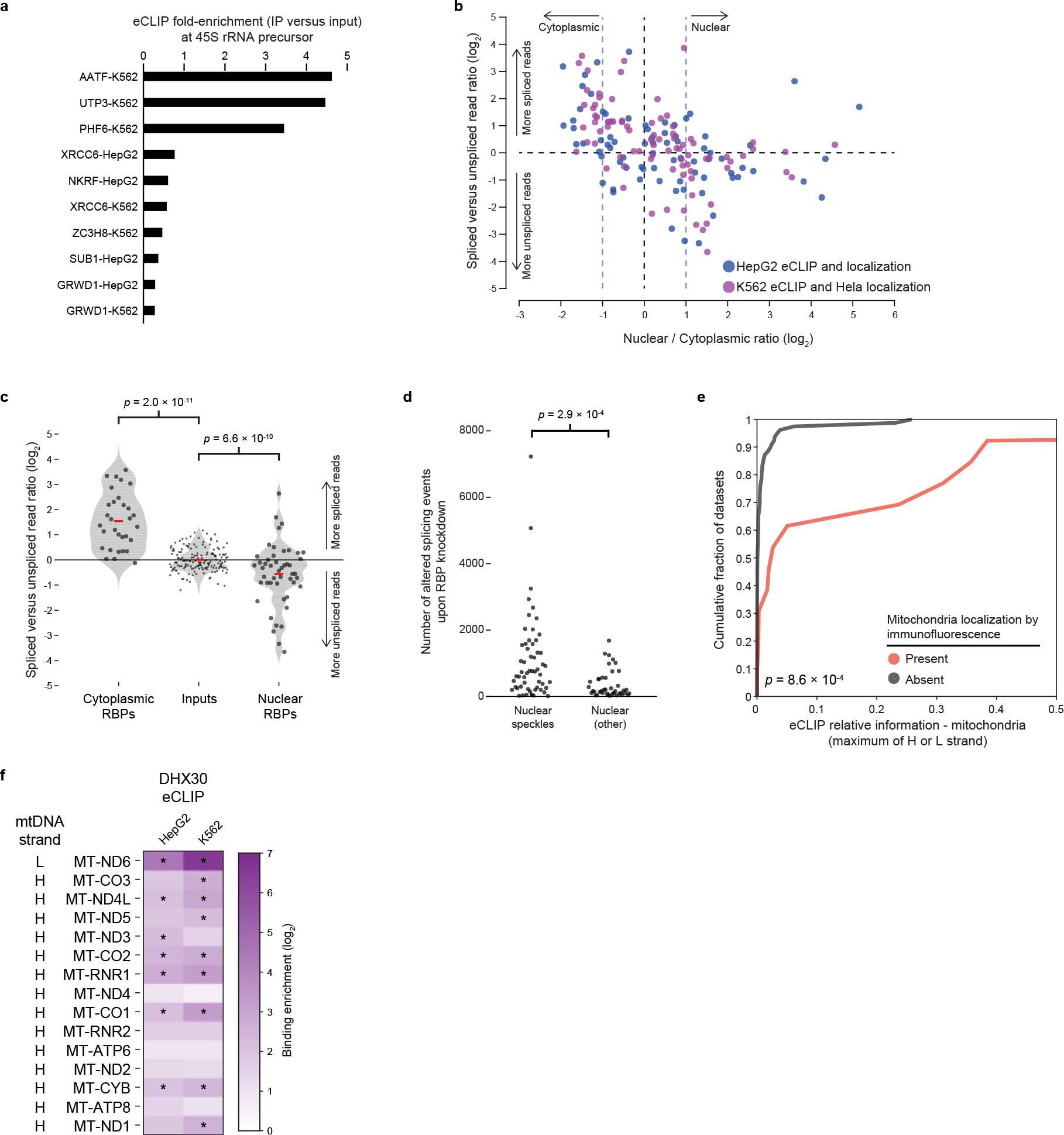
eCLIP binding patterns in subcellular space. (a) Bars indicate fold-enrichment for the 45S ribosomal RNA precursor observed for 8 RBPs with eCLIP data, nucleolar localization observed in immunofluorescence imaging, and no human RNA processing function identified in literature searches. (b) Points indicate (x-axis) nuclear versus cytoplasmic ratio from immunoflourescence (IF) imaging versus (y-axis) ratio of spliced versus unspliced exon junction reads, normalized to paired input. RBPs profiled by eCLIP and IF in HepG2 cells are indicated in blue, and RBPs profiled by eCLIP in K562 cells (in purple) were paired with IF experiments performed in Hela cells. eCLIP data shown is from replicate 1. (c) Points indicate values as in (a), with RBPs separated into nuclear (nuclear / cytoplasmic ratio ≥ 2) and cytoplasmic (nuclear / cytoplasmic ratio ≤ 0.5). Significance was determined by Kolmogorov-Smirnov test, and red line indicates mean. eCLIP data shown is from replicate 1. (d) Points indicate the number of differential splicing events observed upon knockdown of each RBP, separated by the presence or absence of localization in (left) nuclear speckles or (right) nuclear but not nuclear speckles. Significance was determined by Kolmogorov-Smirnov test. (e) Cumulative distribution curves indicate total relative information content for the mitochondrial genome for RBPs with mitochondrial localization by IF (red) and all other RBPs (grey). Significance was determined by Kolmogorov-Smirnov test. (f) Heatmap indicates DHX30 eCLIP enrichment across all exons for all mitochondrial protein coding and rRNA transcripts.* indicates significant eCLIP signal (fold-enrichment ≥ 4 and *p* ≤ 0.00001 in IP versus input).

## Table legends

**Supplementary Data 1. Manual annotation of RBP functions.**

**Supplementary Data 2. ENCODE accession identifiers of datasets used.**

**Supplementary Data 3. RBP gene expression in ENCODE cell lines and tissues.**

**Supplementary Data 4. Summary information for eCLIP experiments.**

**Supplementary Data 5. Summary information for RNA-seq experiments.**

**Supplementary Data 6. Summary information for RBNS experiments.**

**Supplementary Data 7. Summary information for ChIP-seq experiments.**

**Supplementary Data 8. Automated and manual quality assessment of eCLIP datasets.**

**Supplementary Data 9. Summary information for questionable quality eCLIP experiments.**

**Supplementary Data 10. Summary information for eCLIP experiments failing quality assessment.**

**Supplementary Data 11. eCLIP blacklist regions.**

**Supplementary Data 12. Overlap between eCLIP and ChIP-seq peaks.**

**Supplementary Data 13. eCLIP adapters used.**

